# Pretrained gene representations transfer mean expression more broadly than spatial patterns in virtual spatial transcriptomics

**DOI:** 10.64898/2026.09.15.751768

**Authors:** Tingjun Chen, Stephanie C. Hicks

## Abstract

Virtual spatial transcriptomics models combine tissue images with pretrained gene representations to predict expression for unseen genes, but what these representations contribute is unclear. We separated their effects into overall gene-mean and spatial variation using held-out genes and individuals across four cohorts spanning three human brain regions and HER2-positive breast cancer. With fixed representations derived from regulatory DNA sequences and representations pretrained on single-cell transcriptomes, reductions in gene-mean error accounted for 91–96% of the total mean squared error reductions over matched random vectors. Mean-only models using gene representations without images retained 82–99% of the corresponding full-matrix correlation gain. Spatial gains were smaller, increased with expression variation in training tissue, and varied across cohorts and representations. Thus, pretrained gene representations broadly transfer mean expression but only selectively support spatial recovery, showing that gene-panel extension alone does not establish spatial recovery for new targets.

---

Spatial transcriptomics links gene expression to tissue structure, but collecting paired molecular measurements and tissue images remains costly at scale [1, 2]. Virtual spatial transcriptomics aims to extend spatial profiling to new samples by predicting gene expression from tissue images [3–6]. Most predictors are trained for a predefined set of genes, although recent large-scale models support prediction across a broader range of tissues and larger sets of genes [7–9]. A further goal is to predict genes not used to train the spatial predictor, with the long-term aim of generalizing across all genes and biological samples.

*Gene-query* models are designed to address this challenge by taking a representation of the target gene alongside the tissue image [10–14]. This representation, often called a gene embedding, is a numerical vector derived from molecular data or gene annotations. A shared predictor uses the image and gene vector to estimate expression at each location, allowing it to apply information learned from training genes to other genes. GeneQuery, for example, represents genes through text descriptions of their functions and predicts their expression from histology images [11]. DeepSpot-M uses gene representations from pretrained models of DNA, RNA, proteins, single cells and biomedical text to predict spatial expression from histology [13].

The information available to these predictors depends on how the gene representations were learned. Decima is a model trained to predict expression across cell types and conditions from the DNA sequence surrounding a gene [15]. scGPT learns representations of genes through pretraining on single-cell transcriptomes [16]. Vectors extracted from either model can supply gene information to a separately trained spatial predictor. Such information could help estimate a gene’s mean expression across tissue locations, recover its pattern of variation across those locations, or both.

An overall prediction score can obscure this distinction. Correlations and errors computed over the complete location-by-gene matrix combine differences in mean expression between genes with variation across locations within each gene. For example, a model that assigns one accurately predicted mean to every location can therefore improve overall agreement without recovering a gene-specific spatial pattern. Existing approaches assess spatial agreement using correlations for individual genes or measures of spatial structure. Others train models to predict relative expression, expression differences or deviations from gene means [14, 17–20]. Independent benchmarking has also shown that conclusions depend on the metric and evaluation setting [21]. A complementary question is how much pretrained gene representations improve mean-expression estimates and within-gene spatial prediction relative to matched controls, and which genes benefit when excluded from spatial-model fitting.

Here we evaluate gene-mean prediction and within-gene spatial recovery for held-out genes in new biological individuals across four cohorts spanning three human brain regions and HER2-positive breast cancer. We use fixed Decima and scGPT representations, matched gene-vector controls and independently fitted no-image models to assess what pretrained gene information contributes to prediction. We find broad gains in gene-mean prediction but smaller average spatial gains, which increase with expression variation in training tissue and differ across cohorts and representations. We also assess these two components in retrained GeneQuery models and a released DeepSpot-M model under their respective training settings.

## 1. Results

### 1.1 Separating mean-expression prediction from spatial recovery

For gene *g* at spot *i*, we decomposed normalized log-expression as *Y*_*ig*_ = *µ*_*g*_ + *δ*_*ig*_, where *µ*_*g*_ is the mean across evaluated spots and *δ*_*ig*_ is the deviation from that mean (**Fig. 1a**). Applying the same decomposition to predicted gene expression *Ŷ*_*ig*_ gives an exact partition of mean squared error (MSE):

**Figure 1.**
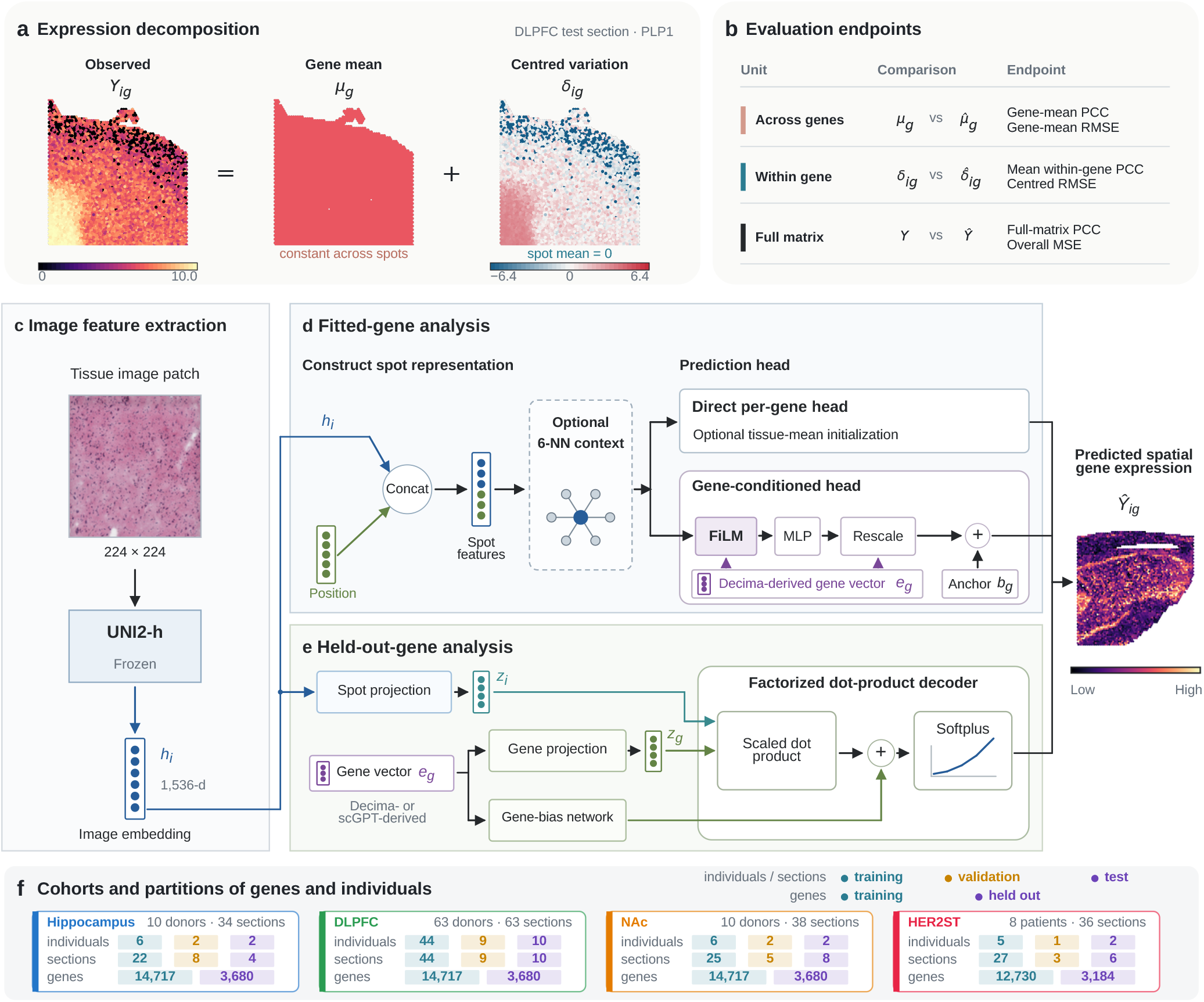
Separating mean expression and spatial variation in fitted and held-out genes. **a**, DLPFC *PLP1* expression decomposed into an across-spot gene mean and centred within-gene variation. **b**, Metrics for gene means, within-gene variation and the complete matrix. **c**, Frozen UNI2-h encodes each 224 *×* 224 tissue patch as a 1,536-dimensional spot vector. **d**, Fitted-gene models combine this vector with coordinate descriptors, with optional six-nearest-neighbour (6-NN) aggregation. Predictions use either a direct per-gene head, optionally initialized with training-tissue means, or a Decima-conditioned feature-wise linear modulation (FiLM) model followed by gene-dependent rescaling and addition of a training-tissue- or sequence-derived anchor. **e**, The held-out-gene model combines separate spot and gene encoders through a scaled dot product and gene-bias network, followed by softplus. Gene vectors come from frozen Decima or scGPT representations; coordinates and graph inputs are omitted. The shared output illustration represents predictions from either model family. **f**, Training, validation and test partitions for individuals and sections, and training and held-out gene counts, in each cohort.

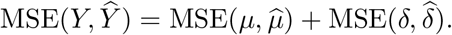

The identity holds because every gene is evaluated over the same spots and the centred errors sum to zero within each gene. It assigns any MSE improvement to better prediction of gene means, within-gene variation or both.

We assessed mean-expression prediction using Pearson correlation across genes (gene-mean PCC) and the root mean squared error (RMSE) of gene means. To assess spatial prediction, we correlated observed and predicted expression across spots for each gene, then averaged these correlations over genes (mean within-gene PCC). Because pooled correlations can include differences between sections, we also evaluated spatial agreement using pooled section-centred residuals and correlations calculated separately within sections (Methods; **Fig. 1b**). Centred RMSE measured errors in within-gene variation, including its magnitude. Full-matrix PCC and overall MSE assessed the complete expression matrix.

We used two separately trained model families (**Fig. 1c–e**). The held-out-gene analysis excluded evaluation genes from fitting and selection of a shared expression predictor (decoder), and tested them in new donors or patients. The fitted-gene analysis used genes included during optimization to establish how changes to the prediction model affect the two components. The held-out-gene decoder combined frozen UNI2-h image features with fixed Decima or scGPT gene vectors, without coordinates or graph inputs.

The analysis included 171 sections: 34 hippocampal sections from ten donors, 63 dorsolateral prefrontal cortex (DLPFC) sections from 63 donors, 38 nucleus accumbens (NAc) sections from ten donors and 36 HER2ST sections from eight patients (**Fig. 1f**). Each cohort was modelled separately. The primary gene split reserved 3,680 of 18,397 genes in each brain cohort and 3,184 of 15,914 genes in HER2ST for evaluation.

### 1.2 Mean initialization and neighbourhood context improve different aspects of prediction

For genes included in model training, mean-expression initialization and neighbourhood information had different effects on prediction (**Fig. 2**; **Extended Data Fig. 1**). Initializing the gene-specific output layer with mean expression from the training samples increased full-matrix PCC by 0.022–0.071 across cohorts, whereas pooled-spot mean within-gene PCC changed by only *−*0.009 to 0.002. Combining information from the six nearest spots and encouraging neighbouring predictions to vary smoothly increased within-gene PCC by 0.0029–0.0110 on this background, with larger gains for high-variance genes (**Extended Data Fig. 2**). A predictor that used Decima vectors to rescale and shift shared spot features through feature-wise linear modulation (FiLM) did not consistently improve within-gene PCC over the direct predictor; effects varied by cohort and model configuration. Illustrative maps showed lower correlation for hippocampal *SLC17A7* and higher correlation for HER2ST *A2M*. Full comparisons are provided in Supplementary Note 1 and Supplementary Tables S2 and S20.

**Figure 2.**
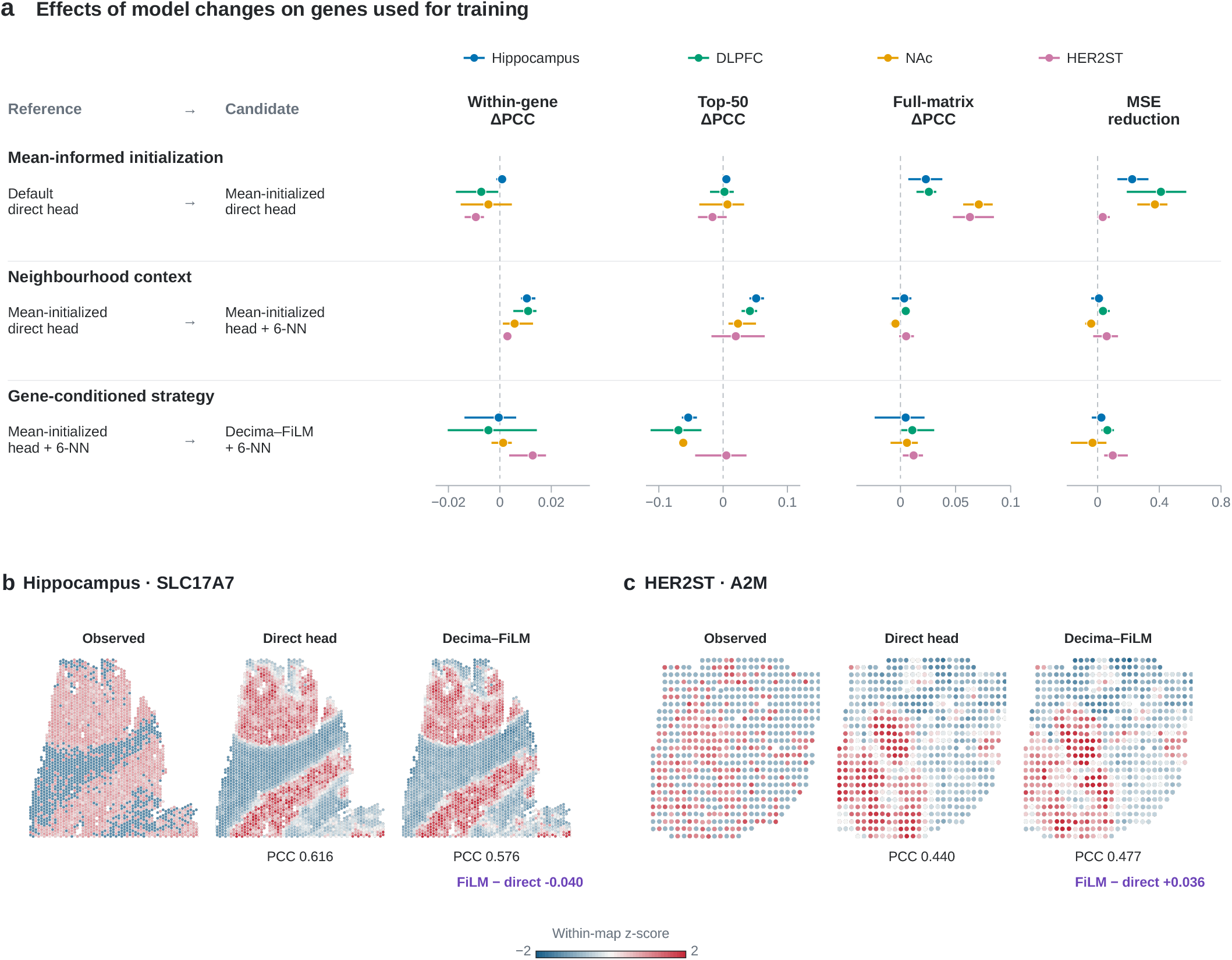
Mean initialization and neighbourhood context improve different components of prediction. **a**, Three representative comparisons: mean-informed direct-head initialization (variant 3 versus 1), neighbourhood context on the mean-initialized background (4 versus 3), and Decima–FiLM on the neighbourhood-enabled background (8 versus 4). **Extended Data Fig. 1** gives all eight variants and matched comparisons. Mean-informed initialization sets trainable output biases to training-tissue gene means and initializes output weights to zero; the 6-NN intervention combines aggregation and Laplacian smoothing. Correlation effects are candidate minus reference; MSE reductions are reference minus candidate. Positive values favour the candidate. Points show means over three independently optimized runs; lines span the minimum and maximum. **b,c**, Hippocampal *SLC17A7* in section GSM8226200_V10B01-086_C1 (**b**) and HER2ST *A2M* in section H2 (**c**), selected as described in Supplementary Methods. Direct and FiLM predictions use variants 4 and 8. Predictions were averaged spotwise across runs before section-level PCC was calculated. Maps were standardized separately within gene, section and condition and clipped to [*−*2, 2] for display. Printed PCCs and differences use unstandardized expression.

### 1.3 Pretrained gene representations transfer mean expression broadly to held-out genes

We trained matched held-out-gene decoders with Decima vectors, dimension-matched random vectors that were unique to each gene but carried no pretrained gene information, or one constant vector shared by all genes. The constant-vector model could vary predictions across tissue locations through the image input, but could not distinguish among target genes. The primary gene split was fixed using hippocampal training individuals before decoder development. Evaluation genes were excluded from fitting and checkpoint selection in every cohort.

Decima produced large matrix-level gains (**Fig. 3a**). Relative to random vectors, full-matrix PCC increased by 0.320–0.418 and MSE decreased by 0.450–0.725. Decima also outperformed the constant-vector control in full-matrix PCC and MSE in every cohort and run.

**Figure 3.**
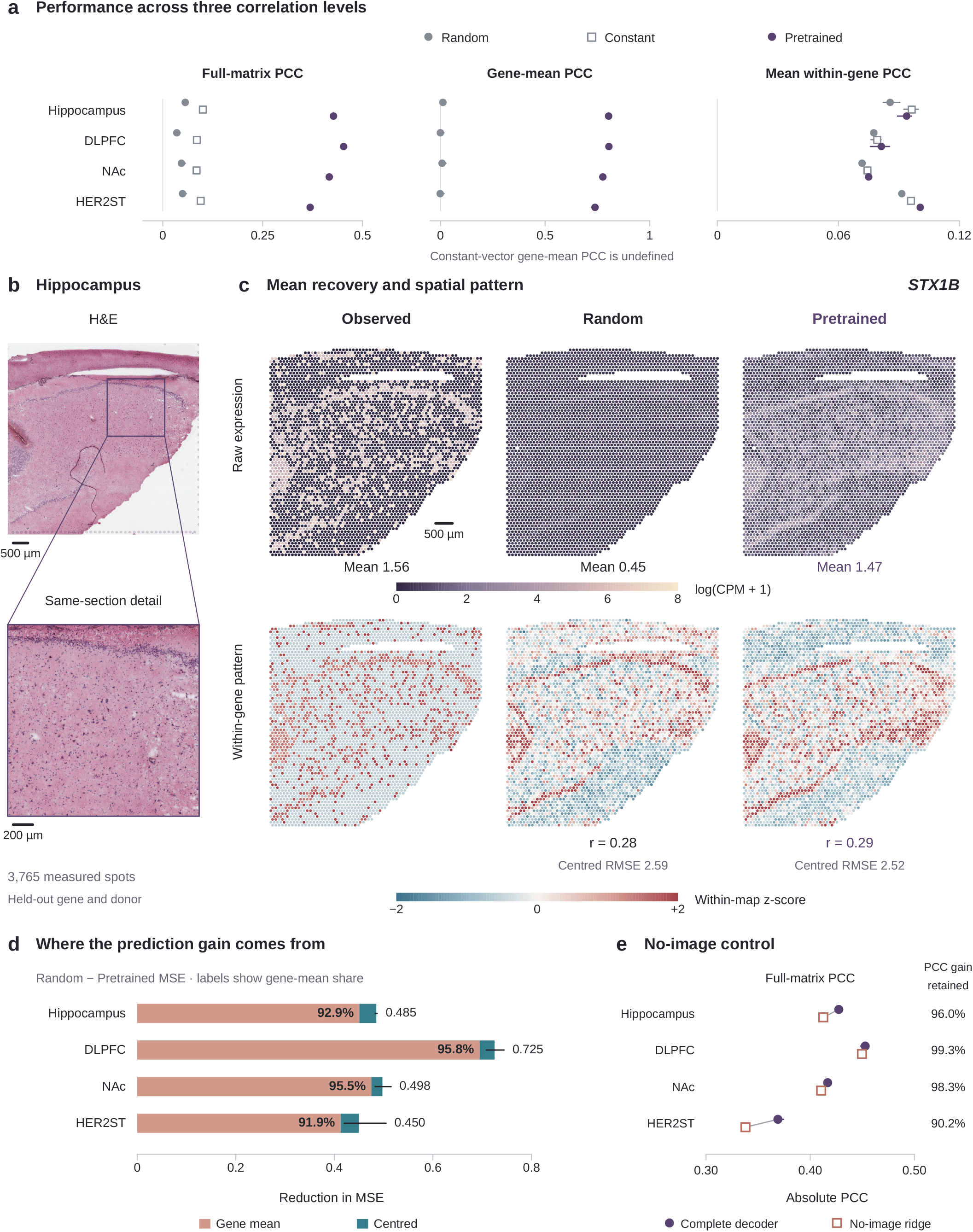
Pretrained gene representations improve gene-mean prediction for held-out genes. **a**,Full-matrix, gene-mean and mean within-gene PCC for random, constant and pretrained vectors in four cohorts. “Pretrained” denotes Decima-derived vectors. Symbols show means over three independently optimized runs; lines span their ranges. Constant-vector gene-mean PCC is undefined. **b**, H&E overview and detail of the hippocampal *STX1B* test section, containing 3,765 spots. The box marks the enlarged field. Scale bars, 500 *µ*m (overview) and 200 *µ*m (detail). **c**, Observed *STX1B* expression and predictions from random- and Decima-vector decoders. Raw maps share a linear log(CPM + 1) colour scale from 0 to 8. Within-gene maps were standardized separately and clipped to *±*2 standard deviations. Means, PCC and centred RMSE use unstandardized expression. Maps share the overview’s field and orientation; scale bar, 500 *µ*m. Circles mark measured spot centres and are enlarged for legibility; no interpolation or smoothing was applied. The section and seed-42 predictions were selected as described in Supplementary Methods. **d**, Random-minus-Decima MSE reduction partitioned into gene-mean (terracotta) and centred (teal) components. Bars show mean component reductions across runs; black lines span total reductions. Internal labels show the mean paired-run gene-mean share; right-hand labels show mean total reductions. **e**, Absolute full-matrix PCC for the complete decoder and an independently fitted no-image ridge model predicting one mean per gene. Complete-decoder symbols and ranges summarize three runs. Each no-image symbol represents one pretrained-vector ridge fit per cohort. Percentages show the mean paired-run fraction of the complete decoder’s Decima-over-random PCC gain retained by the ridge, using the complete decoder’s random-vector PCC as the common baseline.

The error decomposition showed that better gene-mean estimates accounted for 91.9–95.8% of the MSE reduction (**Fig. 3d**). Gene-mean PCC increased by 0.740–0.806 (**Fig. 3a**). Relative to each control error component, gene-mean MSE decreased by 57.1–67.2%, whereas centred MSE decreased by only 1.0–1.8%. Thus, Decima transferred mean expression much more strongly than variation across spots.

An independently fitted model recovered most of the matrix-level correlation gain without receiving an image (**Fig. 3e**; **Extended Data Fig. 3**). Ridge regression mapped each gene vector to one mean-expression value learned from training individuals and assigned that value to every test spot. With Decima vectors, this model achieved full-matrix PCCs of 0.338–0.450; random-vector and identity-shuffled controls remained near zero. Using the complete decoder’s random-vector performance as a common reference, the no-image models retained 90.2–99.3% of the complete decoder’s PCC gain.

Hippocampal *STX1B* illustrates the distinction (**Fig. 3b,c**). The Decima decoder predicted a mean of 1.47, close to the observed 1.56, whereas the random-vector decoder shown predicted 0.45. Yet their spatial PCCs were nearly identical: 0.29 and 0.28, respectively. The difference was primarily in expression level.

### 1.4 Spatial prediction gains vary across held-out genes

Across all eligible held-out genes, Decima improved pooled-spot mean within-gene PCC by only 0.0032– 0.0091 over random vectors. To test whether this average obscured stronger gains for some genes, we ranked held-out genes by their observed expression standard deviation in training individuals and divided them into deciles (Methods).

After centring observed and predicted expression separately within each test section and pooling the residuals (**Extended Data Fig. 4**), the Decima-over-random within-gene PCC effect ranged from approximately *−*0.003 to 0 in the lowest-variance decile and from 0.017 to 0.040 in the highest (**Fig. 4a**). Effects relative to the constant-vector control followed a similar, smaller trend. The spatial advantage therefore increased with training-tissue variation.

**Figure 4.**
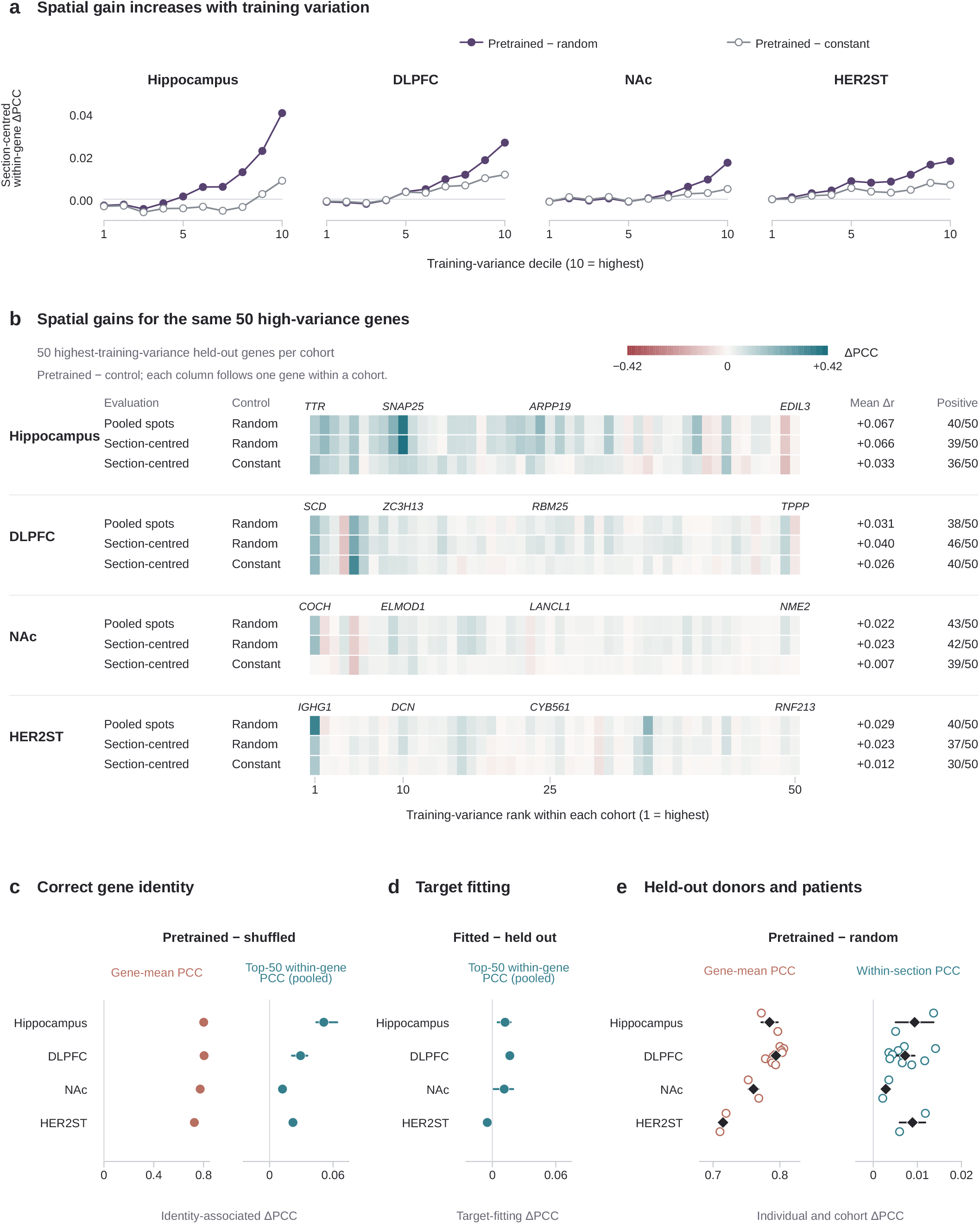
Spatial prediction gains vary with training-tissue expression variation. “Pretrained” denotes Decima-derived vectors. **a**, Mean section-centred within-gene PCC effects across cohort-specific deciles of training-spot standard deviation. Filled purple points compare pretrained and random vectors; open grey points compare pretrained and constant vectors. Effects were averaged across three paired runs within gene, then within decile. **b**, Effects for the 50 held-out genes with the highest training variance in each cohort. Columns follow training-variance rank, with the same gene in the same column across rows within a cohort. Gene symbols label training-variance ranks 1, 10, 25 and 50 in each cohort. Rows show pretrained-minus-random PCC effects over pooled spots, the same contrast using section-centred within-gene PCC, and pretrained-minus-constant effects using section-centred within-gene PCC. Centring removed observed and predicted means separately within each section before pooling spots. The colour scale is linear, symmetric and unclipped. Annotations show mean effects and counts of genes with positive effects. Effects were averaged across three paired runs within gene. Gene sets differ between cohorts. **c**, Effects of correct gene identity relative to assignments shuffled before training. Gene-mean PCC uses the full held-out panel; mean within-gene PCC uses pooled test spots for the fixed top-50 targets. **d**, Effects of including the previously held-out gene panel in decoder fitting, evaluated by pooled-spot mean within-gene PCC on the fixed top-50 targets. In **c,d**, points show means over three paired runs and lines span their ranges, with genes averaged within run first. Spatial axes have the same limits. **e**, Pretrained-minus-random effects on gene-mean and within-section PCC, averaging runs and, where applicable, sections within individual. Open circles denote test donors or patients (n = 2, 10, 2 and 2 for hippocampus, DLPFC, NAc and HER2ST); diamonds show equally weighted cohort means. Lines show 95% descriptive individual-block bootstrap intervals from 20,000 resamples. Metrics use all eligible held-out genes.

For the 50 highest-variance held-out genes in each cohort, Decima improved mean within-gene PCC by 0.022–0.067 across pooled test spots and by 0.023–0.066 for section-centred within-gene PCC (**Fig. 4b**; **Extended Data Fig. 5**). Between 37 and 46 of the 50 genes improved, depending on cohort and centring strategy. Gains over the constant-vector control were 0.006–0.035 and 0.007–0.033, respectively. These remaining gains distinguish gene-dependent spatial information from a map shared by all targets.

Correct gene–vector assignments contributed to this spatial signal. Compared with shuffling assignments before training, correctly aligned Decima vectors increased gene-mean PCC by 0.726–0.804 but mean within-gene PCC by only 0.0005–0.0061 over the full held-out panel (**Extended Data Fig. 6b**). For the fixed high-variance genes, the spatial gain increased to 0.012–0.051 and was positive in every run (**Fig. 4c**). Including the previously held-out gene panel in decoder fitting and checkpoint selection changed pooled within-gene PCC on the same top-50 targets by *−*0.0048 to 0.0166 (**Fig. 4d**; **Extended Data Fig. 6c**). Alternative target partitions, detection restrictions and gene-mean definitions preserved the broader ordering of mean and spatial gains (**Extended Data Figs. 3 and 7**).

The contrast also held across test donors and patients (**Fig. 4e**). Gene-mean gains were positive in every test donor or patient; within-section spatial gains were positive at the cohort level but much smaller.

### 1.5 Broad mean-expression transfer recurs with scGPT representations

We repeated the analysis using fixed gene representations from scGPT, which was pretrained on single-cell transcriptomes. Each gene was represented by a single vector, without cell-specific expression input. Representations were available for 3,574 held-out genes per brain cohort and 3,179 in HER2ST. Relative to dimension-matched random vectors, scGPT increased full-matrix PCC by 0.346–0.494 and gene-mean PCC by 0.805–0.947, but pooled-spot mean within-gene PCC by only 0.0006–0.0161. Gene means accounted for 91.5–95.3% of the MSE improvement (**Fig. 5a**). For scGPT, correct gene–token assignments were required for the large mean-expression gains, and no-image ridge models retained 81.8–90.4% of the full-matrix PCC gain above the complete decoder’s random-vector baseline (**Extended Data Fig. 9a,b**).

**Figure 5.**
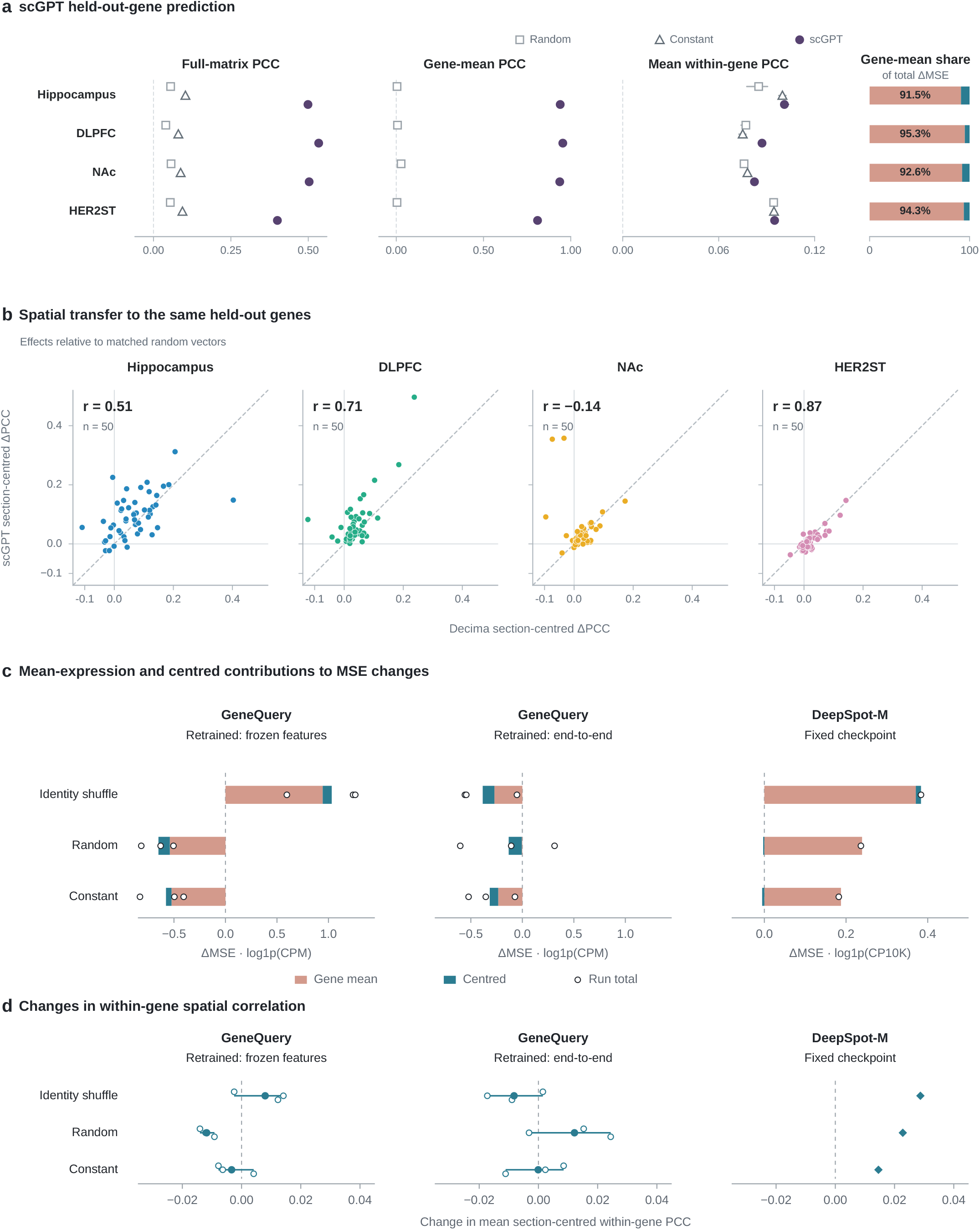
Mean-expression and spatial prediction across gene representations and models. **a**, Absolute full-matrix, gene-mean and mean within-gene PCC for static scGPT tokens and dimension-matched random and constant vectors. Evaluation includes 3,574 held-out genes per brain cohort and 3,179 in HER2ST. Symbols show means over three independently optimized runs; lines span their ranges. Constant-vector gene-mean PCC is undefined. Stacked bars show mean per-run shares of MSE improvement over random vectors attributable to gene-mean (terracotta) and centred (teal) error. **b**, Gene-level section-centred within-gene PCC effects for the same 50 highest-training-variance targets in each common held-out panel. Each axis shows pretrained-minus-random effects for one representation, averaged over three runs within gene. Panels have identical limits and equal aspect ratios. Labels give descriptive Pearson correlations between the two representation-specific effects across the same 50 targets. Dashed diagonals are identity lines. **c,d**, Gene-vector controls in GeneQuery and DeepSpot-M on HER2ST. GeneQuery uses 138 genes excluded from downstream fitting and three independently fitted runs per condition in each training setting. DeepSpot-M uses 135 targets exposed during upstream spatial training, with inference-time token controls in one fixed checkpoint. All targets satisfy the observed-expression eligibility rule for section-centred within-gene PCC. **c**, Signed gene-mean (terracotta) and centred (teal) contributions to control-minus-correct MSE; open points show run totals. Errors use log(1 + CPM) for GeneQuery and log(1 + CP10K) for DeepSpot-M, where CP10K denotes counts per 10,000, with separate axis limits. **d**, Correct-minus-control changes in mean section-centred within-gene PCC. GeneQuery open points show paired runs, filled circles their means and lines their ranges; diamonds show the fixed DeepSpot-M checkpoint. Positive values favour correct vectors in both panels. Supplementary Table S21 gives absolute performance.

Spatial effects differed between representations and cohorts. On the same 50 high-variance targets, scGPT-over-random section-centred within-gene PCC gains were 0.095 in hippocampus, 0.068 in DLPFC, 0.040 in NAc and 0.012 in HER2ST. Across the complete common held-out panels, they were 0.0159, 0.0131, 0.0057 and *−*0.0001 (**Extended Data Fig. 9d**). Gene-level spatial effects from Decima and scGPT correlated at 0.51, 0.71, *−*0.14 and 0.87 for the high-variance sets, respectively (**Fig. 5b**). The representations thus shared broad mean-expression transfer but differed in which genes gained spatial accuracy.

### 1.6 Mean-expression and spatial prediction in GeneQuery and DeepSpot-M

We next assessed mean-expression and spatial prediction separately in GeneQuery and DeepSpot-M on HER2ST (**Fig. 5c,d**; **Extended Data Fig. 10**; Supplementary Table S21). We reconstructed GeneQuery from its public implementation and retrained it with genes and patients excluded from fitting and selection. With the image encoder fixed, correctly matched descriptions reduced pooled-test MSE by 1.032 relative to shuffled descriptions; 0.944 (91.5%) came from gene-mean error. MSE did not improve relative to random or constant vectors. Mean section-centred within-gene PCC increased by 0.0080 relative to shuffled descriptions but decreased by 0.0118 and 0.0034 relative to random and constant vectors.

With end-to-end training, correctly matched descriptions increased gene-mean PCC by 0.352 but increased MSE by 0.385 relative to shuffled descriptions; MSE was higher in all three runs. Mean section-centred within-gene PCC decreased by 0.0082 relative to shuffled descriptions and increased by 0.0122 relative to random vectors. Thus, improved gene-mean agreement did not consistently yield better overall or spatial prediction in this reconstruction.

For DeepSpot-M, we evaluated gene-vector controls in the released model’s scGPT gene-representation pathway without retraining; the 135 queried genes had been exposed during upstream spatial training. Evaluation used log(1 + *x*)-transformed counts per 10,000, as reported for DeepSpot-M training. Relative to shuffled vectors, correct vectors reduced MSE by 0.3835, with 96.7% of the reduction attributable to gene-mean error. Mean section-centred within-gene PCC was 0.0720 with correct vectors, compared with 0.0433, 0.0492 and 0.0575 with shuffled, random and constant vectors, respectively. Relative to random and constant vectors, correct vectors therefore had higher spatial PCC but slightly higher centred MSE (**Fig. 5c,d**).

## 2 Discussion

We use an exact decomposition of prediction error to separate two parts of held-out-gene prediction that are usually evaluated together: a gene’s overall expression level and its variation across tissue. Our results show that pretrained gene representations help mean-expression prediction more broadly than spatial-pattern recovery. Across held-out genes and individuals, most of the overall improvement arose from better gene-mean estimates, whereas spatial gains depended on the target gene, representation and cohort. Predicting additional genes and recovering their spatial patterns therefore need to be assessed separately.

Pretrained gene representations can be learned from diverse molecular and knowledge sources, including genomic and protein sequence, transcriptomic profiles and biomedical text. Decima and scGPT were pretrained on different inputs—regulatory sequence and single-cell transcriptomes, respectively [15, 16]. Despite these different pretraining modalities, both representations improved gene-mean prediction for held-out genes in new individuals (**Fig. 3e**; **Extended Data Fig. 9b**). This gene-level signal could provide a useful baseline when extending spatial prediction to a larger gene panel. Gene-mean prediction also transferred across cohorts, with stronger agreement among the three brain cohorts than between brain and breast cancer (**Extended Data Fig. 3d**).

Spatial gains were more heterogeneous. Genes with greater expression variation in the training tissue tended to show larger performance gains, including after section-wise centring and relative to the constant-vector control. Of note, among high-variance targets, shuffling gene–vector assignments reduced these performance gains. Decima and scGPT also produced different gene-level spatial effects on the same targets across cohorts. Our grouping variable was observed expression variance in the training tissue, which reflects both biological variation and measurement noise and is not a formal measure of spatial variability.

The distinction between mean-expression and spatial prediction also has implications for model design and optimization. In DeepSpot-M, correct gene vectors increased within-gene spatial correlation, while most of the MSE reduction relative to shuffled vectors came from gene-mean error. In GeneQuery, higher gene-mean correlation did not consistently correspond to lower MSE. We saw the same separation in the fitted-gene experiments: mean initialization mainly affected gene-level expression, whereas neighbourhood information affected within-gene spatial prediction. Model components and training objectives can therefore affect gene-level abundance and within-gene spatial structure to different degrees. Objectives based on relative, differential or residual expression [14, 17–20] may better align optimization with spatial variation, but whether they generalize to genes and biological samples excluded from fitting remains to be tested.

There are limitations to our work. Our primary analyses tested genes and donors or patients excluded from downstream fitting and model selection within each cohort; related genes or tissues could have been present during representation pretraining. Cross-tissue spatial prediction and generalization to genes excluded from representation pretraining remain to be evaluated. GeneQuery and DeepSpot-M also have different training histories; in particular, the DeepSpot-M genes examined here had already been seen during upstream spatial training. Three cohorts contained only two test individuals, limiting population-level inference, and tissue, assay platform and image modality were not independently varied. In addition, richer image information or other learning objectives may yield different spatial gains. One open problem raised by these results is to identify which properties of the target gene, its representation and the tissue context determine whether spatial gains persist in unseen samples.

## 3 Methods

### 3.1 Study design, cohorts and partitions

We predicted spot-level gene expression from registered tissue images in four cohorts, fitting separate models within each cohort. Donors or patients were assigned to training, validation and test sets before model development. Data from training individuals were used to fit model parameters and calculate training-derived expression statistics. Data from validation individuals were used to select model checkpoints. The fitted-gene analysis compared direct per-gene heads with Decima-conditioned predictors. The held-out-gene analysis used a separate decoder family and excluded evaluation genes from both fitting and checkpoint selection. Primary comparisons used independently optimized runs with seeds 42, 123 and 456.

The hippocampus Visium analysis used 34 sections from ten donors in GEO accession GSE264692 [22, 23]: six donors (22 sections) for training, two (eight sections) for validation and two (four sections) for testing. Two of the 36 H&E capture areas listed in the source manifest were absent from the input archive. The dorsolateral prefrontal cortex (DLPFC) cohort, accession GSE307403 [24, 25], comprised one Visium section from each of 63 donors, partitioned into 44 training, nine validation and ten test donors. The nucleus accumbens (NAc) cohort, accession GSE307586 [26, 27], comprised 38 sections from ten donors: six donors (25 sections) for training, two (five sections) for validation and two (eight sections) for testing. Donors shared between hippocampus and NAc retained the same partition. Each brain cohort used an aligned 18,397-gene panel.

HER2ST comprised 36 sections from eight patients with HER2-positive breast cancer [28, 29]. Patients A–E (27 sections) supplied training data, F (three sections) validation data and G–H (six sections) test data. The gene universe was defined from training-patient matrices and mapped to 15,914 Decima-supported Ensembl genes. Cohort and partition sizes are summarized in Supplementary Table S1; individual and section assignments are provided in the released configuration files.

The primary 80/20 gene partition was constructed from mean log-expression across the 97,209 spots in the 22 final hippocampus training sections. The 18,397 genes were ordered by this statistic, divided into ten equal-rank bins and sampled at 20% per bin without replacement using seed 42. This yielded 14,717 training and 3,680 held-out brain genes; intersection with HER2ST yielded 12,730 and 3,184 genes, respectively. Validation and test expression did not contribute to this split. Training-individual expression for subsequently held-out genes was used to construct the split and variance rankings, but those genes were excluded from the spatial predictor’s loss and checkpoint criterion. Thus, gene exclusion refers to downstream predictor fitting; upstream representation pretraining could include the same genes or related tissues.

### 3.2 Expression preprocessing and gene alignment

For the primary analyses and GeneQuery, raw gene-expression counts were normalized per spot and transformed as

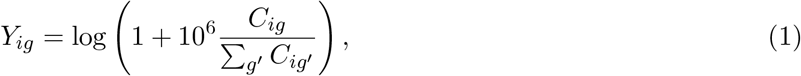

where *C*_*ig*_ is the raw count for gene *g* at spot *i*. Library-size denominators followed the fixed cohort preprocessing: hippocampus and DLPFC totals included all 36,601 source features before restriction to the aligned panel; NAc totals used its aligned panel; HER2ST totals used the 15,914-gene training-patient universe after zero filling. Performance differences between each candidate model and its matched control were calculated separately within each cohort, using the same evaluated genes and spots. Gene means are across-spot averages of these log(CPM + 1) values, where CPM denotes counts per million.

Visium analyses retained spots labelled <monospace>in_tissue=1</monospace>; HER2ST retained spots in the supplied selection files. No additional library-size, detected-gene or mitochondrial-fraction thresholds were applied. Visium genes, annotated using the 10x Genomics GRCh38 2020-A reference (GENCODE v32/Ensembl release 98), were aligned by exact Ensembl-identifier matching to the Decima-supported universe. HER2ST symbols were mapped only when unambiguous, restricted to genes present in training patients and reindexed in each section, assigning zero counts to absent features. Identifier handling and ordered lists are detailed in Supplementary Methods.

For each gene *g*, the training mean 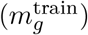 and standard deviation 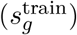 were calculated from normalized log-expression pooled across spots from all training individuals. Standard deviations were floored at 10^*−*3^ for use as fitted-gene scales. Training standard deviations also defined variance rankings. Anchors, scales and feature-standardization moments used training data only.

### 3.3 Frozen image and gene representations

UNI2-h, a ViT-H/14 pathology encoder pretrained on H&E and immunohistochemistry tiles [30, 31], provided a frozen 1,536-dimensional vector per spot. Hippocampus, NAc and HER2ST all used H&E images. DLPFC used the supplied three-channel RGB rendering of its immunofluorescence image. Spot-centred patches were bilinearly resized to 224 *×* 224 pixels and normalized with ImageNet [32] channel moments. Visium patch side lengths matched the spot diameter at the analysis image scale; HER2ST patches spanned 100 *µ*m. Fitted-gene models additionally received 31 spatial descriptors comprising an in-tissue indicator, array and pixel coordinates, normalized coordinates and their sine and cosine encodings (Supplementary Methods). These descriptors were concatenated with the image features and standardized using training-section moments. For spot-level input, held-out-gene models used image features without coordinate descriptors.

Decima [15] is a reference-sequence model with a Borzoi [33] backbone and transcriptomic supervision across cellular and disease contexts. We used Decima v0.5.1, model v1_rep0, on strand-aware 524,288-bp hg38 windows with nucleotide and gene-body channels. Averaging the final frozen feature map along the sequence axis produced one 1,920-dimensional vector per gene, shared across spots, individuals and cohorts.

For scGPT [16], we used the official whole-human checkpoint, described as pretrained on 33 million normal human cells. Each gene was represented by its 512-dimensional token-embedding row after checkpoint layer normalization. These were static vectors: no cell-specific expression or contextual transformer output was supplied. The brain-specific checkpoint was used only for sensitivity analysis. Supplementary Methods give extraction details, model revisions, file digests and pretraining-metadata checks; Supplementary Table S15 summarizes provenance and coverage. Packaged metadata do not enumerate all upstream studies, so related-tissue or downstream-source overlap cannot be excluded.

At inference, predictors received image patches at predefined locations and fixed gene vectors, without expression measurements or genotypes from the test individual. All analyses evaluated new individuals; held-out-gene analyses additionally evaluated genes excluded from downstream fitting and checkpoint selection.

### 3.4 Fitted-gene experiments

The direct predictor and Decima-conditioned predictor shared a 256-dimensional spot backbone with three message-passing layers. The direct model used a 128-unit head followed by trainable per-gene output weights and biases. Default initialization used Xavier output weights and zero biases. Mean-informed initialization instead used zero output weights and training-gene means as biases; both remained trainable.

The conditioned model used feature-wise linear modulation (FiLM) [34] to transform the shared spot representation according to the Decima vector. Its output was

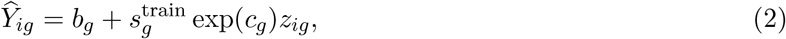

where *z*_*ig*_ is the raw spatial output for spot *i* and gene *g*, and *c*_*g*_ is a learned gene-level log-scale correction clipped to [*−*0.75, 0.75]. The anchor *b*_*g*_ was either the fixed training mean 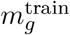 or a learned function of the Decima representation:

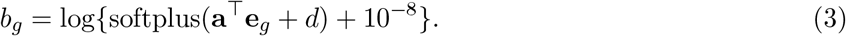

Here, **e**_*g*_ is the fixed Decima representation for gene *g*, **a** is a learned projection vector and *d* is a learned scalar intercept. The parameters **a** and *d* were optimized jointly with the spatial predictor under the same training objective. The spatial output *z*_*ig*_ was not constrained to have zero mean across spots, so both the anchor and the scaled spatial term could contribute to the predicted gene mean. We also considered replacing FiLM with concatenation and gene-to-spot cross-attention while retaining the graph-enabled backbone and training protocol.

Neighbourhood context used a directed six-nearest-neighbour graph within each section. Weights were exp(−∥**p**_*i*_ − **p**_*j*_∥ ^2^*/*0.1) where **p**_*i*_ and **p**_*j*_ are the section-normalized two-dimensional coordinates of spots *i* and *j*, respectively, and were row-normalized for message passing. The graph-enabled condition also included Laplacian smoothing. To isolate the contribution of neighbourhood information, the zero-edge control used the same architecture but removed interactions between spots: neighbour aggregation was set to zero and Laplacian smoothing was disabled, while spot-wise transformations and residual layers were retained. Thus, comparison with the graph-enabled model measures the joint effect of neighbour aggregation and Laplacian smoothing.

The loss was

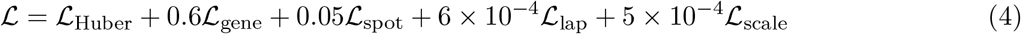

Here, *L*_Huber_ measures prediction error in normalized log-expression. The gene and spot terms are centred cosine losses across spots within genes and across genes within spots, respectively. The Laplacian term penalizes differences in raw spatial outputs between neighbouring spots, and the scale term penalizes squared gene-level log-scale corrections in conditioned models. Detailed definitions and implementation are provided in Supplementary Methods. Training used Adam [35], learning rate 2.5 *×* 10^*−*4^, weight decay 10^*−*4^ and at most 12 epochs, with a minimum of eight and early-stopping patience four. Each step paired one section with a chunk of at most 96 genes. Checkpoints maximized validation mean within-gene PCC plus 0.25 times full-matrix PCC.

We evaluated eight fitted-gene model variants to assess the effects of mean-expression information, neigh-bourhood context and gene conditioning (**Extended Data Fig. 1a**). Variants 1 and 2 were default direct heads without and with the graph; variants 3 and 4 added mean-informed initialization. Variant 5 was the spatially constant Decima-derived mean control. Variants 6 and 7 were Decima–FiLM without and with the graph, and variant 8 used the graph and fixed training-mean anchor. Mean-initialization comparisons were 3 versus 1 and 4 versus 2; neighbourhood comparisons were 2 versus 1, 4 versus 3 and 7 versus 6; conditioning comparisons were 6 versus 1, 7 versus 2 and 8 versus 4. Conditioning comparisons jointly change gene input, parameter sharing, decoder structure, scale correction and anchoring. Effects were calculated within cohort and seed before aggregation. Layer definitions, loss implementation and illustrative-map selection are given in Supplementary Methods.

### 3.5 Held-out-gene decoder and optimization

The held-out-gene decoder combined the frozen UNI2-h image vector **x**_*i*_ for spot *i* with the fixed gene vector **e**_*g*_ assigned to gene *g* under the corresponding representation or control condition:

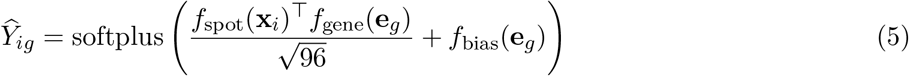

Here, *f*_spot_ and *f*_gene_ are trainable encoders that map their inputs to a shared 96-dimensional space, and *f*_bias_ is a trainable network that produces a scalar gene-specific bias. Spot and gene encoders used layer normalization, a 512-unit hidden layer, a Gaussian error linear unit (GELU), dropout 0.1 and projection to 96 dimensions. The gene-bias network used layer normalization and a 256-unit GELU hidden layer. The decoder operated independently on each spot, without coordinates or graph inputs.

Pretrained vectors were standardized feature-wise using training genes. Dimension-matched random controls used fixed standard-normal vectors; constant controls supplied a zero vector to every gene. Each condition was independently fitted with the same architecture, biological and gene partitions, and checkpoint-selection subset of training genes. Each training step sampled 256 spots and up to 512 training genes and used all sampled spot–gene pairs. The loss was mean squared error (MSE) plus 0.15 times one minus mean gene-wise PCC, with batch variances stabilized by 10^*−*6^.

Training used AdamW for six epochs, learning rate 2 *×* 10^*−*4^, weight decay 10^*−*4^ and gradient clipping at norm 1.0. Checkpoints maximized validation mean within-gene PCC minus 0.02 times MSE on 2,048 training genes sampled uniformly without replacement using the analysis seed. This subset was shared across vector conditions within cohort and seed. Held-out genes and test individuals did not contribute to fitting or selection.

The scGPT replication retained this protocol and removed out-of-vocabulary genes identically across all vector conditions. It used 14,248 training and 3,574 held-out genes per brain cohort, and 12,709 training and 3,179 held-out genes in HER2ST: held-out coverage of 97.1% and 99.8%. Standardization used covered training genes. Decima endpoints were recalculated without retraining on these same covered targets for representation comparisons, each against its own dimension-matched random control. Supplementary Methods specify randomization and batching details.

### 3.6 No-image and gene-identity controls

Independent ridge models mapped each frozen gene vector to mean log-expression in training individuals. The penalty was selected from {0.01, 0.1, 1, 10, 100, 1,000, 10,000} by minimum RMSE on an internal subset of downstream training genes. This fixed subset, sampled with seed 42, contained the smaller of 2,048 genes or one-fifth of the training set and was shared across conditions. Models were refitted on all training genes, negative predictions were clipped at zero, and each predicted mean was assigned to every test spot. Pretrained, dimension-matched random and identity-shuffled conditions used the same fitting procedure. For both Decima and scGPT, full-matrix correlation-gain retention used the complete decoder’s random-vector PCC as a common baseline: this value was subtracted from both no-image and complete-decoder pretrained-vector PCCs before taking their ratio. Ratios were calculated within cohort and seed, then averaged across seeds 42, 123 and 456. Each representation and cohort used one pretrained-vector ridge fit. Supplementary Methods give the formula. Cross-cohort transfer applied the source-cohort ridge and standardization unchanged to target-cohort held-out genes.

Identity-shuffled controls permuted pretrained vectors separately within training and held-out genes before fitting. Fixed assignments were maintained through training, selection and testing. This preserved each partition’s vector multiset and covariance while disrupting gene correspondence. The rule was shared by Decima and scGPT spatial predictors and ridge models; each seed supplied an independent permutation and fit.

A genomic-covariate ridge and covariate-adjusted gene-mean correlations assessed information beyond simple annotations (Supplementary Methods; Supplementary Table S18).

The target-fitted reference used the same Decima vectors, decoder, image features and biological partitions, but allowed primary held-out genes into fitting and checkpoint selection. Evaluation remained restricted to those targets in test individuals. Besides all eligible genes and training-defined high-variance sets, exploratory top-quartile and top-decile strata used observed test variation.

### 3.7 Component-resolved evaluation and aggregation

For each gene, observed and predicted test means and centred values were

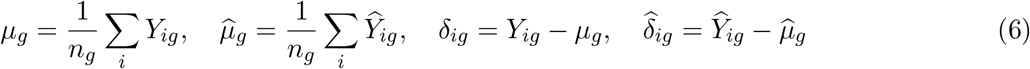

Here, *n*_*g*_ is the number of evaluated test spots for gene *g, µ*_*g*_ and 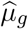 are the observed and predicted test-set means, and *δ*_*ig*_ and 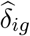 are deviations from those respective means. The same test spots were used for every gene within cohort and run, giving the exact decomposition

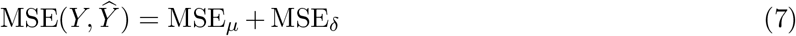

The mean component averaged 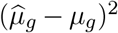 across genes; the centred component averaged 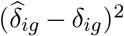 across spot–gene pairs. If finite observation counts differed between genes, the mean term was weighted by *n*_*g*_ to preserve the identity.

Gene-mean PCC correlated means across genes, and gene-mean RMSE was the square root of their mean squared difference. Constant-vector decoders predict the same mean for every gene, making gene-mean PCC and its contrasts undefined; their MSE and RMSE remained defined. Pooled within-gene PCC correlated expression across all test spots for each gene without section centring. Eligibility depended only on observed standard deviation exceeding 10^*−*6^ and was shared across matched conditions. Eligible genes with predicted standard deviation at or below 10^*−*6^ contributed within-gene PCC zero, retaining a common denominator. Mean pooled within-gene PCC (abbreviated mean within-gene PCC in figures and tables) averaged these scores equally across eligible genes. Centred RMSE used all evaluated centred spot–gene values.

Full-matrix PCC correlated all finite spot–gene pairs after flattening, without the within-gene eligibility filter. Gene-centred full-matrix PCC instead removed each eligible gene’s observed and predicted means before flattening, retaining weighting by spatial variance and observation count.

Correlation effects were candidate minus control. For each specified candidate–control comparison, error reductions were defined as ΔMSE = MSE_control_ *−* MSE_candidate_, with the same convention for the mean and centred components. We reported the gene-mean share of a positive MSE improvement, Δ MSE_*µ*_ */*Δ MSE, and reductions normalized to each control component, Δ MSE_*µ*_ */* MSE_*µ*,control_ and Δ MSE_*δ*_ */* MSE_*δ*,control_. The share describes the source of an improvement and can exceed 100% when centred error increases.

Primary evaluation pooled all test spots within each cohort. For section-centred analyses, observed and predicted expression were separately centred within each test section and gene. We then used two correlation summaries. Section-centred within-gene PCC concatenated the centred spots across sections within each gene before calculating PCC and then averaged PCC equally across eligible genes; **Fig. 4a,b** and **Fig. 5b** use this endpoint. Within-section PCC instead calculated gene-wise PCC separately within each section, averaged across eligible genes, and then averaged first within biological individual and then equally across individuals; this endpoint was used for individual-level summaries. MSE components were aggregated before taking square roots. An observation-weighted summary pooled squared errors across spot–gene pairs. All evaluations used existing checkpoints without refitting.

For GeneQuery and DeepSpot-M, mean section-centred within-gene PCC averaged per-gene correlations of pooled section-centred residuals using the same observed-expression eligibility rule. Section-centred full-matrix PCC instead correlated the flattened residual matrices. Matrix-level and gene-mean PCCs and their contrasts were reported as missing when either vector’s standard deviation was at or below 10^*−*6^, including centred no-image predictions. The MSE decompositions in the main text and **Fig. 5c** used means over pooled test spots. By contrast, **Extended Data Fig. 10a** decomposed MSE within each section and averaged components across each patient’s three sections; GeneQuery components were then averaged across runs. Patient-level shares were ratios of the averaged mean-component and total reductions. These summaries use different centring units.

### 3.8 Target stratification and sensitivity analyses

Held-out genes were ranked within cohort by training-expression standard deviation, a pooled statistic that includes within-section variation, shifts among sections or donors and measurement noise. The top 50 genes were fixed across conditions and seeds and evaluated over pooled test spots and after section centring. All were covered by scGPT. A complementary analysis divided observed-eligible targets into ten fixed training-variance deciles. Effects were calculated within gene and seed, averaged across runs and summarized within each set. Decima–scGPT comparisons used identical covered targets; same-gene effects were compared using Pearson and Spearman correlations. Gene-level distributions summarize variation among correlated targets.

Four alternative training-only gene partitions tested expression-stratified resampling (seeds 123 and 456), embedding-cluster separation and chromosome blocking. The cluster split assigned entire Decima-vector clusters within expression bins to training or held-out sets; the chromosome split withheld chromosomes 1, 14, 15 and X. Each was independently fitted in one paired seed-42 pretrained-versus-random run per cohort. We also summarized nearest-training-vector similarity and its post hoc association with performance. Cluster construction, split-balance checks and exact assignments are described in Supplementary Methods.

A historical gene-partition sensitivity, whose stratification was not fully independent of the final biological split, is documented separately in Supplementary Methods and Supplementary Table S19.

Additional analyses replaced mean log-expression with log {1 + mean(CPM) } or restricted targets by observed test detection: at least 10% or 25% of spots, presence in every test section, or above-median detection fraction. Detection subsets were shared across conditions and affected evaluation only. Illustrative maps were selected using documented marker-based and performance screens, including post hoc selection of *STX1B*; the full rules and selection records are given in Supplementary Methods and Source Data.

### 3.9 Decoder architecture and pathway analyses

We independently fitted bias-free, concatenation and residual-only decoders under the primary biological and gene partitions with the same frozen inputs and vector controls. The bias-free model replaced the gene-bias network with a single shared intercept. The concatenation multilayer perceptron (MLP) jointly processed normalized spot and gene vectors without a gene-only skip branch; its 2,441,345 parameters closely matched the primary decoder’s 2,371,777. Both retained the primary loss and checkpoint criterion. The residual-only decoder instead fitted exactly section-centred targets, centred its signed outputs during training, validation and testing, and selected checkpoints using within-section PCC and centred MSE. Layer definitions and parameter counts are given in Supplementary Methods and Supplementary Table S12.

Frozen-checkpoint analyses suppressed either the bias or interaction logit before softplus, or permuted held-out gene identities independently in the two branches. Parameters remained fixed. The branches were fitted jointly, and their nonlinear readouts do not add to the full prediction. Intervention and constant-prediction conventions are specified in Supplementary Methods; results appear in **Extended Data Fig. 8b–d** and Supplementary Tables S10 and S11.

### 3.10 External implementation analyses

We reconstructed GeneQuery’s public gene-aware pipeline [11] using its HER2 description vectors and verified their recovered gene-row order. Of 785 public-panel symbols, 34 lacked an unambiguous Decima-metadata match; intersection with the fixed split retained 613 training and 138 held-out genes. Patients A–E were used for fitting, F for selection and G–H for testing. The query head combined projected ResNet-50 image features and description vectors through the public two-layer gene transformer. We fitted both frozen-image and end-to-end variants, with semantic, identity-shuffled, random and constant vectors independently trained for seeds 42, 123 and 456. Evaluation genes and test patients were excluded from fitting and selection. Both variants used AdamW, learning rate 10^*−*4^, weight decay 10^*−*3^, at most 100 epochs and early-stopping patience 12, selecting minimum validation MSE.

For DeepSpot-M [13], we evaluated the released full checkpoint through its scGPT pathway on 601 fitted-panel and 135 held-out-panel genes and 3,097 test spots from patients G–H. The same image features were decoded with correct, partition-shuffled, Gaussian or repeated fitted-panel-mean gene tokens, changing both the query adapter and gene router inputs. These were inference-time interventions in one checkpoint whose target tokens had been trainable during upstream spatial training. We evaluated its unmodified outputs against unsmoothed counts normalized to 10,000 per spot and log-transformed, following the documented training scale. The library-size denominator retained the fixed 15,914-gene HER2ST universe.

For each external representation, an independent no-image ridge was fitted to training-patient gene means on the corresponding spatial model’s expression scale, using standardized, correctly assigned gene vectors. The penalty was selected on an internal 20% fitted-gene subset before refitting on all fitted genes. Negative predictions were clipped at zero. All predictions used the endpoint and constant-prediction conventions above. Supplementary Methods provide source revisions, checkpoint digests, preprocessing and reconstruction details.

Two secondary analyses supplied fitted-gene context: a donor-disjoint hippocampus comparison with adapted ST-Net, Hist2ST, BLEEP and STPath implementations (**Supplementary Table S4**), and recovery of a training-defined HER2ST tumour program in annotated sections from test patients (**Supplementary Table S5**). Their methods and results are given in Supplementary Methods and Supplementary Note 5.

### 3.11 Statistical summaries and computational reproducibility

Optimization seeds measure computational variability, including control-vector assignments where applicable. Figures summarize runs as specified in their legends; displayed ranges span the minimum and maximum. No hypothesis tests treated the three runs as biological replicates.

Biological-individual uncertainty used section-centred paired pretrained-versus-random effects averaged across runs within section. We used 20,000 bootstrap draws, either resampling individuals after averaging their sections or resampling individuals and then sections within each selected occurrence. Leave-one-individual-out summaries assessed sensitivity to each test individual. Target-level resampling used the 100 Decima-vector clusters as blocks, preserving within-cluster dependence. Intervals were the 2.5th and 97.5th percentiles. Three cohorts had only two test individuals, so their intervals describe finite-sample sensitivity.

Sample sizes were determined by the available public resources. All accessible sections meeting the input requirements were retained; no section, individual or target was excluded from quantitative analyses because of test performance. Individual partitions were fixed before fitting. Investigators were not blinded to cohort, partition or condition. This study was a secondary computational analysis of de-identified source-study data; no new participants were recruited or specimens collected. Ethics approvals and consent procedures are reported in the original cohort publications [22, 24, 26, 28].

Analyses used Python, NumPy, SciPy, pandas, scikit-learn and PyTorch on Linux; neural-network jobs used Slurm and NVIDIA GPUs. Primary analyses used Python 3.11. The release includes configuration files, a version-pinned CPU environment and installation and integration records. Supplementary Methods describe the research environments and the scope of these execution checks.

### 3.12 Use of generative artificial intelligence

During manuscript preparation, the authors used ChatGPT and Codex (OpenAI) for language editing, document organization and code review. The authors checked all scientific claims, numerical results, references, figures, tables and source-data records against the underlying analyses and take full responsibility for the final content.

### 3.13 Data availability

The human hippocampus Visium data are available from the Gene Expression Omnibus (GEO) under accession GSE264692. The DLPFC Visium cohort is available under GSE307403; associated access conditions are described in that record. The NAc Visium data used here are available under GSE307586, with processed data at Zenodo doi:10.5281/zenodo.17089020. HER2ST count matrices, histology images, spot coordinates and pathology annotations are available from Zenodo at doi:10.5281/zenodo.4751624. Source data are provided for **Figs. 1–5** and **Extended Data Figs. 1–10**.

### 3.14 Code availability

Code for model fitting, held-out-gene evaluation and the manuscript analyses is available at https://github.com/tjchen020524/SpatioS2E. The repository includes analysis scripts, input-preparation tools, biological-individual and gene partitions, environment specifications and reproduction instructions.

Third-party model weights are obtained from their original providers under the applicable access and license terms.

### 3.15 Author contributions

T.C.: conceptualization, methodology, software, validation, formal analysis, investigation, data curation, visualization and writing—original draft. S.C.H.: conceptualization, methodology, supervision, project administration, funding acquisition and writing—review and editing.

## 3.16 Acknowledgments

Computational resources were provided by the Joint High Performance Computing Exchange (JHPCE), a high-performance computing facility in the Department of Biostatistics at the Johns Hopkins Bloomberg School of Public Health. We thank the brain donors and their families, the patients who participated in HER2ST, and the investigators and staff who generated and shared the hippocampus, DLPFC, NAc and HER2ST datasets.

---

**Extended Data Fig. 1.**
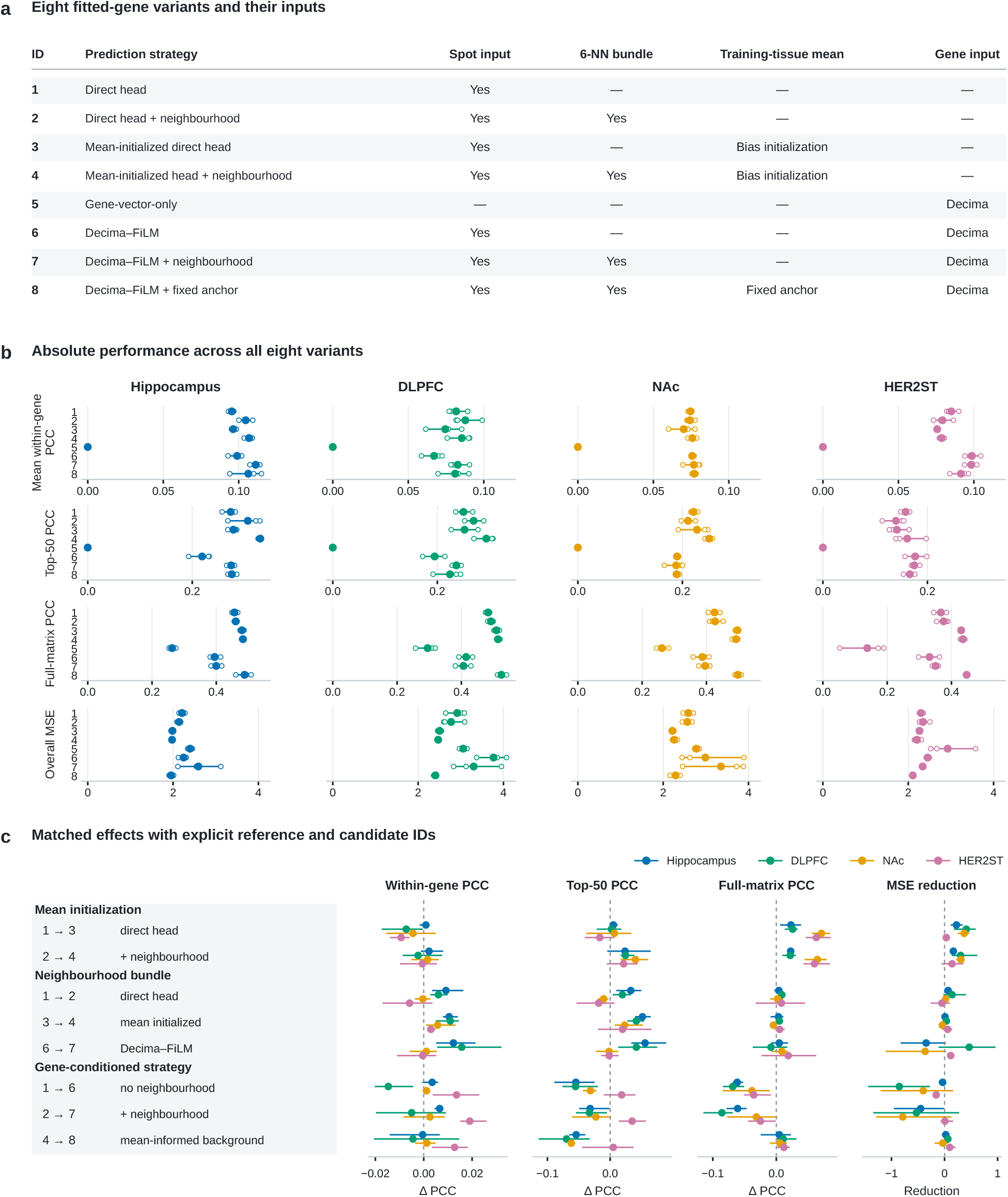
Model changes have different effects on mean-expression and spatial prediction for fitted genes. **a**, Eight variants differing in spot input, training-tissue-mean information, neighbourhood context and gene conditioning. In variants 3 and 4, trainable output biases are initialized to training-tissue gene means and output weights to zero; in variant 8, the training-tissue mean is a fixed additive anchor. The 6-NN intervention combines aggregation and Laplacian smoothing. **b**, Absolute mean within-gene PCC, top-50 PCC, full-matrix PCC and overall MSE. Rows within each axis follow variant IDs 1–8 from top to bottom; columns identify cohorts. Filled points show means, open circles show three independently optimized runs and lines span the run values. Axes are shared across cohorts within each metric. **c**, Matched effects, with arrows identifying reference and candidate variants. Correlation effects are candidate minus reference; MSE effects are reference minus candidate. Positive values favour the candidate. Points and lines show the mean and range over three runs on fixed biological partitions and gene panels. Supplementary Table S2 provides complete values.

**Extended Data Fig. 2.**
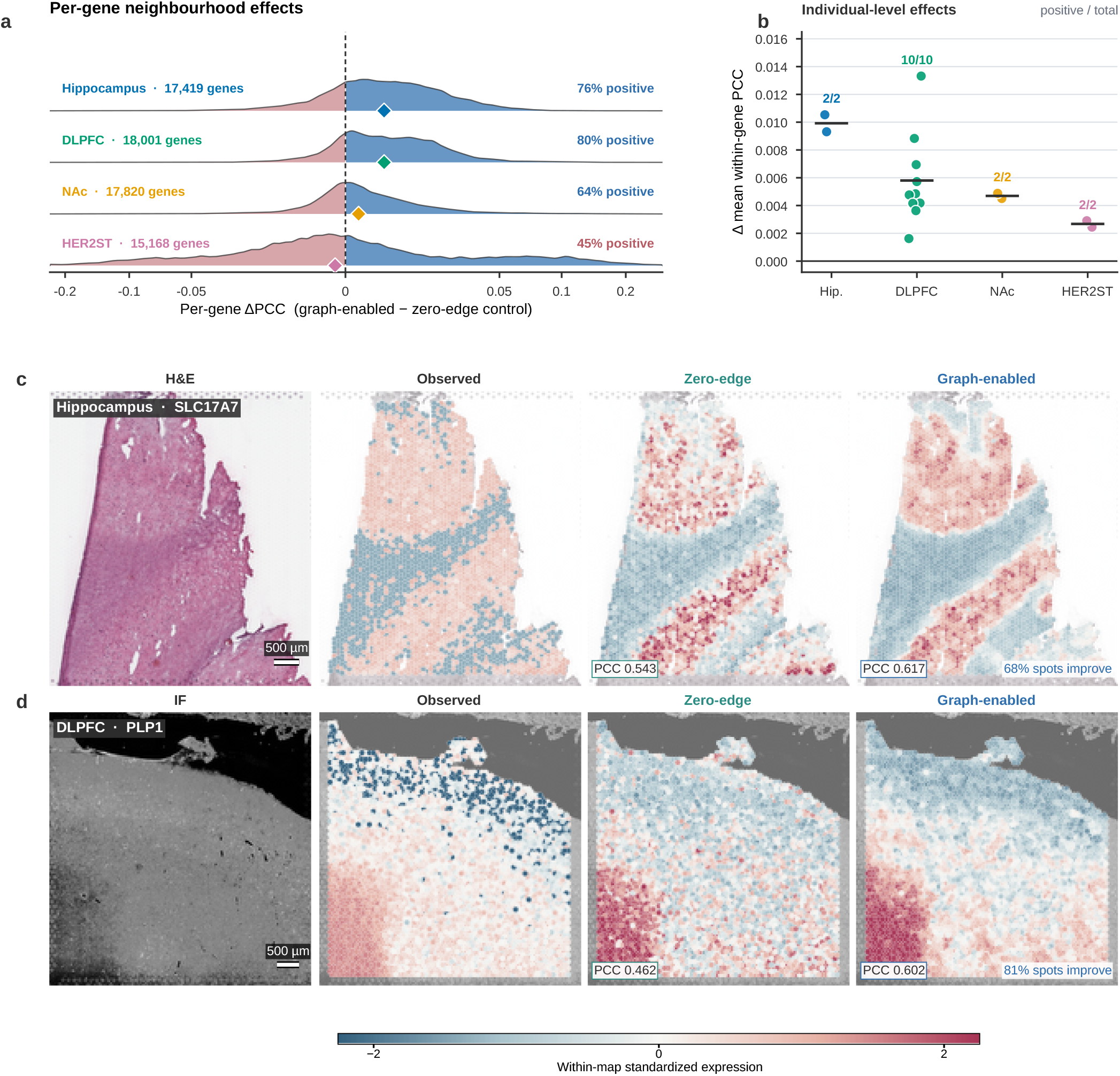
Neighbourhood context gives heterogeneous fitted-gene spatial gains. **a**, Per-gene PCC changes after adding 6-NN aggregation and Laplacian smoothing to the mean-informed direct model. Paired effects were averaged over three runs within gene; diamonds mark medians and labels give the fraction of positive effects. **b**, Mean within-gene PCC effects in each test individual after averaging paired runs. Donors or patients are the biological units. **c,d**, Illustrative hippocampal *SLC17A7* and DLPFC *PLP1* examples for the neighbourhood comparison (variant 4 versus 3), selected as described in Supplementary Methods and displayed using seed-42 predictions. Maps were standardized separately within gene, section and model for display; printed PCCs use unstandardized expression. Scale bars, 500 *µ*m.

**Extended Data Fig. 3.**
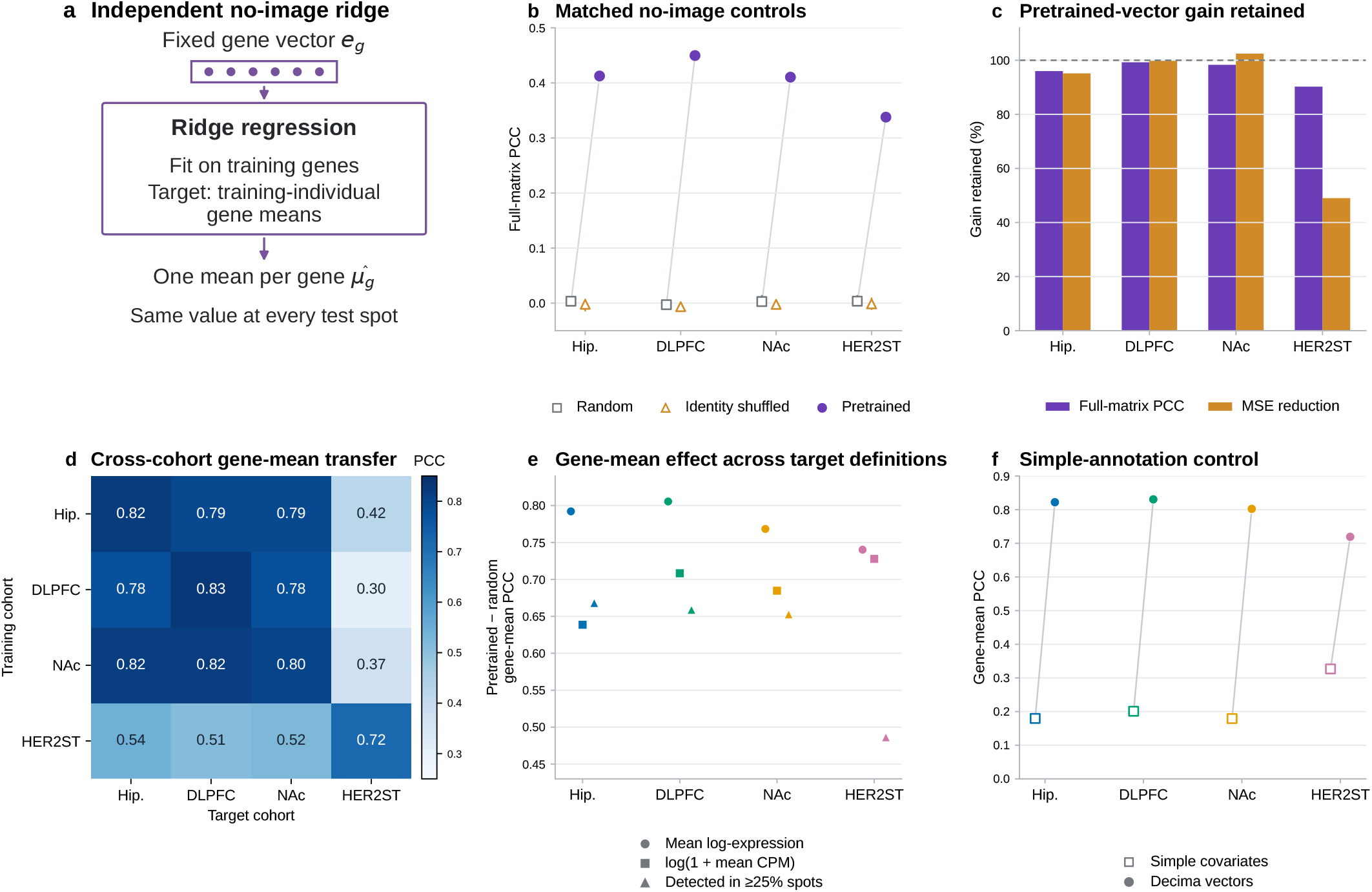
No-image models retain most of the matrix-level gain from pretrained gene vectors. **a**, An independent ridge model is fitted on training genes to predict their mean log-expression across spots from training individuals. It assigns one predicted mean to each held-out gene at every test spot, without image, coordinate or other spot inputs. **b**, Full-matrix PCC for pretrained, random and identity-shuffled no-image models. **c**, Fractions of the complete decoder’s Decima-over-random PCC gain and MSE reduction retained without images, using the complete decoder’s random-vector performance as the common baseline for both models. Percentages are averaged over three seed-paired comparisons. **d**, Cross-cohort gene-mean transfer without refitting in the target cohort. **e**, Complete-decoder pretrained-versus-random effects under alternative gene-mean definitions and target-inclusion rules. **f**, Gene-mean PCC for independently fitted no-image ridge models using simple genomic covariates (open squares) or Decima vectors (filled circles), evaluated on the same held-out targets. Both models use training-individual gene means and each yields one deterministic result. Supplementary Table S18 reports their absolute performance and gene-mean agreement after subtracting covariate predictions. Complete decoders in panel e use three optimization runs; repeated random-vector ridge controls in panel b vary vector realization.

**Extended Data Fig. 4.**
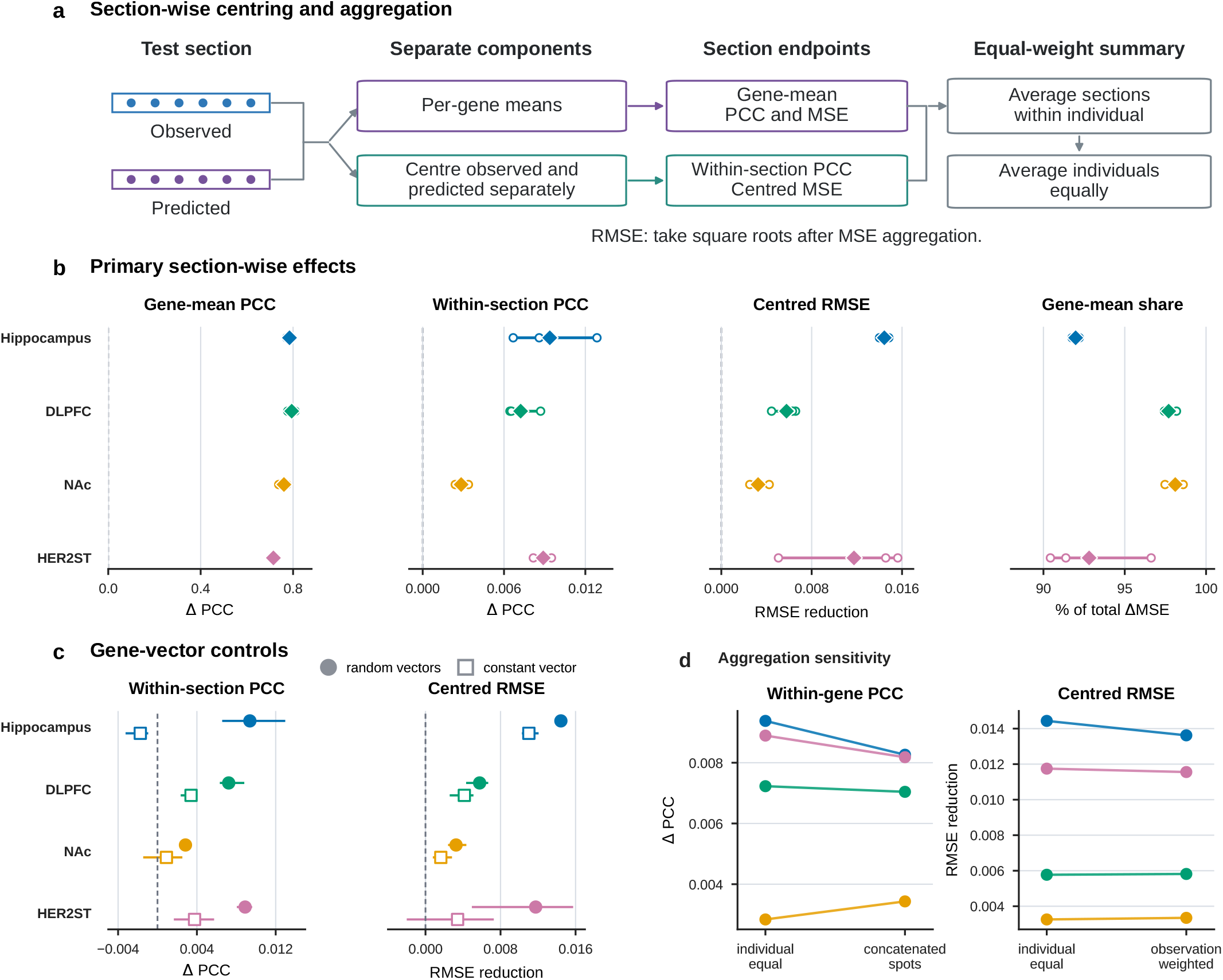
Section-wise centring preserves the contrast between mean and spatial transfer. **a**, Observed and predicted expression were centred separately for each gene within each test section. Gene means were evaluated across genes, and centred predictions across spots within genes. Section-level metrics were averaged within individuals and then equally across individuals; MSE components were aggregated before taking square roots to obtain RMSE. **b**, Decima-minus-random effects calculated within section, averaged within individual and then equally across individuals. **c**, Within-section PCC gains and centred-RMSE reductions relative to random and constant controls. **d**, For PCC, the equal-individual summary is within-section PCC; the concatenated-spots summary is section-centred within-gene PCC, calculated after pooling section-centred residuals within each gene. For RMSE, equal-individual summaries are compared with observation-weighted squared errors. Open symbols show three optimization runs; filled symbols show their mean. Lines span the run values.

**Extended Data Fig. 5.**
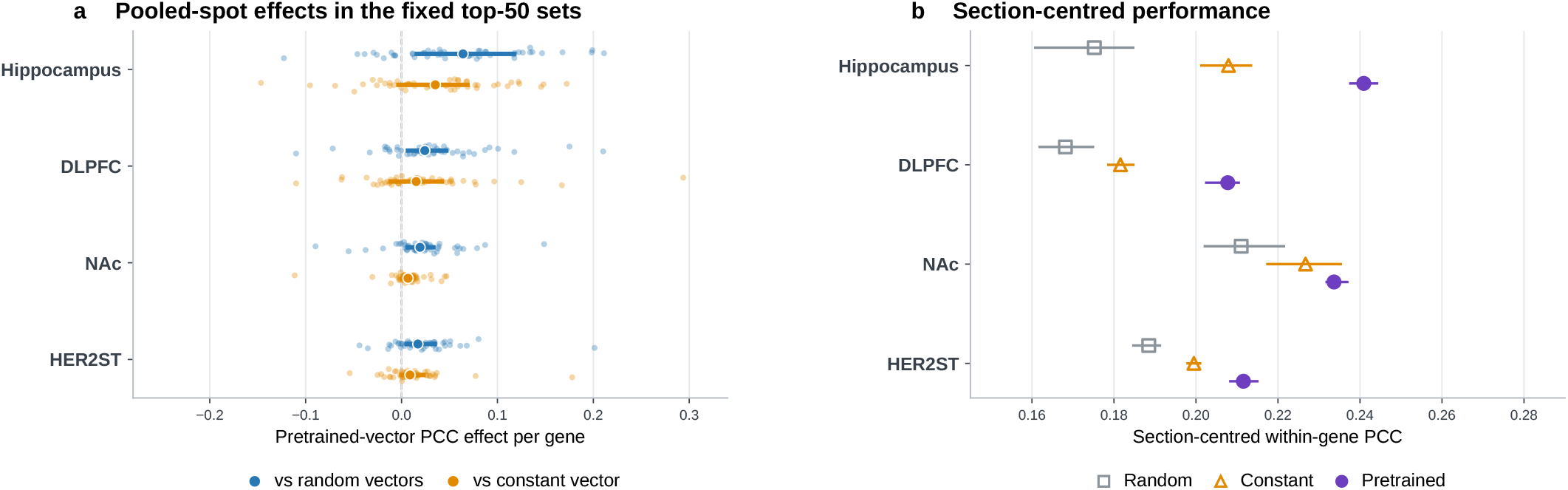
High-variance held-out genes show selective spatial gains. **a**, Pooled-spot within-gene PCC effects for the 50 held-out genes with the highest training-spot standard deviation in each cohort. Small points show gene-level effects averaged over three paired runs; larger points and thick lines show medians and interquartile ranges. **b**, Absolute section-centred within-gene PCC, calculated by removing observed and predicted gene means separately within each test section, concatenating the residuals within gene, then averaging gene-wise PCCs. Gene-level distributions describe variation among 50 correlated targets. **Figure 4c,d** shows identity and target-fitting effects for these genes; complete comparisons are in Supplementary Table S14 and Source Data.

**Extended Data Fig. 6.**
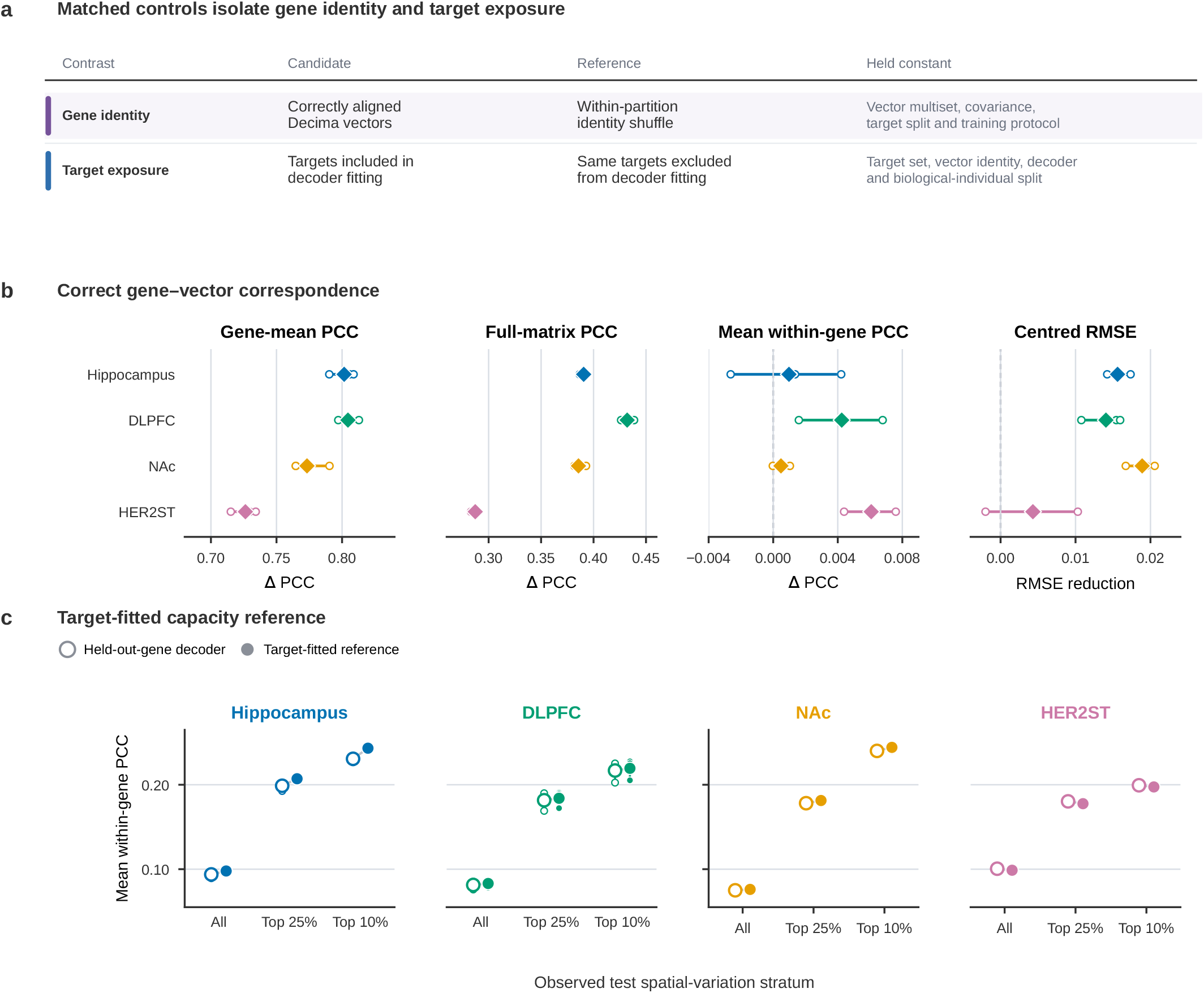
Correct gene identity mainly improves mean prediction, while target fitting adds modest spatial gains. **a**, Control design. Training-time identity shuffling preserves each partition’s Decima-vector multiset and covariance while breaking gene–vector correspondence. The target-fitted reference retains the same evaluation genes, vectors, decoder and individual split but includes evaluation genes in fitting and checkpoint selection. **b**, Effects of correct identity relative to shuffling. Correlation effects are pretrained minus shuffled; centred-RMSE reductions are shuffled minus pretrained. Positive values favour correct identity. Small open points show three runs; diamonds and lines show means and ranges. **c**, Absolute mean within-gene PCC for held-out and target-fitted models over all eligible genes and the top quartile or decile of observed test-set standard deviation. Small points show runs; large points and lines show means and ranges. These outcome-defined strata describe performance conditional on test signal. The target-fitted comparison measures the effect of including evaluation genes in fitting under the same model and image representation. Colours identify cohorts.

**Extended Data Fig. 7.**
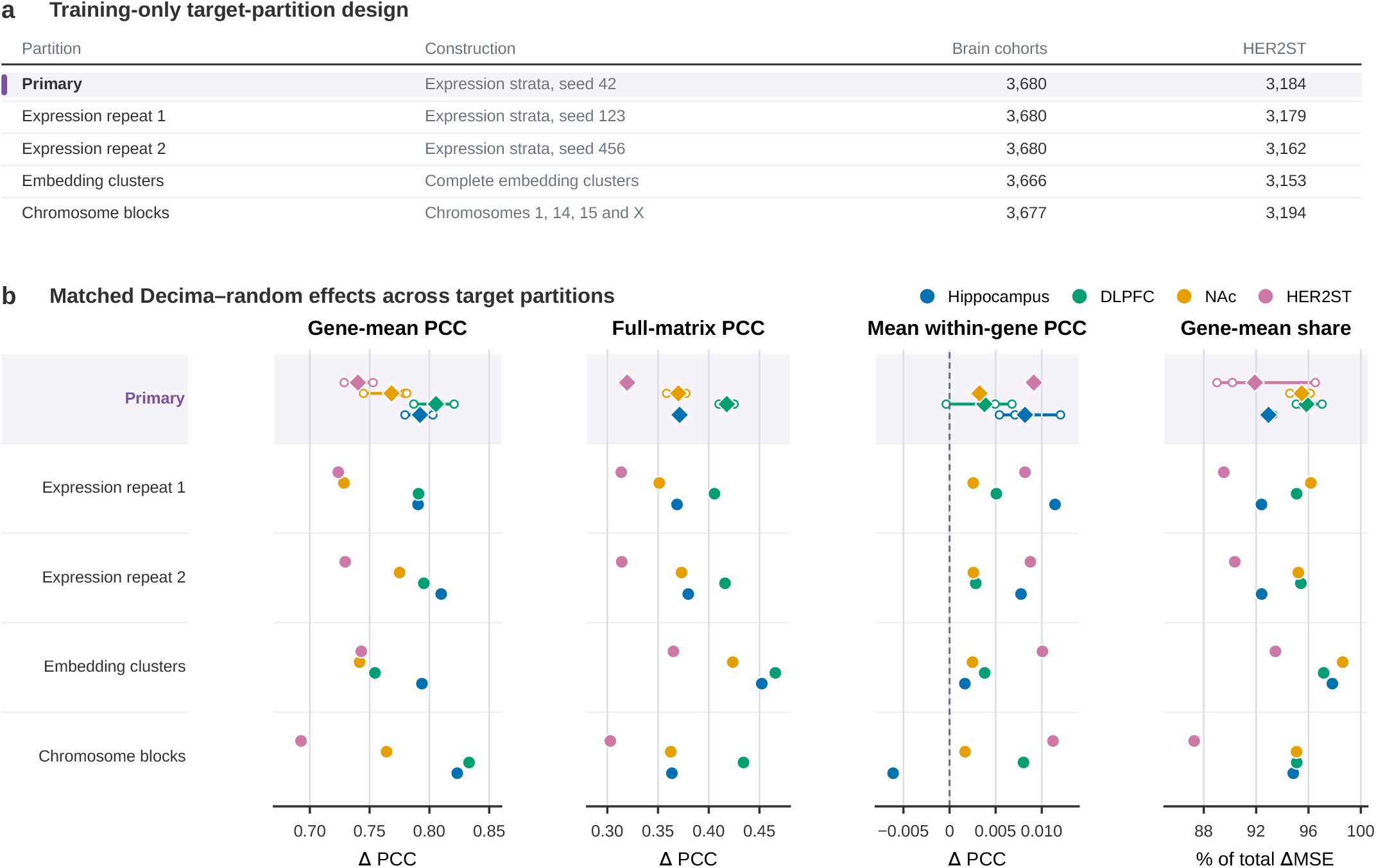
Mean-expression gains persist across training-only gene partitions. **a**, Five gene partitions: the primary expression-stratified split, two independent expression-stratified repeats, an embedding-cluster-disjoint split and a chromosome-blocked split. Construction used only final training individuals where expression statistics were required. The 18,397-gene master panel defines brain-cohort assignments; HER2ST counts show intersections with its 15,914-gene universe. Complete embedding clusters did not cross the training–held-out boundary. Nearest-training-gene vector similarities are reported in Source Data. **b**, Decima-minus-random effects after independent fitting under each split: gene-mean PCC, full-matrix PCC, mean within-gene PCC and the gene-mean share of MSE reduction. Colours identify cohorts. Primary-split diamonds show means, open points show three runs and lines span their ranges. Each alternative split used one paired seed-42 run per cohort, shown without an interval. All splits use the same pretrained Decima vectors.

**Extended Data Fig. 8.**
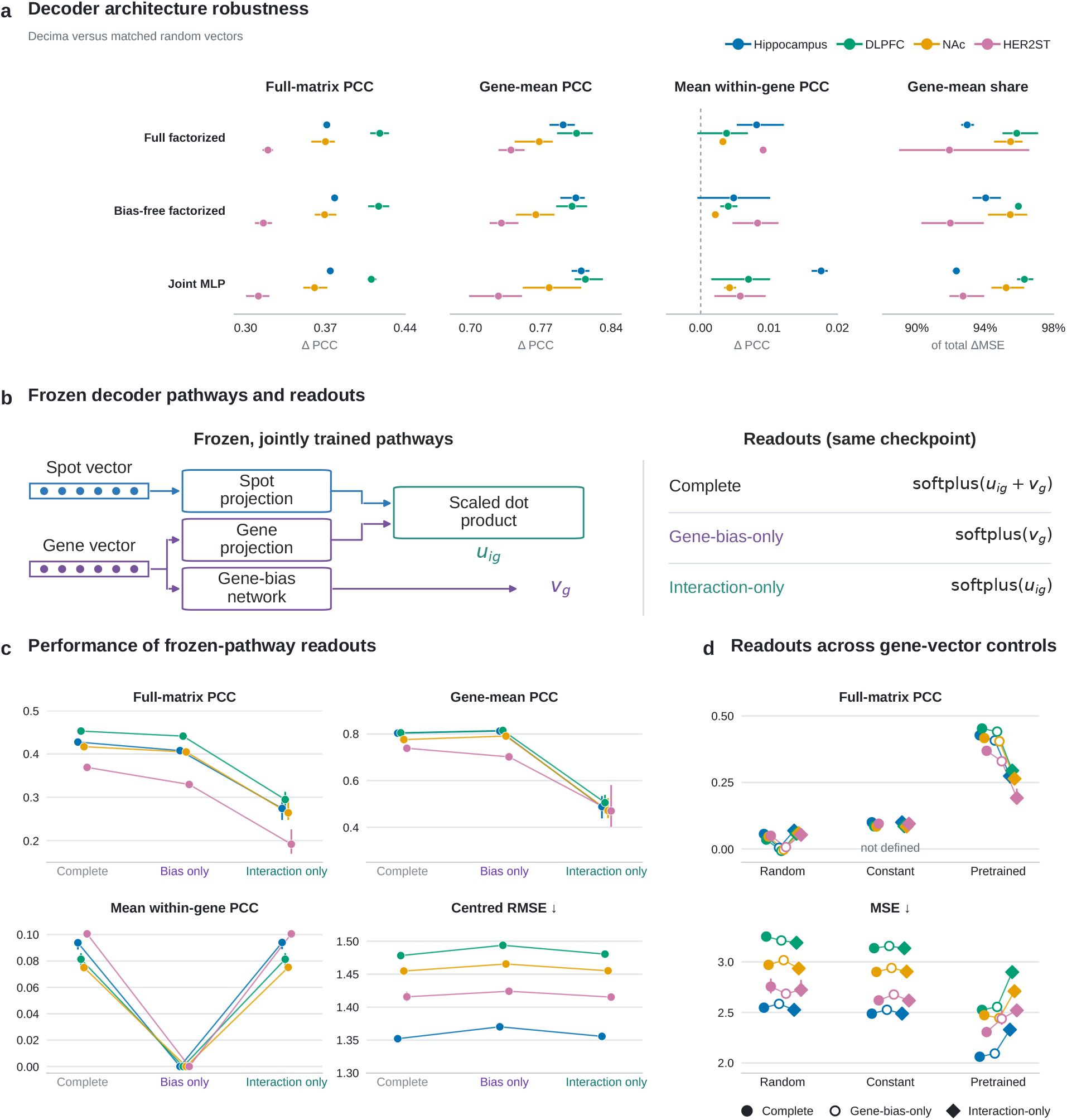
Decoder comparisons and pathway readouts distinguish mean prediction from spatial recovery. **a**, Decima-over-random effects for the complete factorized decoder, a bias-free factorized decoder with one shared intercept, and a parameter-matched joint MLP. Columns show full-matrix, gene-mean and mean within-gene PCC gains and the gene-mean share of MSE reduction. Architectures were independently trained on the primary split. **b**, Readouts from jointly trained, frozen primary decoders. With interaction logit *u*_*ig*_ and gene-bias logit *v*_*g*_, complete, gene-bias-only and interaction-only predictions are softplus(*u*_*ig*_ + *v*_*g*_), softplus(*v*_*g*_) and softplus(*u*_*ig*_), respectively. Each isolated readout sets the other logit to zero before softplus, without refitting parameters. **c**, Metrics for these readouts with Decima vectors. Bias-only predictions are constant across spots and contribute within-gene PCC zero under the observed-expression eligibility rule. **d**, Full-matrix PCC and MSE for all three vector conditions. Constant-vector bias-only full-matrix PCC is undefined because the entire prediction matrix is constant. Colours identify cohorts; points and lines show means and ranges over three runs. Panels **b–d** evaluate the jointly trained pathways in isolation, with checkpoint parameters held fixed.

**Extended Data Fig. 9.**
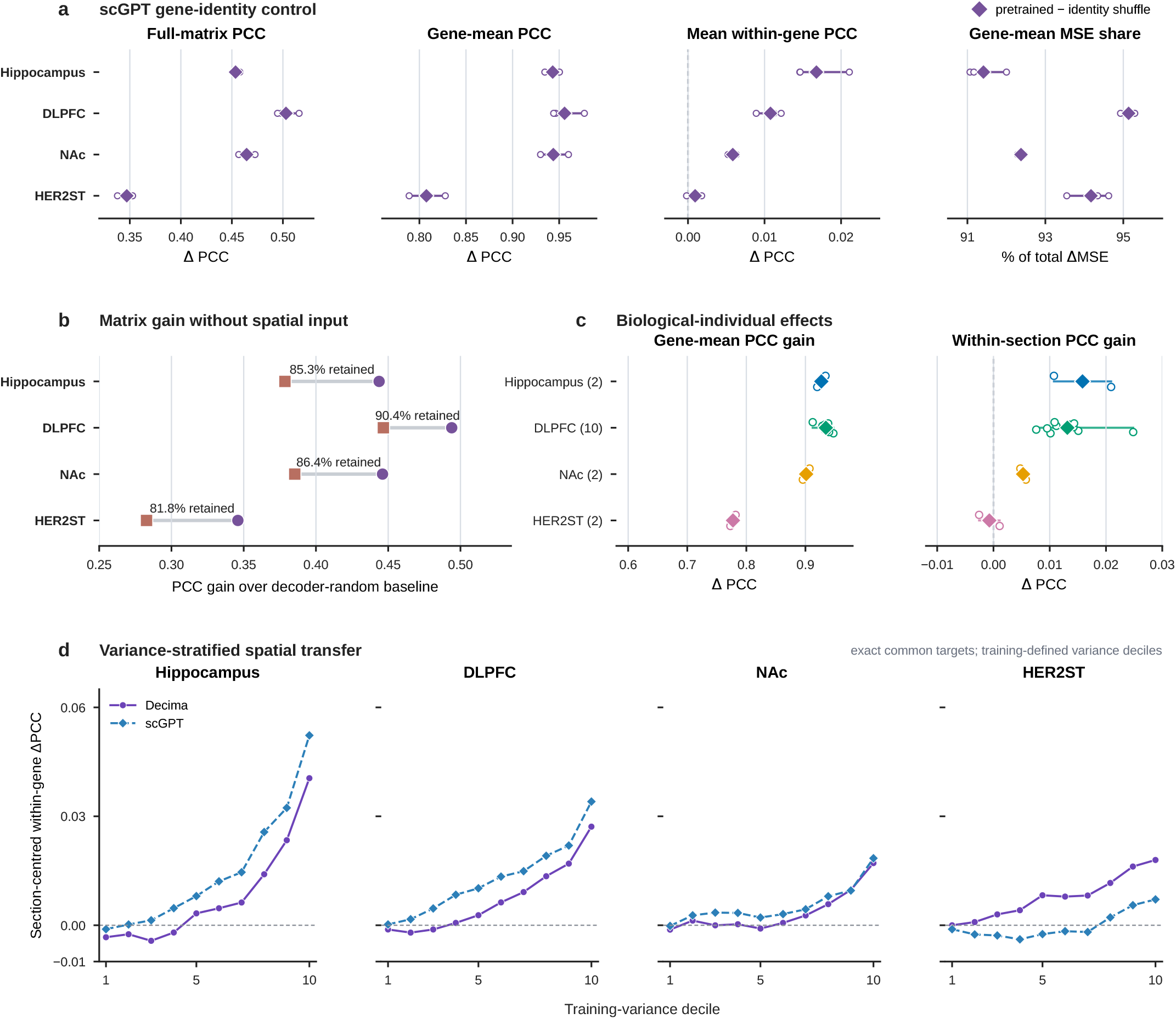
Static scGPT tokens reproduce broad mean-expression transfer and selective spatial gains. **a**, Correctly aligned scGPT tokens compared with assignments shuffled before training. Open points show three runs; filled points and lines show means and ranges. Gene-mean MSE share is the fraction of improvement over this control attributable to gene means. **Figure 5a** shows random-control performance; Supplementary Table S15 gives provenance and coverage. **b**, Full-matrix PCC improvements from complete decoders (circles) and independent no-image ridge models (squares), both relative to the complete decoder’s random-vector PCC. Points show mean improvements over three runs; labels show the mean seed-paired fraction retained without images. The same common-baseline definition is used for Decima in **Fig. 3e. c**, scGPT-minus-random effects after averaging runs and sections within each individual. Open points show donors or patients, diamonds show equally weighted cohort means, and lines span the observed individuals. Parentheses give individual counts. **d**, Section-centred within-gene PCC effects across training-variance deciles of the exact common held-out panels. Observed and predicted values were centred within section and concatenated within gene before calculating PCC. Solid purple lines show Decima; dashed blue lines show scGPT. Each is compared with its dimension-matched random control. Effects were averaged across runs within gene, then within decile. Axes are shared across cohorts and include negative effects. **Fig. 5b** compares effects on the same high-variance genes.

**Extended Data Fig. 10.**
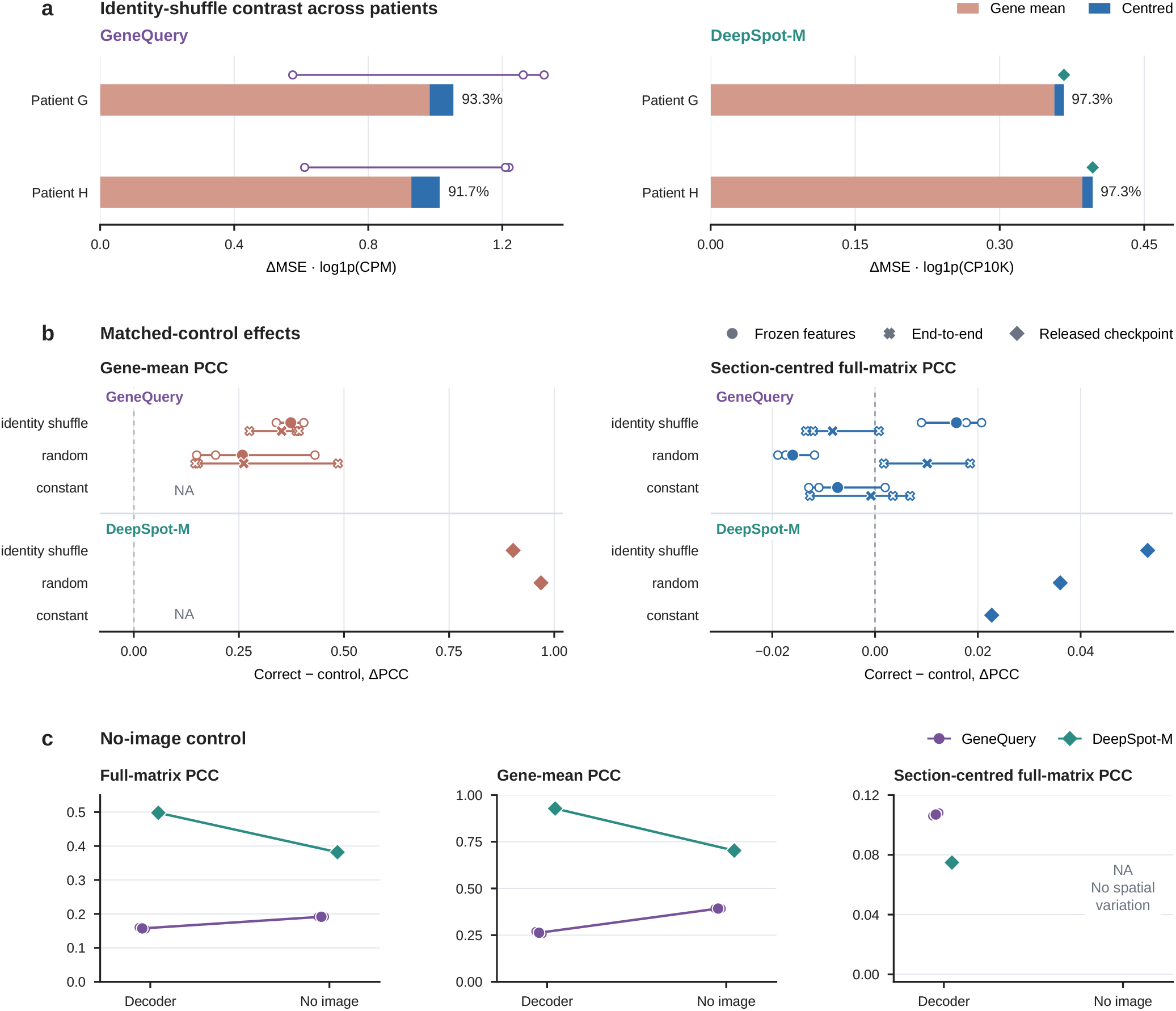
Mean-expression and spatial effects of gene-vector controls in GeneQuery and DeepSpot-M. **a**, Correct-versus-identity-shuffle MSE attribution in HER2ST patients G and H. Components were calculated within each section, then averaged over the patient’s three sections and across runs. Bars show these mean contributions; points show run totals or the fixed checkpoint. Percentages are ratios of the mean gene-mean contribution to the mean total reduction. The main text and **Fig. 5c** instead use gene means over all pooled test spots. GeneQuery uses 138 downstream-held-out genes and three frozen-feature runs. DeepSpot-M uses 135 panel-held-out genes in its released checkpoint, whose token rows were trainable during upstream spatial training. Both were evaluated on six sections and 3,097 spots. GeneQuery MSE uses log(1 + CPM); DeepSpot-M MSE uses log(1 + CP10K). Axes have different limits. **b**, Gene-mean and section-centred full-matrix PCC effects, calculated as correct minus control, for identity-shuffled, random and constant vectors. Constant-vector gene-mean PCC and its contrasts are undefined (NA). Open circles and filled circles show individual frozen-feature GeneQuery runs and their means. Offset open crosses and larger filled crosses show individual end-to-end runs and their means; lines span the three-run minimum and maximum for each training setting. Diamonds show the fixed DeepSpot-M checkpoint. GeneQuery controls were independently fitted; DeepSpot-M interventions were applied at inference. **c**, Absolute full-matrix, gene-mean and section-centred full-matrix PCC for models with correctly assigned gene vectors and independently fitted no-image models; GeneQuery uses frozen-feature runs. No-image predictions lack within-section variation, making section-centred full-matrix PCC undefined (NA). Comparisons are within implementation, as target coverage, expression scale, training and upstream exposure differ. Mean section-centred within-gene PCC effects appear in **Fig. 5d**; Supplementary Table S21 gives absolute results across all vector conditions.

