## Supplementary Information for "Pretrained gene representations transfer mean expression more broadly than spatial patterns in virtual spatial transcriptomics"

#### Contents

| Item | Content |
| --- | --- |
| Extended Data | **Figures 1–10** are included in the manuscript file: fitted-gene calibration, neighbourhood context, no-image prediction, section centring, high-variance targets, gene identity, gene partitions, decoder architecture and pathways, scGPT replication and GeneQuery and DeepSpot-M analyses |
| Supplementary Methods | Input processing, model extraction and implementation, sensitivity analyses, spatial-example selection, external analyses and computational records |
| Supplementary Notes | Five notes covering analyses not reported in full in the main Results |
| Supplementary Tables | Tables S1, S3, S4, S6, S7, S9, S10, S12–S18, S20 and S21 are included in this PDF; Tables S2, S5, S8, S11 and S19 are supplied in the workbook |
| Workbook | **Chen_Hicks_Supplementary_Tables.xlsx** contains searchable sheets for Tables S1–S21; full-precision run-level values remain in Source Data |

### Supplementary Methods

#### Input processing and gene identifiers

The two hippocampal capture areas absent from the input archive were Br2743 V11U08-081\_C1 and V11U08-081\_D1. DLPFC samples were 10-μm fresh-frozen Visium sections; HER2ST used first-generation Spatial Transcriptomics sections.

For the Visium cohorts, source feature tables supplied Ensembl identifiers and gene symbols from the 10x Genomics GRCh38 2020-A reference (GENCODE v32/Ensembl release 98). We used the released Decima gene metadata, whose published preprocessing workflow combined annotations from the 10x Genomics GRCh38 2020-A and 2024-A references with supplementary mappings from NCBI Gene [1]. We retained Gene Expression rows and intersected Ensembl identifiers with the Decima-supported identifiers by exact string matching, storing the intersection in lexicographic order. We did not strip version suffixes or substitute aliases. When one identifier had conflicting source symbols, the first mapping in the sorted feature-file traversal was retained for annotation; model alignment used the Ensembl identifier. Hippocampus and DLPFC used the same ordered 18,397-gene intersection. NAc retained this order for genes present in every processed section.

HER2ST matrices were indexed by gene symbol; the source study used GRCh38 with Ensembl release 86 [2]. We mapped symbols through the Visium-Decima intersection only when a symbol occurred once, discarding ambiguous mappings. The resulting universe was restricted to mapped symbols observed in patients A–E. Ensembl identifiers were required to be unique; duplicate features were not aggregated. Each section was reindexed to this fixed universe, with absent features assigned zero counts. The exact ordered gene lists accompany the manuscript configuration record.

The released configuration files in `configs/manuscript` contain cohort assignments and ordered gene lists for the primary partition. The `gene_splits_sensitivity` subdirectory contains the lists for the historical and alternative partitions. The accompanying manifest records list sizes, checksums and primary–historical overlap.

#### Training-derived anchors and scales

All expression-derived priors used training individuals only. For fitted gene $g$, the training-tissue anchor and scale were

$$m_g^{\text{train}} = \frac{1}{N_{\text{train}}} \sum_{i \in \text{train}} Y_{ig}, \quad (1)$$

$$s_g^{\text{train}} = \max \left\{ \left[ \frac{1}{N_{\text{train}}} \sum_{i \in \text{train}} (Y_{ig} - m_g^{\text{train}})^2 \right]^{1/2}, 10^{-3} \right\}. \quad (2)$$

Training spots were pooled across individuals for these statistics. Validation and test spots did not contribute.

#### Representation extraction and provenance

##### Image patches and UNI2-h

UNI2-h is a Vision Transformer H/14 (ViT-H/14) pathology encoder pretrained on H&E and immunohistochemistry tiles [3, 4]. Hippocampus, NAc and HER2ST used H&E images; DLPFC used the supplied immunofluorescence image. DLPFC files were loaded as their supplied three-channel red–green–blue (RGB) renderings and received the same resizing and ImageNet normalization as H&E images. We loaded MahmoodLab/UNI2-h checkpoint `pytorch_model.bin` at Hugging Face revision `d517a8dd47902dd7c308b3c36f63bce47e7b9a43`, with SHA-256 digest `6e077eda234bebc595868d918d3458d9dd32a050199b0ff04443b2f46a0a3b1e`, using strict state-dictionary matching.

For Visium, square patches were centred on spots, with side length equal to the Space Ranger full-resolution spot diameter at the analysis image scale. Boundary crops were reflection-padded. HER2ST patches had a side length of 100 μm, half the median nearest-centre spacing, and were padded with white at image boundaries. Patches were bilinearly resized to $224 \times 224$ pixels and normalized with ImageNet channel moments. UNI2-h produced one 1,536-dimensional vector per spot and remained frozen in all four cohorts.

Fitted-gene models also received 31 coordinate descriptors: an in-tissue indicator; array row and column; pixel row and column; section-wise min–max-normalized x and y coordinates; and sine and cosine expansions of each normalized coordinate at frequencies 1, 2, 4, 8, 16 and 32. These descriptors were concatenated with the UNI2-h vector. Image and coordinate features were standardized using training-section moments.

### Reference-sequence vectors

Decima is a reference-sequence model built on a Borzoi backbone and trained with transcriptomic supervision across cellular and disease contexts [1, 5]. We used Decima v0.5.1, model v1\_rep0, from local checkpoint rep0.ckpt with SHA-256 digest `9b4efc2967d09d05c34ced1877744ad1d499d3899863463d5107d072046bfb31`.

The reference was soft-masked hg38 from the University of California, Santa Cruz (UCSC), assembly GCA\_000001405.15. Alternative contigs were filtered with genomepy v0.16.3. The FASTA SHA-256 digest was `dbb2cdcc81772c7a1df59cca5b1321048d531f8e88e4ad5ae6162e52b71f4814`. Decima metadata defined a strand-aware 524,288-bp window per gene, extending by default 163,840 bp upstream and 360,448 bp downstream of the annotated gene start, with shifts near chromosome ends. Four channels encoded nucleotides and a fifth marked the gene body.

The final frozen feature map was averaged along the sequence axis to obtain a 1,920-dimensional gene vector. Reference inputs were shared across individuals; individual genotypes, somatic variants and allele-specific features were not supplied.

### scGPT token extraction

We used the official scGPT whole-human checkpoint, described in its model-zoo metadata as pretrained on 33 million normal human cells [6]. The packaged best\_model.pt file had SHA-256 digest `6cb5d451ab5c4b33eb673adbe4fddc61d2389df1b89b7651a9fe2e557572b922`. Its 60,697-token vocabulary had digest `acca93d114ca62c3f0f50debbd23e8c87f0714f4737764454f6b2b13f2e8580f`, and its argument file had digest `c18e075e018140cb8b2d9029387b9de26607a5ce6a8ccabd6ead70cd76b95d60`. No revision identifier was supplied; these digests identify the files used.

For each gene, we extracted the 512-dimensional row from encoder.embedding.weight and applied encoder.enc\_norm with epsilon $10^{-5}$. The decoder received fixed token vectors, without cell-specific expression or contextual transformer outputs. Extraction used local scGPT commit `cebd6fae655b9c585a4807daa3ac31bb764f06b4`. Packaged metadata did not enumerate all pretraining datasets, so overlap with related tissues or downstream source studies cannot be excluded. The brain-specific checkpoint was used only in a checkpoint-sensitivity analysis.

### Pretraining metadata and vocabulary coverage

For Decima, we searched packaged supervision metadata for downstream accessions and cohort names in tissue, organ, disease, study, dataset, region and subregion fields. We separately recorded other profiles labelled hippocampus, prefrontal cortex, nucleus accumbens or breast. Gene metadata identified whether each downstream held-out gene belonged to the upstream training, validation or test gene partition. The upstream gene-partition counts and identifier-search outcome are summarized in Supplementary Table S15. These checks document recorded pretraining exposure; string matching cannot establish absence from every related or derivative dataset.

For scGPT, we recorded checkpoint, vocabulary and argument-file digests; upstream data path and model-zoo description; token extraction and normalization; vocabulary coverage by cohort and gene partition; and all out-of-vocabulary genes. Covered and missing genes were compared using training-only mean, standard deviation, detection fraction and biotype. Because coverage can depend on these properties, representation comparisons used identical covered held-out targets. Packaged metadata did not provide a complete accession-level study list, preventing exclusion of all overlap with related downstream resources.

### Model implementation and optimization

#### Fitted-gene layers and conditioning

The main Methods define the expression factorization and matched variants. The learned anchor was optimized only through the spatial-expression loss. A 128-unit multilayer perceptron (MLP) mapped the gene vector to the log-scale correction and was initialized to return zero.

Concatenated spot features were projected to 256 dimensions. Three message-passing layers combined separate linear transformations of each spot state and its weighted neighbour aggregate, followed by layer normalization, a rectified linear unit (ReLU), dropout 0.1 and a residual connection. A final 256-dimensional MLP followed message passing. FiLM used a one-hidden-layer network with 256 hidden units to map $\mathbf{e}_g$ to channel-wise scale and shift parameters for the shared spot representation [7]. A shared 128-unit output network produced $z_{ig}$; its final layer was initialized to zero. FiLM retained a common spatial backbone while introducing multiplicative and additive gene-dependent interactions. The Decima network remained frozen.

To test sensitivity to the conditioning operator, we replaced FiLM with either a one-hidden-layer MLP on concatenated spot and gene features or four-head gene-to-spot cross-attention over four learned tokens projected from each spot representation. The concatenation model used algebraically equivalent separate linear projections of spot and gene vectors, followed by summation, ReLU and projection to 256 dimensions. Cross-attention used the gene vector as query and spot-feature tokens as keys and values, followed by spot and gene skip projections and layer normalization. Both alternatives retained the frozen inputs, spatial backbone, graph setting, expression factorization, optimization protocol and three seeds of the graph-enabled FiLM model.

Auxiliary delta-supervision weights were zero, and $z_{ig}$ could have a nonzero across-spot mean. The anchor $b_g$ and spatial term therefore both contribute to the predicted gene mean. Gene means and centred components were calculated from exported predictions.

The fitted-gene implementation reserved additional cell-composition input channels, fixed to zero in all experiments. The spatial graph used no radius cutoff; outgoing edge weights were normalized by their row sum during message passing.

#### Fitted-gene losses and batching

Huber beta was 1.0. The gene loss was one minus centred cosine similarity across spots within each gene; the spot loss was the analogous quantity across genes in the minibatch. Centred norms were stabilized at $10^{-8}$, and genes or spots with observed centred norm at most $10^{-6}$ were excluded from the corresponding average. Laplacian loss was the edge-weighted squared difference between raw spatial outputs, divided by the number of edges and genes in the minibatch. Direct-model gene biases cancel in neighbour differences. The scale penalty was the mean squared gene-conditioned log-scale correction and was absent from the direct model.

Each optimization step paired all retained spots in one section with one gene chunk. Training sections were shuffled between epochs. Gene order was independently shuffled and traversed in balanced chunks of at most 96 genes, with the cursor continuing across sections and resetting only after the full gene list had been visited. Hippocampus, DLPFC, NAc and HER2ST used 9, 5, 8 and 7 chunks per section, respectively; across training sections, this schedule visited every gene at least once per epoch. Validation and testing used chunks of at most 128 genes. Adam [8] used learning rate $2.5 \times 10^{-4}$, weight decay $10^{-4}$, coefficients (0.9, 0.999), epsilon $10^{-8}$ and gradient clipping at norm 1.0. Training lasted at most 12 epochs, with a minimum of eight and early-stopping patience four. No learning-rate scheduler or automatic mixed precision was used. Checkpoints maximized validation mean within-gene PCC plus 0.25 times full-matrix PCC.

#### Held-out-gene batching and optimization

Training spots were pooled into one shuffled manifest. Each step sampled 256 spots and, independently, up to 512 training genes without replacement; predictions and losses used all sampled spot–gene pairs. The final incomplete spot batch was discarded. The correlation loss was one minus mean gene-wise PCC across the 256 spots, with predictions and observations centred and $10^{-6}$ added to variances before taking square roots. The objective was MSE plus 0.15 times this correlation loss.

Models were trained for six epochs with AdamW, learning rate $2 \times 10^{-4}$, weight decay $10^{-4}$, coefficients (0.9, 0.999), epsilon $10^{-8}$ and gradient clipping at norm 1.0. No scheduler or automatic mixed precision was used. Checkpoints maximized validation mean within-gene PCC minus 0.02 times MSE over 2,048 training genes sampled uniformly without replacement using the analysis seed. This subset was not expression-stratified and was shared by vector conditions within cohort and seed. A new subset was drawn for each seed. Held-out genes and test individuals did not contribute to fitting or selection.

For the held-out-gene correlation loss, the implemented batch estimator for gene $g$ was

$$r_g = \frac{B^{-1} \sum_i (\hat{Y}_{ig} - \bar{\hat{Y}}_g)(Y_{ig} - \bar{Y}_g)}{\{B^{-1} \sum_i (\hat{Y}_{ig} - \bar{\hat{Y}}_g)^2 + 10^{-6}\}^{1/2} \{B^{-1} \sum_i (Y_{ig} - \bar{Y}_g)^2 + 10^{-6}\}^{1/2}}, \quad (3)$$

where $B$ is the number of spots and $G$ the number of genes in the batch; the correlation loss was $1 - G^{-1} \sum_g r_g$. Python, NumPy and PyTorch random-number generators were seeded, including available CUDA devices. cuDNN benchmark mode remained enabled and deterministic algorithms were not forced; analysis seeds consequently characterize computational variation rather than guarantee bitwise equality across accelerator stacks.

### Random and identity-shuffled vectors

For analysis seed $s$, dimension-matched standard-normal vectors were drawn with seed $s + 12,345$ and remained fixed throughout fitting and evaluation. Constant controls supplied zero vectors. NumPy's default generator used seed $s + 810,001$ to permute pretrained vectors separately within training and held-out partitions. Assertions verified preservation of each partition's vector multiset. scGPT used the same rules after vocabulary filtering, with 512-dimensional random vectors. Every vector condition was independently optimized; no weights were shared across conditions.

### Sensitivity analyses and numerical summaries

#### Gene partitions and split balance

We assessed split balance using training mean and standard deviation, gene length, reference-window guanine-cytosine (GC) fraction, standardized Decima-vector norm, biotype and chromosome. Numeric variables were summarized by held-out-minus-training standardized mean differences and 5th, 50th and 95th percentiles. Categorical variables were summarized by proportions and total-variation distance. This descriptive assessment did not alter the split.

The embedding-cluster-disjoint partition standardized Decima vectors, reduced them to 50 principal components using randomized principal component analysis (PCA) with seed 42 and then fitted ten MiniBatchKMeans clusters separately within each of the ten training-expression bins. MiniBatchKMeans used batch size 512, ten initializations, at most 300 iterations, reassignment ratio 0 and random seed $42 + b$ for expression bin $b$. Within each bin, a deterministic subset-sum search selected complete clusters whose total size was closest to 20% of genes. The chromosome-blocked partition retained chromosomes 1, 14, 15 and X as the fixed held-out blocks. Repeated expression-stratified partitions used sampling seeds 123 and 456.

Each alternative split was evaluated in one paired seed-42 retraining with pretrained and random vectors in all four cohorts. The three-run primary analysis assessed optimization variability. We recorded held-out-set overlap and cosine similarity between each held-out vector and its nearest training-gene vector. Within the primary split, genes were also divided into cohort-specific quartiles of this similarity, and component effects were recalculated. This quartile analysis was post hoc; the cluster and chromosome partitions were retrained interventions. All partitions changed downstream gene assignments while leaving Decima pretraining unchanged; gene-family membership was unrestricted.

A historical prespecified split was retained as a secondary sensitivity analysis. It contains 14,717 training and 3,680 held-out brain genes, and 12,788 training and 3,126 held-out HER2ST genes. Its stratification used an earlier hippocampus working set that included three sections subsequently assigned to the final test set. It was therefore not fully independent of the final biological split. The historical and primary held-out brain sets shared 831 genes, with 2,849 unique to each (Jaccard index, 0.127). This comparison is reported in Supplementary Table S19 in the workbook and Source Data.

Pretrained Decima, random and constant conditions were independently trained for seeds 42, 123 and 456 in every cohort under both the primary and historical partitions. Architecture, individual splits, optimization, checkpoint selection and metrics were otherwise unchanged. This comparison changes gene membership and the resulting fitted parameters while retaining the rest of the evaluation design.

#### Ridge implementation and genomic covariates

Gene-mean-only ridge models selected $\alpha$ from $\{0.01, 0.1, 1, 10, 100, 1,000, 10,000\}$ by minimum validation root mean squared error (RMSE), breaking ties in favour of the smaller value. The models included an intercept and used the `lsqr` solver, tolerance $10^{-5}$ during selection and $10^{-6}$ for the final refit. Genomic-covariate ridge models used the same regularization grid and validation subset, standardizing numeric inputs and one-hot encoding categorical inputs within the fitted pipeline.

Gene-mean, full-matrix and centred endpoints were calculated analytically from the constant-across-spot predictions. We also correlated gene means from the correctly aligned ridge model with those from each complete pretrained-vector decoder.

For both Decima and scGPT, the fraction of full-matrix PCC gain retained without images used the complete decoder's random-vector PCC as a common baseline:

$$R_s = \frac{r_{\text{ridge,pretrained}} - r_{\text{decoder,random},s}}{r_{\text{decoder,pretrained},s} - r_{\text{decoder,random},s}}.$$

Here $r$ denotes full-matrix PCC on the same evaluated targets and spots within a representation and cohort, and $s$ indexes seeds 42, 123 and 456. The pretrained-vector ridge fit was fixed within each representation and cohort. Figure 3e and Extended Data Figs. 3c and 9b show $100 \text{ mean}_s(R_s)$. The MSE-retention calculation in Extended Data Fig. 3c used the same decoder-random baseline, with control-minus-prediction errors in both numerator and denominator. Ratios were calculated before averaging over seeds; this differs from a ratio of mean improvements. Denominators must be positive. The ratio is not bounded by 100% and does not measure a fraction of spatial information. Source Data provide the absolute inputs, per-run ratios and displayed summaries in `no_image_common_reference_per_run.tsv`, `no_image_common_reference_summary.tsv` and `no_image_displayed_summary.tsv`.

A second ridge model tested whether simple annotations reproduced mean-expression transfer. Predictors were log gene length, GC fraction, ambiguous-base fraction in the 524,288-bp reference window, chromosome, gene type and strand. Numeric features were standardized and categorical features one-hot encoded using training genes. Decima supervision-summary fields, including mean counts, track number and pretrained-model performance, were excluded. The target was the training-tissue gene mean. Absolute performance and adjusted gene-mean correlations are reported in Supplementary Table S18.

We also subtracted the covariate prediction from observed test gene means and pretrained-vector-predicted means before recalculating gene-mean PCC. This assesses ordering beyond the specified annotations. Isoform structure, mappability and capture efficiency were outside the covariate set.

#### Alternative mean transformation and detection restrictions

The primary mean is $\text{mean}(\log(\text{CPM} + 1))$. To assess the order of averaging and transformation, we reconstructed CPM as $\exp(Y_{ig}) - 1$, retained sparse zeros, accumulated values in double precision, averaged across spots and then applied $\log(1 + x)$. We retrained the no-image ridge on $\log[1 + \text{mean}(\text{CPM})]$ using training genes and individuals only. We also compared complete-decoder gene means with this alternative observed target in test individuals. Both definitions use spot-normalized CPM.

We retained held-out genes detected in at least 10% or 25% of test spots, present in every test section, or above the cohort-specific median detection fraction. Each subset was defined once from observed test expression and shared across pretrained and control conditions. Gene-mean PCC, mean within-gene PCC and exact MSE components were recalculated. These post-fitting restrictions did not affect training or checkpoint selection.

#### Target stratification and resampling

Held-out targets were ranked by expression standard deviation in training individuals. The 50 highest-ranked genes were fixed within each cohort and shared across conditions and seeds. Validation and test expression did not contribute to ranking. All selected genes met the observed-expression PCC eligibility criterion. Existing checkpoints were evaluated without retraining or reselection. Within-gene PCC was evaluated over pooled test spots and over pooled section-centred residuals. Centred RMSE was calculated as defined in the main Methods. The same targets were evaluated with identity-shuffled and target-fitted controls.

PCC effects were calculated within gene and seed, then averaged over paired runs. Cohort summaries included the median, interquartile range and fraction of positive effects, describing variation among the correlated targets.

Observed-eligible held-out genes were divided into ten cohort-specific deciles of training standard deviation, with assignments fixed across conditions and seeds. Effects were calculated within gene and seed, averaged across runs within gene, then summarized by decile means, medians, interquartile ranges and positive fractions. The primary Decima analysis used all eligible held-out genes; representation comparisons used the exact scGPT-covered set for both representations.

Biological uncertainty used section-centred pretrained-versus-random effects. Paired run effects were first averaged within section. For the individual-block bootstrap, section metrics were averaged within individual and test individuals were sampled with replacement 20,000 times. For the hierarchical bootstrap, individuals were sampled first and sections resampled within each selected occurrence; repeated occurrences of an individual received independent section resamples. We also recalculated cohort effects after omitting each individual in turn. Intervals use the 2.5th and 97.5th percentiles.

The 100 clusters from the embedding-cluster partition defined target-level resampling blocks. Complete held-out-gene clusters were sampled with replacement 20,000 times. Each draw’s pretrained-versus-random effect was calculated within seed and then averaged across seeds. Intervals use the 2.5th and 97.5th percentiles. Cluster resampling retains local dependence within Decima-vector clusters rather than treating genes as independent. Intervals are descriptive; no hypothesis tests were performed.

All biological-individual and target-cluster bootstrap summaries used 20,000 draws. Individual-block resampling used seed $23,081 + c$, hierarchical individual-then-section resampling used seed $91,103 + c$, and target-cluster resampling used seed $72,001 + c$, where $c$ was the zero-based cohort index in the fixed order hippocampus, DLPFC, NAc and HER2ST. The target-cluster bootstrap sampled the 100 complete clusters defined by the embedding-cluster construction.

#### Numerical and aggregation checks

Per-gene centred MSE was also verified as total MSE minus squared mean error; numerical negatives were truncated to zero. Section-centred metrics used a common observed-eligible gene set for matched conditions. DLPFC contributed one section per test donor, while the other cohorts could contribute several.

Two additional summaries were used. First, section-centred values were concatenated within gene across sections before calculation of PCC and its average over eligible genes (section-centred within-gene PCC). Constant concatenated predictions contributed zero. Second, squared errors and observation counts were pooled across all spot–gene pairs before calculating RMSE components. This observation-weighted summary complements equal weighting of individuals. All section-centred evaluations reused the corresponding fitted checkpoints without retraining or reselection.

#### Alternative decoders and pathway readouts

The architecture comparison used the primary split, frozen UNI2-h features, individual partitions, vector preprocessing and pretrained, dimension-matched random and constant controls. Each architecture and condition was independently optimized with seeds 42, 123 and 456 in every cohort. Complete-decoder results came from the primary analysis; bias-free, joint-MLP and residual-only models were separately trained.

##### Bias-free factorized decoder

This model retained the 512-unit encoders and 96-dimensional latent space but removed the gene-bias network:

$$\hat{Y}_{ig} = \text{softplus} \left( \frac{f_{\text{spot}}(\mathbf{x}_i)^\top f_{\text{gene}}(\mathbf{e}_g)}{\sqrt{96}} + b_0 \right). \quad (4)$$

The scalar $b_0$ was shared across genes and spots. Although explicit gene-specific biases are absent, the average spot representation can still support gene means. The model had 1,875,905 trainable parameters, compared with 2,371,777 in the complete decoder. Loss, six-epoch training and checkpoint selection were unchanged.

##### Joint concatenation MLP

The joint decoder applied a one-hidden-layer MLP to normalized spot and gene vectors. For memory efficiency, its first layer used the equivalent sum of separate projections:

$$\mathbf{q}_{ig} = \text{GELU} [W_x \text{LayerNorm}(\mathbf{x}_i) + W_e \text{LayerNorm}(\mathbf{e}_g) + a], \quad (5)$$

$$\hat{Y}_{ig} = \text{softplus} [\mathbf{w}^\top \text{dropout}(\mathbf{q}_{ig}) + c]. \quad (6)$$

Hidden width 704 gave 2,441,345 trainable parameters, 1.029 times the complete decoder. The model lacks an explicit gene-only skip branch but can learn a gene main effect by reducing dependence on the spot input. Objective, training, checkpoint selection and control-vector realizations matched the primary analysis.

#### Residual-only decoder

For section $s$, targets were $R_{isg} = Y_{isg} - \bar{Y}_{sg}$. A bias-free interaction with the same 512-unit encoders and 96-dimensional latent space produced signed outputs $Z_{isg}$ without softplus. Predictions were centred exactly within section and gene during training, validation and testing: $\hat{R}_{isg} = Z_{isg} - \bar{Z}_{sg}$.

The objective was residual MSE plus 0.15 times gene-wise correlation loss. Checkpoints maximized validation mean within-section gene PCC minus 0.02 times section-centred MSE. Section metrics were averaged within individual and then equally across individuals; MSE was aggregated before taking square roots. The model had 1,875,904 parameters. Raw-expression, full-matrix and gene-mean-calibration endpoints are not defined for this task.

#### Frozen-checkpoint readouts

We evaluated all 36 primary held-out-gene checkpoints—four cohorts, three seeds and three vector conditions—with complete, gene-bias-only and interaction-only readouts. Decima readouts were reused from the branch-intervention analysis; random and constant checkpoints used the same definitions. Checkpoint checksums were verified and complete readouts were checked against the primary results. The gene-bias-only readout was

$$\hat{Y}_{ig}^{\text{bias}} = \text{softplus}\{f_{\text{bias}}(\mathbf{e}_g)\}, \quad (7)$$

and the interaction-only readout was

$$\hat{Y}_{ig}^{\text{int}} = \text{softplus}\left\{\frac{f_{\text{spot}}(\mathbf{x}_i)^\top f_{\text{gene}}(\mathbf{e}_g)}{\sqrt{96}}\right\}. \quad (8)$$

The bias-only readout sets the interaction logit to zero and is constant across spots. It receives no tissue input at inference, although its parameters were learned jointly with the complete decoder. The interaction-only readout sets the bias logit to zero before softplus.

We intervened in the 12 pretrained-Decima checkpoints: four cohorts and three seeds. Held-out vectors were permuted using seed $s + 54,321$, preserving their multiset. Correct or permuted vectors were supplied independently to the bias network and gene encoder, giving four conditions: correct identity in both branches, only in the bias branch, only in the interaction branch, or in neither branch. Intact predictions reproduced the stored primary endpoints. Bias-only readouts were also evaluated and reused in the pathway analysis.

### Spatial-example selection

#### Fitted-gene examples

**Figure 2b,c** compares variant 4, the mean-informed graph-enabled direct head, with variant 8, the graph-enabled Decima-FiLM model. Predictions were averaged spotwise across seeds 42, 123 and 456 before section-level PCC was calculated.

Hippocampal *SLC17A7* was selected as a brain marker with a small negative pooled strategy effect, and HER2ST *A2M* as a tumour-cohort target with a modest positive effect. Candidate sections were examined in the fixed test-partition order. For hippocampus, we selected the first section with both PCCs between 0.20 and 0.70 and a FiLM-minus-direct difference between $-0.080$ and $-0.005$. For HER2ST, the corresponding intervals were 0.40–0.70 and 0.010–0.080.

Maps were standardized separately within gene, section and model and clipped to $[-2, 2]$ for display. Printed PCCs use unstandardized expression. The complete section screen and selection record accompany the source data.

### Held-out-gene example

The example in **Figure 3b,c** was selected using a recorded two-stage procedure. We first screened all 14,224 cohort-gene combinations across test spots and three runs. Eligibility required observed mean log-expression at least 0.75; pretrained-vector mean absolute error at most 25% of the observed mean and below both controls in every run; within-gene-PCC eligibility in all conditions and runs; observed spatial standard deviation at least 0.50; pretrained-vector PCC at least 0.10 in every run; and higher pretrained-vector PCC than both controls in every run. This retained 54 hippocampal, 56 DLPFC, 76 NAc and 58 HER2ST genes.

We then selected hippocampal *STX1B* post hoc from a recorded brain-marker shortlist, requiring both controls to predict means at least 0.25 in every run, and pretrained vectors to give lower gene-mean error and higher mean within-gene PCC across runs. *STX1B* also met all initial every-run criteria. After fixing gene and cohort, we chose the test section with observed *STX1B* mean nearest the median across sections, without reference to prediction performance. Maps use the recorded seed-42 checkpoints. The complete screen, display score and selection record accompany the source data.

### Neighbourhood-effect summaries and examples

Graph-enabled and zero-edge PCCs were calculated within gene and seed. Graph-minus-zero-edge effects were averaged across seeds before calculation of cohort medians and positive fractions. Density plots used an inverse-hyperbolic-sine horizontal transformation to show effects near zero while accommodating larger values; ticks show the original scale.

All test sections from an individual were combined. Mean within-gene PCC was calculated for each graph condition and seed, and paired effects were averaged across seeds within individual.

Candidates came from cohort-relevant marker lists. They had to rank among the 1,000 highest-variance training genes, show positive cohort-level graph effects in every seed and positive section-level effects in every test section, and achieve mean graph-enabled PCC at least 0.30. Eligible sections required zero-edge PCC at least 0.30, graph-enabled PCC at least 0.35, a PCC gain at least 0.04 and lower absolute error for at least 55% of spots. Genes were examined in a recorded interpretability order. For the first eligible marker, we displayed the section with the highest graph-enabled PCC. Maps were standardized separately for display; printed PCCs use unstandardized expression.

### External gene-query implementations

#### GeneQuery reconstruction

We reconstructed the public GeneQuery gene-aware pipeline at commit `7d61d38497316cce384fc11cdf7158cea752d413` [9]. Its official HER2 description-vector matrix had SHA-256 digest `61f482f4a5e4377f1085d6a95d28ff59cf721a9d4311e1abae9fc7813efbd83d`. The otherwise unpublished row order was recovered by filtering the original count-matrix header to the public 785-gene panel. The first 30 recovered symbols reproduced all 30 author-released descriptions, and recomputed Bio\_ClinicalBERT vectors numerically matched the first 30 official rows.

Of the 785 public-panel symbols, 34 lacked an unambiguous match in the Decima gene metadata. Intersecting the remaining genes with the fixed training-derived split retained 613 training and 138 held-out genes. Patients A–E were used for fitting, F for selection and G–H for testing. Frozen ImageNet-pretrained ResNet-50 features from the public `timm.create_model('resnet50')` call were projected to 256 dimensions, added to projected 768-dimensional description vectors and passed through the public two-layer gene transformer and scalar output head. The query head was fitted independently for seeds 42, 123 and 456 on $\log(\text{CPM} + 1)$. Semantic, dimension-matched random and constant conditions were separately fitted. Identity controls shuffled vectors separately within training and held-out genes before fitting. Evaluation genes and patients G–H were excluded from optimization and selection.

Because public GeneQuery launch scripts use a trainable image encoder, we also fitted end-to-end reconstructions with the full ResNet-50 and query head using seeds 42, 123 and 456 under the same exclusions. Semantic, identity-shuffled, random and constant-vector conditions were fitted independently within each seed; reported effects are means of paired differences across the three runs. This reconstruction retains the public gene-aware head but replaces the original random spot split with individual- and gene-disjoint testing.

Both GeneQuery settings used AdamW with learning rate $10^{-4}$, weight decay $10^{-3}$, training batch size 4 and evaluation batch size 8. Training ran for at most 100 epochs with automatic mixed precision on CUDA. The checkpoint minimizing validation MSE was retained, and early stopping used a patience of 12 epochs without improvement. Explicit run configurations are supplied with the code.

### DeepSpot-M checkpoint interventions

For DeepSpot-M, we used source commit `3ec046fffb9bed05974283a1a020ee01ebe6c22` and the gated released full checkpoint, with SHA-256 digest `7c57a60b82f3a32b54433430fccab4af2de400b89ca78c97d07cb63f0c6721b3` [10]. It contained 1,353,199,365 parameters and 19,338 gene entries. We selected the scGPT pathway and intersected the fixed HER2ST panel, retaining 601 fitted-panel and 135 held-out-panel genes. The same 3,097 spots from patients G–H received the released Midnight preprocessing. Each image batch passed through the backbone once. Stored patch tokens were decoded with correct gene-token rows, rows shuffled within panel partition, dimension-matched Gaussian rows or the fitted-panel mean token repeated across targets. The same intervened row was used by both query adapter and gene router.

These were inference-time interventions in one fixed checkpoint, evaluated on targets whose token rows had been trainable during upstream spatial training.

We evaluated the unmodified checkpoint outputs on the $\log(1 + \text{CP10K})$ scale specified for DeepSpot-M training [10]. Observed values were converted from the stored HER2ST log-CPM matrix as $Y_{ig}^{(10^4)} = \log[1 + (\exp(Y_{ig}^{(10^6)}) - 1)/100]$. The denominator was the per-spot count sum over the fixed 15,914-gene training-patient universe after zero filling, before restriction to the 135 targets. The reference was unsmoothed; we did not apply the original benchmark’s reference smoothing. The independent no-image ridge was fitted to training-patient gene means on the same CP10K scale, with the same fitted-gene penalty-selection procedure. All 135 targets were eligible for pooled and section-centred within-gene PCC; the corresponding count was 138 in both GeneQuery settings. Eligibility and constant-prediction conventions were unchanged.

No-image ridge fitting and component evaluation followed the main Methods, including the distinction between pooled-test and section-centred MSE decompositions.

### Donor-disjoint hippocampus external benchmark

#### Common design and evaluation

ST-Net [11], Hist2ST [12], BLEEP [13] and STPath [14] were retrained on the same hippocampal split used in the primary analysis: 22 sections from six training donors, eight sections from two validation donors and four sections from two test donors. The corresponding spot counts were 97,209, 32,044 and 14,406. All methods predicted the same ordered panel of 18,397 normalized $\log(\text{CPM} + 1)$ targets, where CPM denotes counts per million. Seeds 42, 123 and 456 controlled initialization, data order and method-specific stochastic operations. Prepared image inputs or image embeddings could be reused as immutable inputs, but no fitted checkpoint, adapter, partition assignment or metric from an earlier experiment was reused. The STPath image-feature adapter was refitted using training donors only.

Every method used the common validation donors and was evaluated only after checkpoint selection. Full test prediction matrices were aligned by section, spot barcode and gene identifier and passed to the common matrix- and gene-level evaluator. Checkpoint objectives and the adaptations needed to emit 18,397-gene predictions differed across architectures. The comparison evaluates these adapted implementations under a common biological split.

#### ST-Net

The ST-Net adaptation used a DenseNet-121 initialized with ImageNet weights and replaced its final layer with an 18,397-output regression head. It received 224-pixel spot-centred patches, used no gene filtering or expression transform beyond the locked analysis matrices and estimated channel normalization moments from training patches. Optimization used stochastic gradient descent for 10 epochs, batch size 4, learning rate $10^{-6}$, momentum 0.9 and zero weight decay. A validation prediction matrix was exported after every epoch, and the checkpoint minimizing validation sum of squared errors was selected independently for each seed; the selected epochs were 2, 1 and 1 for seeds 42, 123 and 456, respectively. Test inference averaged the eight rotations and reflections used by the implementation. The wrapper adapted the upstream data interface to the donor-disjoint split and complete output panel while retaining the model and image-regression procedure.

#### Hist2ST

Hist2ST was instantiated with 112-pixel spot-centred patches, six-nearest-neighbour adjacency, 64 positional bins, convolution kernel size 5, patch-embedding size 7, block depths 2, 8 and 4, 16 attention heads, 32 channels and dropout 0.2. To make complete-panel regression tractable, at most 1,024 spots were sampled deterministically per training or validation section; all test spots were used for final evaluation. The output layer predicted all 18,397 targets directly. Because the locked endpoint is continuous normalized log-expression, training used mean squared error (MSE) and disabled the zero-inflated negative-binomial, negative-binomial and self-distillation terms in the upstream implementation. Models were optimized for 10 epochs with Adam, learning rate $10^{-4}$ and weight decay $10^{-4}$. The checkpoint with minimum validation MSE was retained; selected epochs were 9, 8 and 6 for seeds 42, 123 and 456, respectively.

### BLEEP

The BLEEP code was taken from upstream repository revision `23959670c7407510c1716a173184bdcc18833a35`. A pretrained ResNet-50 image encoder and a complete-panel expression projection mapped 224-pixel patches and 18,397-gene spot vectors to 256-dimensional contrastive embeddings. Image patches were white padded at boundaries, normalized with ImageNet moments and augmented during training by horizontal and vertical flips and rotations in 90-degree increments. Models were trained for 10 epochs with AdamW, batch size 64, learning rate $10^{-4}$ and weight decay $10^{-4}$; the checkpoint minimizing validation contrastive loss was selected. Epoch 1 was selected for all three seeds. At test time, each image embedding retrieved the 50 most similar training-spot expression embeddings by cosine similarity. A softmax over those similarities supplied the weights used to average the corresponding complete 18,397-gene training-spot expression vectors. Thus, the full-panel adaptation enlarged the expression projection and retrieval output but retained BLEEP's contrastive and nearest-neighbour prediction structure.

### STPath

STPath used the official pretrained checkpoint with SHA-256 `03d49af98103c22eaae064632a366ad6ba2c1e627adc f47e00b13746a3b348fe`. Immutable 2,048-dimensional ImageNet-ResNet-50 spot embeddings were projected to the 1,536-dimensional image-input width expected by STPath using principal components fitted on training-donor spots only; the adapter artifact had SHA-256 `88406b00cfc72e4b0b1aa046c8e18b5dde1ca88280ccdd76c1482eea914bb4cc`. The technology and organ tokens were set to Visium and brain. The gene head and image-input projection were trained, the pretrained gene-token embedding was frozen and the final transformer block was unfrozen, yielding 22,890,580 trainable parameters. Models were optimized for eight epochs with AdamW, learning rate $10^{-4}$, weight decay $10^{-4}$ and gradient clipping at norm 1.0. Training steps sampled at most 512 genes and 2,048 spots; validation steps used at most 1,024 genes and 2,048 spots. The checkpoint maximizing validation full-matrix Pearson correlation coefficient (PCC) was retained, which was epoch 8 for every seed. Predictions were subsequently exported in spot and gene chunks over the complete aligned target panel.

### HER2ST tumour-program analysis

Pathology annotations were used only for a secondary analysis. In training sections A1, B1, C1, D1 and E1, tumour spots were invasive cancer or cancer in situ; non-tumour spots were adipose tissue, breast glands, connective tissue or immune infiltrate. Eligible genes had training mean above 0.05 and standard deviation above $10^{-3}$. For each gene, we calculated the within-section tumour-minus-non-tumour mean difference, divided by training standard deviation and averaged across the five sections. The 50 largest and 50 smallest contrasts defined positive and negative programme members.

Within each test section, each programme gene was standardized using training moments, multiplied by its programme direction and averaged across the 100 genes. The fixed programme was applied without reselection to annotated sections G2 and H1. Pathology discrimination used the area under the receiver-operating-characteristic curve; programme recovery used PCC between observed and predicted scores across spots.

### Computational reproducibility

Analyses ran on Linux through Slurm, with NVIDIA GPUs for neural-network training and inference. The primary codebase, ST-Net and BLEEP used Python 3.11 in `decima_env`; Hist2ST and STPath used Python 3.10 in `hist2st_env`. Historical records did not preserve a complete immutable package and GPU-runtime specification. The code release provides a version-pinned Linux CPU environment with CPython 3.10.19 and PyTorch 2.1.2 (`requirements-lock-linux-x86_64-py310.txt`). Fresh-environment checks covered installation, synthetic fitting and component evaluation, partition handling, and a bounded real-data integration run using the archived cohort inputs. These checks establish software execution on the tested inputs, not reproduction of the full original GPU analyses. Package inventories, commands, hashes and validation outputs are archived with the software release. Model revisions and file digests are specified under Representation extraction and provenance and External gene-query implementations above; third-party weights are obtained from their original providers.

### Supplementary Notes

#### Reporting conventions

Unless stated otherwise, plotted points show arithmetic means across optimization seeds 42, 123 and 456, and ranges connect the corresponding run values. These runs quantify computational variability. In comparisons with random or identity-shuffled gene vectors, they also vary the fixed control-vector assignment. Gene-level distributions describe variation among correlated targets. Biological-individual summaries identify the donor or patient as the sampling unit, and intervals from cohorts with only two test individuals are finite-sample sensitivity summaries. Correlation effects are candidate minus reference and error reductions are reference minus candidate, so positive values favour the candidate. The gene-mean share of a positive total MSE reduction can exceed 100% when centred MSE increases. No-image percentages use the common decoder-random baseline and are averaged over seed-paired ratios as defined above. No null-hypothesis tests were performed using the three optimization runs.

#### Supplementary Note 1: Fitted-gene calibration and neighbourhood effects

The fitted-gene experiment compared matched interventions (**Extended Data Fig. 1** and Supplementary Table S2 in the workbook). Mean-informed initialization preferentially changed full-matrix endpoints, whereas the six-nearest-neighbour aggregation plus Laplacian-smoothness bundle produced its clearest gains within genes. The effect was broad in hippocampus, DLPFC and NAc, where 64–80% of eligible genes improved and the median per-gene effect was positive. HER2ST was more heterogeneous: 45% of genes improved and the median effect was slightly negative, although the cohort-mean effect remained positive (**Extended Data Fig. 2a**). Effects remained positive after averaging optimization runs within held-out biological individuals (**Extended Data Fig. 2b**).

Registered hippocampal *SLC17A7* and DLPFC *PLP1* examples show how the same neighbourhood intervention can sharpen tissue-level gradients (**Extended Data Fig. 2c,d**). These maps are illustrative and were selected using the documented biology-guided rule in Supplementary Methods; printed correlations use unstandardized expression. The fitted-gene Decima-conditioned feature-wise linear modulation (FiLM) strategy remained cohort dependent, with its most consistent positive spatial effects in HER2ST. Because that contrast jointly changes target input, decoder parameterization, sharing, scale correction and anchoring, it is interpreted as an integrated strategy comparison.

A fusion-architecture sensitivity compared FiLM with direct concatenation and gene-to-spot cross-attention on the same graph-enabled background. Neither alternative consistently improved gene-wise spatial recovery across cohorts. FiLM gave the highest cohort-mean PCC among the 50 highest-variance genes in hippocampus, DLPFC and HER2ST and was effectively tied with concatenation in NAc; mean within-gene PCC showed the same overall pattern, with small reversals in HER2ST. Full-matrix PCC and overall MSE varied by cohort and did not favour one fusion operator uniformly. Supplementary Table S20 summarizes these comparisons; complete per-run values are provided in the Source Data.

#### Supplementary Note 2: Section-centred and gene-partition sensitivities

The primary component analysis centres each gene over all test spots in a cohort. Repeating the evaluation after removing observed and predicted means separately within each test section preserved the large gene-mean component and the much smaller within-section component (**Extended Data Fig. 4**; Supplementary Table S6). The ordering also persisted when centred spots were concatenated for PCC and when squared errors were pooled over spot–gene observations rather than averaged equally over biological individuals.

Four additional partitions constructed exclusively from final training individuals tested dependence on downstream target assignment and molecular proximity: two independently sampled expression-stratified repeats, an embedding-cluster-disjoint split and a chromosome-blocked split. After independent decoder fitting, all 16 cohort–partition combinations preserved large gene-mean and full-matrix gains, whereas within-gene effects remained much smaller (**Extended Data Fig. 7**; Supplementary Table S7). Each embedding cluster was assigned wholly to one side of the training–held-out boundary; nearest-training-gene Decima-vector similarities are provided in the Source Data. The historical working-set partition is reported separately in Supplementary Table S19; its construction and overlap with the primary partition are described in Supplementary Methods.

### Supplementary Note 3: Decoder pathway and architecture diagnostics

Frozen-checkpoint readouts from the primary training-only partition separated information available from the jointly optimized gene-bias and spot-gene interaction pathways (**Extended Data Fig. 8b–d**). The gene-bias-only output was spatially invariant within each gene and retained most matrix-level performance, whereas the interaction-only output retained more within-gene structure. Standard endpoint conventions were maintained: an observed-eligible gene with a constant within-gene readout contributed PCC zero, while a matrix- or gene-mean PCC with zero variance across its predicted vector was reported as undefined. Each readout set the other pathway's logit to zero before applying softplus. Decima readouts were reused from the branch-intervention analysis.

The mean-dominant profile did not require the explicit gene-conditioned bias. It persisted in a bias-free factorized decoder containing only one shared intercept and in a parameter-matched concatenation multilayer perceptron (MLP) without a gene-only skip branch (**Extended Data Fig. 8a**; Supplementary Tables S8 and S12). A residual-only decoder trained on exactly section-centred targets yielded modest, comparator-dependent spatial effects (Supplementary Table S9). Under the primary training-only partition, branch-specific gene-identity permutations preferentially damaged gene means when bias-path identity was broken and within-gene endpoints when interaction-path identity was broken; the frozen gene-bias-only readout retained most matrix-level performance (Supplementary Tables S10 and S11). Complete raw-expression architecture results and gene-bias retention are supplied in Supplementary Tables S8 and S11 in the workbook.

### Supplementary Note 4: Mean-only prediction, gene identity and biological sensitivity

An independently trained ridge mapped each frozen gene vector to one training-individual mean log-expression value and broadcast that value over all test spots (**Extended Data Fig. 3**). Matched random and identity-shuffled controls remained near zero, whereas the correctly aligned pretrained vectors reproduced most of the complete decoder's matrix gain. Using the complete decoder's random-vector performance as a common reference, the Decima no-image model retained 90.2–99.3% of the complete decoder's full-matrix PCC gain. Cross-cohort ridge transfer showed a partially shared ordering of gene means, but fitting the mapping within the target cohort remained advantageous.

Train-time identity shuffling preserved the vector multiset and covariance while breaking gene correspondence. Restoring correct identity recovered the large gene-mean signal, whereas transcriptome-wide within-gene effects remained much smaller (**Extended Data Fig. 6a,b**). The target-fitted reference supplied only a modest and cohort-dependent spatial increment over the held-out-gene decoder under the same spot representation and decoder family. Its top-quartile and top-decile signal strata use observed test variation and are therefore descriptive rather than prespecified target groups (**Extended Data Fig. 6c**).

Spatial effects became larger among targets with appreciable training-derived variation. On the fixed training-defined top-50 sets, effects remained positive after section-wise centring and relative to identity-shuffled vectors, although genes and cohorts were heterogeneous (**Extended Data Fig. 5**; Supplementary Tables S13 and S14). Simple genomic-covariate ridge models achieved gene-mean PCCs of 0.179–0.327, below the independently fitted Decima-vector ridge models (**Extended Data Fig. 3f**; Supplementary Table S18). Subtracting the covariate prediction from observed and Decima-predicted gene means preserved substantial agreement, indicating information beyond these specified annotations. Detection restrictions, an alternative gene-mean definition and individual- and target-cluster bootstrap summaries also preserved the mean-dominant ordering.

### Supplementary Note 5: scGPT checkpoint sensitivity and secondary analyses

The parallel representation analysis used fixed rows from the official scGPT whole-human gene-token table after checkpoint layer normalization; no cell-specific expression values or contextual transformer outputs were supplied. Vocabulary filtering was identical across pretrained and control conditions. Correct token identity reproduced the mean-dominant component profile, and an independent no-image ridge retained 81.8–90.4% of the complete decoder's full-matrix PCC gain, using the complete decoder's random-vector performance as the common baseline (**Extended Data Fig. 9a,b**; Supplementary Tables S15–S17). Gene-mean effects were positive in every held-out biological individual, whereas within-section effects were smaller and heterogeneous (**Extended Data Fig. 9c**). Among the 50 high-variance targets, descriptive same-gene Pearson correlations between Decima and scGPT section-centred effects were 0.51, 0.71, –0.14 and 0.87 in hippocampus, DLPFC, NAc and HER2ST, respectively. These correlations summarize related but non-identical spatial effects across correlated genes; the diagonal in **Fig. 5b** marks equal effects. **Extended Data Fig. 9d** shows variance-decile profiles on common targets.

The whole-human checkpoint is the primary cross-cohort scGPT analysis. The organ-specific brain checkpoint preserved the mean-dominant result in brain cohorts but produced a slightly negative HER2ST high-variance spatial effect, illustrating that the smaller spatial component depends on upstream representation and cohort context. Packaged scGPT metadata do not enumerate every upstream study, so related-tissue exposure or downstream-source overlap cannot be excluded.

#### **Donor-disjoint hippocampus benchmark**

This benchmark provides fitted-gene performance context under a common donor split (Supplementary Table S4). ST-Net, Hist2ST, BLEEP and STPath were adapted to the same 18,397-gene panel and compared with two fitted-gene variants from our experimental system. Hist2ST had the lowest overall MSE and highest full-matrix PCC. The mean-anchored Decima-FiLM and direct-head models had similar mean within-gene PCCs (0.106 and 0.105), while the direct-head model had the highest top-50 PCC. Both of the latter models included the neighbourhood bundle. The leading method therefore depended on the evaluation endpoint. Model-specific adaptations and checkpoint criteria limit the ranking's generality.

#### **HER2ST tumour-program analysis**

The HER2ST signed tumour programme was defined using training patients and evaluated without reselection in annotated sections from two test patients (Supplementary Table S5 in the workbook). This analysis evaluates programme-score recovery and pathology discrimination in the held-out patients.

### Supplementary Tables

The searchable workbook `Chen_Hicks_Supplementary_Tables.xlsx` contains one worksheet for every table with the displayed summary values. Full-precision per-run and per-gene records are distributed as Source Data. Unqualified mean within-gene PCC denotes the pooled-spot endpoint; within-section PCC averages section-level correlations within individuals, whereas section-centred within-gene PCC pools section-centred residuals before calculating correlations. This PDF includes Tables S1, S3, S4, S6, S7, S9, S10, S12–S18, S20 and S21. Tables S2 and S8 provide long-form results in the workbook; the secondary tumour-program analysis (S5), gene-bias retention summary (S11) and historical partition comparison (S19) are also retained there.

| Table | Content | Location |
| --- | --- | --- |
| S1 | Cohort design and biological-individual partitions | PDF and workbook |
| S2 | Fitted-gene absolute performance and matched effects | Workbook |
| S3 | Absolute held-out-gene performance | PDF and workbook |
| S4 | Donor-disjoint hippocampus benchmark | PDF and workbook |
| S5 | HER2ST tumour-program performance | Workbook |
| S6 | Section-wise centred held-out-gene effects | PDF and workbook |
| S7 | Training-only gene-partition sensitivity | PDF and workbook |
| S8 | Raw-expression decoder architecture comparison | Workbook |
| S9 | Residual-only decoder performance | PDF and workbook |
| S10 | Frozen-decoder branch-identity interventions | PDF and workbook |
| S11 | Frozen gene-bias retention summary | Workbook |
| S12 | Decoder architectures and parameter counts | PDF and workbook |
| S13 | Training-defined high-variance held-out genes | PDF and workbook |
| S14 | Matched controls for high-variance held-out genes | PDF and workbook |
| S15 | Gene-representation provenance and downstream coverage | PDF and workbook |
| S16 | Static scGPT gene-token replication | PDF and workbook |
| S17 | scGPT checkpoint sensitivity | PDF and workbook |
| S18 | Simple genomic-covariate controls for gene-mean prediction | PDF and workbook |
| S19 | Historical versus primary gene-partition comparison | Workbook |
| S20 | Fitted-gene fusion-architecture sensitivity | PDF and workbook |
| S21 | External-model performance across gene-vector conditions | PDF and workbook |

**Table S1.** Cohort design and biological-individual partitions. Individual and section counts are shown as total (train/validation/test). Aligned-panel sizes are shown for each cohort; no biological individual crossed partitions.

| Cohort | Tissue | Assay | Split unit | Individuals | Sections | Aligned genes |
| --- | --- | --- | --- | --- | --- | --- |
| Hippocampus | Anterior hippocampus | 10x Visium | Donor | 10 (6/2/2) | 34 (22/8/4) | 18,397 |
| DLPFC | Dorsolateral prefrontal cortex | 10x Visium | Donor | 63 (44/9/10) | 63 (44/9/10) | 18,397 |
| NAc | Nucleus accumbens | 10x Visium | Donor | 10 (6/2/2) | 38 (25/5/8) | 18,397 |
| HER2ST | HER2-positive breast cancer | Spatial Transcriptomics | Patient | 8 (5/1/2) | 36 (27/3/6) | 15,914 |

Hippocampus, DLPFC and NAc are distributed through GEO accessions GSE264692, GSE307403 and GSE307586, respectively. Their source processing used the 10x Genomics GRCh38 2020-A reference (GENCODE v32/Ensembl release 98); the HER2ST source study used GRCh38 with Ensembl release 86. The hippocampus analysis used 34 of the 36 Visium-H&E capture areas listed in the GEO sample manifest; Br2743 capture areas V11U08-081\_C1 and V11U08-081\_D1 were absent from the analysis input archive. HER2ST data are available at Zenodo (doi:10.5281/zenodo.4751624). DLPFC has one section per donor. HER2ST test data comprise patients G and H (six sections in total); only sections G2 and H1 have the pathology annotations used in Supplementary Table S5.

**Table S3.** Absolute held-out-gene performance under the primary training-only partition. Values are mean ± standard deviation across three analysis runs. The random condition redraws fixed standard-normal gene vectors in each run, so its variability combines vector realization and optimization and is not a biological confidence interval.

| Cohort | Gene input | Eligible genes | Overall MSE | Full-matrix PCC | Gene-centred full-matrix PCC | Pooled within-gene PCC | Gene-mean PCC |
| --- | --- | --- | --- | --- | --- | --- | --- |
| Hippocampus | Pretrained gene vectors | 3,482 | $2.061 \pm 0.002$ | $0.427 \pm 0.000$ | $0.164 \pm 0.002$ | $0.094 \pm 0.004$ | $0.803 \pm 0.001$ |
| Hippocampus | Random gene vectors | 3,482 | $2.547 \pm 0.001$ | $0.056 \pm 0.002$ | $0.095 \pm 0.004$ | $0.086 \pm 0.004$ | $0.011 \pm 0.013$ |
| Hippocampus | Constant gene vector | 3,482 | $2.488 \pm 0.004$ | $0.100 \pm 0.004$ | $0.119 \pm 0.004$ | $0.096 \pm 0.004$ | — |
| DLPFC | Pretrained gene vectors | 3,593 | $2.525 \pm 0.019$ | $0.453 \pm 0.004$ | $0.147 \pm 0.010$ | $0.081 \pm 0.005$ | $0.805 \pm 0.004$ |
| DLPFC | Random gene vectors | 3,593 | $3.250 \pm 0.008$ | $0.035 \pm 0.008$ | $0.087 \pm 0.003$ | $0.078 \pm 0.001$ | $-0.001 \pm 0.015$ |
| DLPFC | Constant gene vector | 3,593 | $3.136 \pm 0.003$ | $0.085 \pm 0.003$ | $0.103 \pm 0.003$ | $0.079 \pm 0.003$ | — |
| NAc | Pretrained gene vectors | 3,553 | $2.473 \pm 0.014$ | $0.417 \pm 0.003$ | $0.151 \pm 0.000$ | $0.075 \pm 0.001$ | $0.776 \pm 0.006$ |
| NAc | Random gene vectors | 3,553 | $2.970 \pm 0.004$ | $0.047 \pm 0.008$ | $0.085 \pm 0.002$ | $0.072 \pm 0.001$ | $0.008 \pm 0.017$ |
| NAc | Constant gene vector | 3,553 | $2.901 \pm 0.010$ | $0.085 \pm 0.002$ | $0.100 \pm 0.002$ | $0.074 \pm 0.002$ | — |
| HER2ST | Pretrained gene vectors | 3,027 | $2.306 \pm 0.029$ | $0.369 \pm 0.005$ | $0.152 \pm 0.001$ | $0.101 \pm 0.001$ | $0.739 \pm 0.009$ |
| HER2ST | Random gene vectors | 3,027 | $2.755 \pm 0.065$ | $0.049 \pm 0.008$ | $0.091 \pm 0.002$ | $0.091 \pm 0.001$ | $-0.002 \pm 0.016$ |
| HER2ST | Constant gene vector | 3,027 | $2.619 \pm 0.012$ | $0.095 \pm 0.001$ | $0.110 \pm 0.001$ | $0.096 \pm 0.001$ | — |

Brain-cohort evaluation uses 3,680 held-out genes; HER2ST uses 3,184. Gene-wise eligibility is defined only by observed standard deviation greater than $10^{-6}$, and an eligible gene with a constant predicted map contributes PCC zero. Thus, all gene-vector conditions use the same observed-defined denominator. Gene-mean PCC correlates observed and predicted gene means across targets; it is undefined for the constant-vector condition and shown as a dash. Gene-centred full-matrix PCC subtracts observed and predicted gene means before flattening the eligible residual matrices.

**Table S4.** Donor-disjoint hippocampus benchmark. Values are mean ± standard deviation across three analysis runs; standard deviations quantify optimization variability and are not confidence intervals.

| **a. Matrix-level endpoints** |  |  |
| --- | --- | --- |
| Model | Overall MSE | Full-matrix PCC |
| BLEEP | 1.940 ± 0.025 | 0.473 ± 0.009 |
| Hist2ST | 1.875 ± 0.015 | 0.496 ± 0.006 |
| ST-Net | 1.895 ± 0.000 | 0.487 ± 0.000 |
| STPath | 2.031 ± 0.001 | 0.432 ± 0.000 |
| Mean-anchored Decima-FiLM + 6-NN | 1.952 ± 0.055 | 0.488 ± 0.025 |
| Direct head + 6-NN | 2.142 ± 0.047 | 0.461 ± 0.005 |

| **b. Gene-wise endpoints** |  |  |  |  |
| --- | --- | --- | --- | --- |
| Model | Mean PCC | Median PCC | Top-50 PCC | Top-2,000 PCC |
| BLEEP | 0.055 ± 0.000 | 0.036 ± 0.001 | 0.263 ± 0.014 | 0.155 ± 0.004 |
| Hist2ST | 0.045 ± 0.019 | 0.034 ± 0.020 | 0.133 ± 0.045 | 0.123 ± 0.031 |
| ST-Net | 0.017 ± 0.009 | 0.012 ± 0.005 | 0.052 ± 0.022 | 0.036 ± 0.022 |
| STPath | 0.017 ± 0.000 | 0.011 ± 0.000 | 0.044 ± 0.002 | 0.038 ± 0.001 |
| Mean-anchored Decima-FiLM + 6-NN | 0.106 ± 0.011 | 0.085 ± 0.011 | 0.276 ± 0.009 | 0.235 ± 0.033 |
| Direct head + 6-NN | 0.105 ± 0.005 | 0.083 ± 0.003 | 0.306 ± 0.033 | 0.246 ± 0.017 |

Lower MSE is better; higher correlation is better. “Top” gene sets use a variance ranking defined from training individuals. Mean-anchored Decima-FiLM + 6-NN is fitted-gene variant 8, with the fixed training-tissue mean anchor, Decima-conditioned FiLM and the neighbourhood bundle. Direct head + 6-NN is variant 2, with the default direct per-gene head and the same neighbourhood bundle but without a gene vector or mean-informed initialization. Comparator implementations were adapted to the common full-panel output and donor split; the resulting ranking is endpoint-dependent. Matrix-level and gene-wise endpoints favoured different methods, with similar mean within-gene PCCs for the two neighbourhood-enabled models.

**Table S6.** Section-wise centred held-out-gene effects. Values are arithmetic means [minimum, maximum] across three paired analysis runs and are pretrained-vector minus control for correlations or control minus pretrained-vector for RMSE. Primary quantities first average sections within each biological individual and then average individuals equally.

| Cohort | Control | Gene-mean PCC | Gene-centred full-matrix PCC | Within-section gene PCC | Section-centred RMSE | Fraction of gene-mean MSE reduced | Fraction of centred MSE reduced | Gene-mean share (%) |
| --- | --- | --- | --- | --- | --- | --- | --- | --- |
| Hippocampus | random | 0.7845 [0.7721, 0.7953] | 0.0711 [0.0672, 0.0743] | 0.0094 [0.0067, 0.0129] | 0.0144 [0.0140, 0.0148] | 0.6392 [0.6375, 0.6414] | 0.0210 [0.0204, 0.0216] | 92.0 [91.8, 92.2] |
| Hippocampus | constant | NA | 0.0457 [0.0425, 0.0497] | -0.0017 [-0.0031, -0.0011] | 0.0110 [0.0104, 0.0119] | 0.6109 [0.6077, 0.6152] | 0.0161 [0.0152, 0.0175] | 93.0 [92.6, 93.4] |
| DLPFC | random | 0.7941 [0.7769, 0.8090] | 0.0494 [0.0473, 0.0518] | 0.0072 [0.0064, 0.0087] | 0.0058 [0.0044, 0.0066] | 0.6382 [0.6234, 0.6512] | 0.0079 [0.0061, 0.0089] | 97.7 [97.4, 98.2] |
| DLPFC | constant | NA | 0.0349 [0.0331, 0.0359] | 0.0034 [0.0025, 0.0039] | 0.0041 [0.0027, 0.0050] | 0.5985 [0.5811, 0.6102] | 0.0056 [0.0037, 0.0068] | 98.0 [97.7, 98.7] |
| NAc | random | 0.7605 [0.7379, 0.7729] | 0.0477 [0.0459, 0.0488] | 0.0028 [0.0024, 0.0034] | 0.0033 [0.0025, 0.0042] | 0.5597 [0.5432, 0.5809] | 0.0045 [0.0035, 0.0058] | 98.1 [97.5, 98.6] |
| NAc | constant | NA | 0.0355 [0.0325, 0.0378] | 0.0009 [-0.0013, 0.0024] | 0.0016 [0.0009, 0.0027] | 0.5243 [0.5132, 0.5444] | 0.0022 [0.0012, 0.0037] | 98.9 [98.2, 99.4] |
| HER2ST | random | 0.7148 [0.7045, 0.7259] | 0.0546 [0.0529, 0.0579] | 0.0089 [0.0082, 0.0095] | 0.0117 [0.0051, 0.0156] | 0.5427 [0.5322, 0.5584] | 0.0165 [0.0071, 0.0219] | 92.8 [90.4, 96.6] |
| HER2ST | constant | NA | 0.0357 [0.0338, 0.0374] | 0.0038 [0.0018, 0.0056] | 0.0034 [-0.0019, 0.0072] | 0.4633 [0.4430, 0.4857] | 0.0048 [-0.0027, 0.0102] | 97.1 [94.0, 101.8] |

All endpoints are calculated separately in each section, averaged within biological individual and then averaged equally across biological individuals. Gene-centred full-matrix PCC correlates the flattened residual matrices within each section. Constant-vector gene-mean PCC and its contrasts are undefined (NA). The fractions divide each component MSE reduction by that component's MSE under the matched control; for example, 0.6392 denotes a 63.92% reduction. Gene-mean share is the percentage of total primary MSE reduction attributable to reduced section-specific gene-mean error.

**Table S7.** Training-only gene-partition sensitivity corresponding to **Extended Data Fig. 7**. Correlation effects are Decima minus matched random vectors. The primary partition shows mean [minimum, maximum] across three paired optimization runs; each alternative partition shows its single paired seed-42 run.

| Cohort | Gene partition | Runs | Gene-mean<br>PCC gain | Full-matrix<br>PCC gain | Mean within-gene<br>PCC gain | Gene-mean share<br>of MSE reduction (%) |
| --- | --- | --- | --- | --- | --- | --- |
| Hippocampus | Primary expression-stratified | 3 | 0.7921 [0.7795, 0.8029] | 0.3713 [0.3685, 0.3730] | 0.0082 [0.0054, 0.0120] | 92.9 [92.7, 93.3] |
| Hippocampus | Expression-stratified repeat 123 | 1 | 0.7905 | 0.3687 | 0.0114 | 92.4 |
| Hippocampus | Expression-stratified repeat 456 | 1 | 0.8099 | 0.3798 | 0.0077 | 92.4 |
| Hippocampus | Embedding-cluster-disjoint | 1 | 0.7938 | 0.4523 | 0.0017 | 97.8 |
| Hippocampus | Chromosome-blocked | 1 | 0.8233 | 0.3637 | -0.0061 | 94.8 |
| DLPFC | Primary expression-stratified | 3 | 0.8055 [0.7870, 0.8207] | 0.4178 [0.4101, 0.4250] | 0.0038 [-0.0004, 0.0068] | 95.8 [95.1, 97.0] |
| DLPFC | Expression-stratified repeat 123 | 1 | 0.7909 | 0.4057 | 0.0051 | 95.1 |
| DLPFC | Expression-stratified repeat 456 | 1 | 0.7953 | 0.4160 | 0.0028 | 95.4 |
| DLPFC | Embedding-cluster-disjoint | 1 | 0.7545 | 0.4656 | 0.0038 | 97.2 |
| DLPFC | Chromosome-blocked | 1 | 0.8332 | 0.4343 | 0.0080 | 95.1 |
| NAc | Primary expression-stratified | 3 | 0.7683 [0.7448, 0.7810] | 0.3701 [0.3584, 0.3775] | 0.0032 [0.0032, 0.0033] | 95.5 [94.6, 96.1] |
| NAc | Expression-stratified repeat 123 | 1 | 0.7286 | 0.3513 | 0.0026 | 96.2 |
| NAc | Expression-stratified repeat 456 | 1 | 0.7751 | 0.3731 | 0.0026 | 95.2 |
| NAc | Embedding-cluster-disjoint | 1 | 0.7416 | 0.4237 | 0.0025 | 98.6 |
| NAc | Chromosome-blocked | 1 | 0.7642 | 0.3625 | 0.0017 | 95.1 |
| HER2ST | Primary expression-stratified | 3 | 0.7402 [0.7288, 0.7529] | 0.3196 [0.3156, 0.3233] | 0.0091 [0.0087, 0.0095] | 91.9 [89.0, 96.5] |
| HER2ST | Expression-stratified repeat 123 | 1 | 0.7238 | 0.3137 | 0.0082 | 89.5 |
| HER2ST | Expression-stratified repeat 456 | 1 | 0.7296 | 0.3142 | 0.0088 | 90.4 |
| HER2ST | Embedding-cluster-disjoint | 1 | 0.7430 | 0.3652 | 0.0101 | 93.5 |
| HER2ST | Chromosome-blocked | 1 | 0.6927 | 0.3030 | 0.0112 | 87.3 |

Partition construction used only final training individuals where expression statistics were required. Repeat labels 123 and 456 denote partition-sampling seeds, not optimization seeds. Gene-mean share is calculated within each run as gene-mean MSE reduction divided by total MSE reduction. The historical comparison, which was not fully independent of the final test split, is retained separately in Supplementary Table S19 in the workbook.

**Table S9.** Absolute performance of the residual-only factorized dot-product decoder. Targets and predictions were centred exactly within section. Values are arithmetic mean ± standard deviation across three independently trained analysis runs on the training-only gene partition.

| Cohort | Decoder | Gene vector | Mean within-section gene PCC | Section-centred RMSE |
| --- | --- | --- | --- | --- |
| Hippocampus | Residual-only factorized | Pretrained | $0.1121 \pm 0.0005$ | $1.3471 \pm 0.0011$ |
| | Residual-only factorized | Random | $0.0983 \pm 0.0073$ | $1.3627 \pm 0.0002$ |
| | Residual-only factorized | Constant | $0.1121 \pm 0.0008$ | $1.3597 \pm 0.0006$ |
| DLPFC | Residual-only factorized | Pretrained | $0.0599 \pm 0.0037$ | $1.4615 \pm 0.0010$ |
| | Residual-only factorized | Random | $0.0471 \pm 0.0031$ | $1.4689 \pm 0.0008$ |
| | Residual-only factorized | Constant | $0.0560 \pm 0.0003$ | $1.4658 \pm 0.0000$ |
| NAc | Residual-only factorized | Pretrained | $0.0659 \pm 0.0004$ | $1.4451 \pm 0.0022$ |
| | Residual-only factorized | Random | $0.0636 \pm 0.0014$ | $1.4535 \pm 0.0008$ |
| | Residual-only factorized | Constant | $0.0657 \pm 0.0005$ | $1.4492 \pm 0.0004$ |
| HER2ST | Residual-only factorized | Pretrained | $0.1145 \pm 0.0012$ | $1.4093 \pm 0.0046$ |
| | Residual-only factorized | Random | $0.1106 \pm 0.0013$ | $1.4183 \pm 0.0046$ |
| | Residual-only factorized | Constant | $0.1138 \pm 0.0005$ | $1.4089 \pm 0.0013$ |

**Table S10.** Frozen-decoder branch diagnostics under the primary training-only gene partition. Values are mean [minimum, maximum] across three runs. PCC losses and MSE increases are relative to the intact pretrained-vector decoder; positive values indicate worse performance after intervention.

| Branch-specific gene-identity permutations |  |  |  |  |  |  |
| --- | --- | --- | --- | --- | --- | --- |
| Cohort | Intervention | Full-matrix PCC loss | Overall MSE increase | Gene-mean PCC loss | Mean within-gene PCC loss | Within-section gene PCC loss |
| Hippocampus | Permute bias identity | 0.2959 [0.2715, 0.3285] | 0.558 [0.509, 0.643] | 0.5991 [0.5525, 0.6603] | 0.0009 [0.0009, 0.0010] | 0.0010 [0.0010, 0.0011] |
|  | Permute interaction identity | 0.0896 [0.0815, 0.1026] | 0.191 [0.173, 0.220] | 0.1508 [0.1348, 0.1762] | 0.0263 [0.0253, 0.0269] | 0.0295 [0.0277, 0.0307] |
| DLPFC | Permute bias identity | 0.3148 [0.2975, 0.3426] | 0.851 [0.794, 0.952] | 0.5835 [0.5539, 0.6316] | 0.0013 [0.0009, 0.0016] | 0.0011 [0.0008, 0.0014] |
|  | Permute interaction identity | 0.1019 [0.0978, 0.1076] | 0.311 [0.309, 0.312] | 0.1748 [0.1660, 0.1875] | 0.0289 [0.0266, 0.0310] | 0.0251 [0.0246, 0.0257] |
| NAc | Permute bias identity | 0.2805 [0.2719, 0.2904] | 0.654 [0.605, 0.697] | 0.5560 [0.5404, 0.5713] | 0.0011 [0.0010, 0.0013] | 0.0006 [0.0004, 0.0007] |
|  | Permute interaction identity | 0.1121 [0.1034, 0.1218] | 0.229 [0.208, 0.245] | 0.2078 [0.1897, 0.2258] | 0.0213 [0.0212, 0.0215] | 0.0198 [0.0196, 0.0200] |
| HER2ST | Permute bias identity | 0.2857 [0.2684, 0.2992] | 0.599 [0.527, 0.654] | 0.6413 [0.6033, 0.6658] | 0.0008 [0.0006, 0.0012] | 0.0011 [0.0008, 0.0016] |
|  | Permute interaction identity | 0.0242 [0.0188, 0.0299] | 0.024 [0.020, 0.030] | 0.0408 [0.0241, 0.0511] | 0.0092 [0.0077, 0.0111] | 0.0085 [0.0070, 0.0100] |

**Table S12.** Decoder architectures and parameter counts. Counts use the common 1,536-dimensional spot and 1,920-dimensional gene inputs.

| Decoder | Trainable parameters | Explicit bias | Prediction target |
| --- | --- | --- | --- |
| Full factorized | 2,371,777 | Gene-conditioned | Raw expression |
| Bias-free factorized | 1,875,905 | One shared scalar | Raw expression |
| Concatenation MLP | 2,441,345 | Shared output intercept; no gene-only skip branch | Raw expression |
| Residual-only factorized | 1,875,904 | None | Section-centred residual |

**Table S13.** Held-out-gene performance for training-defined top-50 high-variance targets. Within each cohort, held-out genes were ranked by pooled training-spot expression standard deviation using training biological individuals only. The selected genes were fixed across gene-vector conditions and analysis runs. Values are mean [minimum, maximum] across three runs.

| Cohort | Gene input | Genes | Pooled within-gene PCC | Section-centred within-gene PCC | Centred RMSE |
| --- | --- | --- | --- | --- | --- |
| Hippocampus | Pretrained gene vectors | 50 | 0.241 [0.236, 0.245] | 0.241 [0.238, 0.244] | 3.091 [3.089, 3.094] |
| Hippocampus | Random gene vectors | 50 | 0.174 [0.162, 0.182] | 0.175 [0.161, 0.185] | 3.160 [3.156, 3.163] |
| Hippocampus | Constant gene vector | 50 | 0.206 [0.199, 0.212] | 0.208 [0.201, 0.213] | 3.159 [3.155, 3.163] |
| DLPFC | Pretrained gene vectors | 50 | 0.254 [0.241, 0.261] | 0.208 [0.203, 0.210] | 2.868 [2.863, 2.875] |
| DLPFC | Random gene vectors | 50 | 0.223 [0.212, 0.229] | 0.168 [0.162, 0.175] | 2.931 [2.924, 2.935] |
| DLPFC | Constant gene vector | 50 | 0.233 [0.228, 0.236] | 0.182 [0.179, 0.185] | 2.924 [2.923, 2.926] |
| NAc | Pretrained gene vectors | 50 | 0.281 [0.280, 0.284] | 0.234 [0.232, 0.237] | 3.037 [3.034, 3.041] |
| NAc | Random gene vectors | 50 | 0.259 [0.253, 0.268] | 0.211 [0.202, 0.221] | 3.111 [3.105, 3.114] |
| NAc | Constant gene vector | 50 | 0.275 [0.267, 0.282] | 0.227 [0.217, 0.235] | 3.099 [3.096, 3.105] |
| HER2ST | Pretrained gene vectors | 50 | 0.234 [0.233, 0.235] | 0.212 [0.208, 0.215] | 2.890 [2.888, 2.892] |
| HER2ST | Random gene vectors | 50 | 0.205 [0.203, 0.207] | 0.188 [0.185, 0.191] | 2.930 [2.928, 2.934] |
| HER2ST | Constant gene vector | 50 | 0.215 [0.213, 0.217] | 0.199 [0.198, 0.201] | 2.927 [2.924, 2.929] |

Ranking and target selection used no validation- or test-individual expression. The pooled training-spot ranking can reflect within-section variation, differences among training sections or donors and measurement noise; these genes are therefore described as training-defined high-variance targets rather than as a formal set of spatially variable genes. Pooled within-gene PCC uses concatenated test spots without section centring. Section-centred within-gene PCC first subtracts each section's observed and predicted gene means, then concatenates residuals within gene before calculating PCC. Both columns average gene-wise PCCs over the same 50 targets. Within-section PCC, reported elsewhere, instead averages section-level correlations within individuals and then equally across individuals. All 50 genes satisfied the observed-expression eligibility rule. Centred RMSE removes each gene's mean over pooled test spots and is calculated on the $\log(\text{CPM} + 1)$ scale. Run ranges describe computational variation, not biological confidence intervals.

**Table S14.** Matched controls on the same training-defined top-50 high-variance targets. Mean within-gene PCC effects are candidate minus control. Values are mean [minimum, maximum] across three paired analysis runs. The final two columns summarize gene-level effects after averaging the three analysis runs within each gene.

| Cohort | Comparison | Mean PCC effect | Median gene effect | Genes improved |
| --- | --- | --- | --- | --- |
| Hippocampus | Pretrained minus random | 0.067 [0.059, 0.080] | 0.064 | 40/50 |
| Hippocampus | Pretrained minus constant | 0.035 [0.033, 0.037] | 0.035 | 36/50 |
| Hippocampus | Pretrained minus identity-shuffled | 0.051 [0.044, 0.064] | 0.056 | 39/50 |
| Hippocampus | Target-fitted reference minus held out | 0.012 [0.005, 0.017] | 0.007 | 33/50 |
| DLPFC | Pretrained minus random | 0.031 [0.028, 0.034] | 0.024 | 38/50 |
| DLPFC | Pretrained minus constant | 0.021 [0.012, 0.027] | 0.015 | 30/50 |
| DLPFC | Pretrained minus identity-shuffled | 0.029 [0.021, 0.036] | 0.017 | 39/50 |
| DLPFC | Target-fitted reference minus held out | 0.017 [0.016, 0.017] | 0.002 | 28/50 |
| NAc | Pretrained minus random | 0.022 [0.016, 0.027] | 0.019 | 43/50 |
| NAc | Pretrained minus constant | 0.006 [−0.002, 0.012] | 0.007 | 39/50 |
| NAc | Pretrained minus identity-shuffled | 0.012 [0.010, 0.015] | 0.010 | 43/50 |
| NAc | Target-fitted reference minus held out | 0.011 [0.001, 0.020] | 0.001 | 30/50 |
| HER2ST | Pretrained minus random | 0.029 [0.027, 0.030] | 0.017 | 40/50 |
| HER2ST | Pretrained minus constant | 0.019 [0.018, 0.020] | 0.009 | 37/50 |
| HER2ST | Pretrained minus identity-shuffled | 0.022 [0.020, 0.024] | 0.014 | 39/50 |
| HER2ST | Target-fitted reference minus held out | −0.005 [−0.008, −0.001] | −0.004 | 16/50 |

Identity-shuffled vectors preserve the standardized pretrained-vector multiset but break gene–vector correspondence during downstream fitting. The target-fitted reference uses the same frozen spot representation and decoder family as the held-out-gene decoder but allows the evaluation targets to participate in fitting and checkpoint selection. Gene-level summaries are descriptive and do not treat the 50 correlated genes as independent biological replicates.

**Table S15. Gene-representation provenance and downstream coverage.** Coverage is calculated within the primary downstream target partition. The Decima and scGPT analyses use different vector dimensions and downstream training-gene universes; their performance is therefore compared through within-representation contrasts rather than as a leaderboard.

| Representation | Attribute | Details |
| --- | --- | --- |
| Decima sequence-derived | Upstream modality | Reference sequence with transcriptomic supervision |
|  | Frozen checkpoint | Decima v0.5.1 **v1_rep0**; checkpoint digest **9b4efc2967d0...** |
|  | Vector extraction | Mean over the frozen final feature-map sequence axis for a strand-aware 524,288-bp hg38 window; one vector per gene and no individual variants |
|  | Dimension | 1,920 |
|  | Gene annotation sources | 10x Genomics GRCh38 2020-A, supplemented with 2024-A and NCBI Gene mappings; released metadata used without reannotation |
|  | Downstream held-out coverage | 3,680 brain targets and 3,184 HER2ST targets in the complete primary analysis |
|  | Upstream gene partitions | Brain held-out targets: 2,895 train (78.7%), 425 validation (11.5%) and 360 test (9.8%); HER2ST: 2,517 train (79.1%), 365 validation (11.5%) and 302 test (9.5%) in Decima’s packaged gene metadata |
|  | Recorded study identifiers | No matches to the queried downstream identifiers in packaged supervision metadata; related-tissue profiles were recorded separately |
| scGPT static gene-token | Upstream modality | Single-cell transcriptomic pretraining; official whole-human checkpoint |
|  | Frozen checkpoint | **whole_human_model/best_model.pt**; checkpoint digest **6cb5d451ab5c...**; vocabulary digest **acca93d114ca...** |
|  | Vector extraction | **encoder.embedding.weight** token row followed by checkpoint **encoder.enc_norm**; no cell-specific values or contextual transformer output |
|  | Dimension | 512 |
|  | Downstream held-out coverage | 3,574 of 3,680 brain targets (97.1%) and 3,179 of 3,184 HER2ST targets (99.8%); all fixed top-50 targets covered |

The scGPT model-zoo metadata describe pretraining on 33 million normal human cells, but the packaged checkpoint does not enumerate every upstream dataset; related-tissue exposure or overlap with downstream source studies cannot be excluded. Full checkpoint, vocabulary and argument-file SHA-256 digests are reported in Supplementary Methods and the source-data manifest; vector extraction used local scGPT repository commit **cebd6fae655b...**. Out-of-vocabulary genes had lower training expression and detection and were enriched for non-protein-coding categories; the numerical audit and complete gene list are provided in Source Data files **representation\_scgpt\_covered\_vs\_oov.tsv** and **representation\_scgpt\_oov\_genes.tsv.gz**. Exact common-target analyses therefore use 3,574 held-out genes in each brain cohort and 3,179 in HER2ST. “Held out” denotes exclusion from downstream decoder fitting and selection, not necessarily from representation pretraining. The organ-specific brain checkpoint was evaluated separately as a checkpoint-sensitivity analysis.

**Table S16. Static scGPT gene-token replication and identity control.** Values are mean [minimum, maximum] across three paired analysis runs. All effects are calculated within the scGPT analysis on the same vocabulary-covered targets. Correlation effects are candidate minus control; MSE reduction is control minus candidate. The no-image ridge maps gene vectors to gene means estimated from training individuals and broadcasts the predictions over tissue locations.

| a. Pretrained minus matched random vectors |  |  |  |  |  |
| --- | --- | --- | --- | --- | --- |
| Cohort | Targets | $\Delta$ full-matrix PCC | $\Delta$ gene-mean PCC | $\Delta$ mean within-gene PCC | Gene-mean share of MSE reduction |
| Hippocampus | 3,574 | 0.444 [0.443, 0.445] | 0.935 [0.928, 0.950] | 0.0161 [0.0122, 0.0230] | 91.5 [91.0, 92.2]% |
| DLPFC | 3,574 | 0.494 [0.493, 0.496] | 0.947 [0.944, 0.950] | 0.0101 [0.0094, 0.0107] | 95.3 [95.3, 95.3]% |
| NAc | 3,574 | 0.446 [0.445, 0.447] | 0.909 [0.909, 0.911] | 0.0065 [0.0057, 0.0071] | 92.6 [92.4, 92.7]% |
| HER2ST | 3,179 | 0.346 [0.340, 0.354] | 0.805 [0.791, 0.821] | 0.0006 [0.0000, 0.0014] | 94.3 [93.8, 94.9]% |

| b. High-variance and no-image summaries |  |  |  |
| --- | --- | --- | --- |
| Cohort | Top-50 section-centred within-gene ΔPCC | No-image PCC improvement over decoder-random baseline | PCC gain retained without images (%) |
| Hippocampus | 0.095 [0.086, 0.110] | 0.379 [0.375, 0.381] | 85.3 [84.7, 85.7] |
| DLPFC | 0.068 [0.063, 0.080] | 0.447 [0.444, 0.449] | 90.4 [90.2, 90.7] |
| NAc | 0.040 [0.039, 0.043] | 0.385 [0.385, 0.386] | 86.4 [86.3, 86.5] |
| HER2ST | 0.012 [0.010, 0.013] | 0.283 [0.281, 0.284] | 81.8 [80.0, 83.3] |

| c. Pretrained minus train-time identity shuffle |  |  |  |  |  |
| --- | --- | --- | --- | --- | --- |
| Cohort | Targets | $\Delta$ full-matrix PCC | $\Delta$ gene-mean PCC | $\Delta$ mean within-gene PCC | Gene-mean share of MSE reduction |
| Hippocampus | 3,574 | 0.454 [0.451, 0.458] | 0.943 [0.935, 0.950] | 0.0168 [0.0146, 0.0211] | 91.4 [91.1, 92.0]% |
| DLPFC | 3,574 | 0.503 [0.495, 0.516] | 0.956 [0.944, 0.977] | 0.0108 [0.0089, 0.0122] | 95.1 [94.9, 95.3]% |
| NAc | 3,574 | 0.464 [0.457, 0.473] | 0.944 [0.930, 0.960] | 0.0059 [0.0052, 0.0063] | 92.4 [92.3, 92.5]% |
| HER2ST | 3,179 | 0.347 [0.338, 0.353] | 0.808 [0.789, 0.828] | 0.0009 [-0.0002, 0.0018] | 94.2 [93.5, 94.6]% |

Section-centred within-gene PCC subtracts observed and predicted gene means separately within each test section, concatenates residuals within gene, and averages the resulting gene-wise PCCs. Unqualified mean within-gene PCC uses pooled test spots without section centring. No-image PCC improvements and retained fractions in panel b use the complete decoder's random-vector PCC as the common baseline for both models. Percentages are calculated within each seed before averaging; they are not ratios of the displayed mean improvements. The constant-vector condition is omitted from gene-mean PCC contrasts because it predicts the same gene mean for every target, giving zero across-gene variance and an undefined correlation. Analysis runs are not biological replicates. The top-50 targets were fixed from pooled variance in downstream training individuals before test evaluation.

**Table S17. Sensitivity to the scGPT pretraining checkpoint.** Values are mean [minimum, maximum] across three paired analysis runs for correct token vectors relative to dimension-matched random vectors. The whole-human checkpoint is the primary checkpoint used across the four cohort-specific analyses; the brain checkpoint is retained as a checkpoint-sensitivity analysis. Comparisons are descriptive because the checkpoints have different upstream training domains.

| Checkpoint | Cohort | $\Delta$ full-matrix PCC | $\Delta$ gene-mean PCC | $\Delta$ mean within-gene PCC | Top-50 section-centred within-gene ΔPCC |
| --- | --- | --- | --- | --- | --- |
| Whole-human | Hippocampus | 0.444 [0.443, 0.445] | 0.935 [0.928, 0.950] | 0.0161 [0.0122, 0.0230] | 0.095 [0.086, 0.110] |
|  | DLPFC | 0.494 [0.493, 0.496] | 0.947 [0.944, 0.950] | 0.0101 [0.0094, 0.0107] | 0.068 [0.063, 0.080] |
|  | NAc | 0.446 [0.445, 0.447] | 0.909 [0.909, 0.911] | 0.0065 [0.0057, 0.0071] | 0.040 [0.039, 0.043] |
|  | HER2ST | 0.346 [0.340, 0.354] | 0.805 [0.791, 0.821] | 0.0006 [0.0000, 0.0014] | 0.012 [0.010, 0.013] |
| Brain | Hippocampus | 0.441 [0.439, 0.442] | 0.929 [0.921, 0.944] | 0.0157 [0.0116, 0.0227] | 0.099 [0.089, 0.114] |
|  | DLPFC | 0.493 [0.490, 0.496] | 0.944 [0.940, 0.948] | 0.0107 [0.0093, 0.0131] | 0.066 [0.060, 0.076] |
|  | NAc | 0.442 [0.441, 0.443] | 0.901 [0.901, 0.902] | 0.0068 [0.0055, 0.0077] | 0.040 [0.037, 0.043] |
|  | HER2ST | 0.276 [0.274, 0.279] | 0.655 [0.645, 0.662] | -0.0025 [-0.0036, -0.0019] | -0.004 [-0.008, -0.001] |

Section-centred within-gene PCC subtracts observed and predicted gene means separately within each test section, concatenates residuals within gene, and averages the resulting gene-wise PCCs. Unqualified mean within-gene PCC uses pooled test spots without section centring. The gene vocabulary and covered downstream targets are identical for these two packaged checkpoints. The primary conclusion—large gene-mean and full-matrix gains with much smaller transcriptome-wide within-gene gains—is shared. The HER2ST top-50 section-centred effect changes from slightly negative with the brain checkpoint to positive with the whole-human checkpoint, illustrating checkpoint sensitivity of the smaller spatial component.

**Table S18.** Simple genomic-covariate controls for Decima gene-mean prediction on the primary held-out targets. Panel a compares two independently fitted ridge models; panel b reports gene-mean correlations before and after subtracting the covariate prediction from both observed and Decima-predicted test gene means.

| **a. Independent no-image ridge models** |  |  |  |  |
| --- | --- | --- | --- | --- |
| Cohort | Covariates<br>gene-mean PCC | Decima<br>gene-mean PCC | Covariates<br>full-matrix PCC | Decima<br>full-matrix PCC |
| Hippocampus | 0.1795 | 0.8221 | 0.0901 | 0.4126 |
| DLPFC | 0.2009 | 0.8306 | 0.1088 | 0.4497 |
| NAc | 0.1791 | 0.8023 | 0.0916 | 0.4105 |
| HER2ST | 0.3267 | 0.7194 | 0.1534 | 0.3378 |

| **b. Gene-mean agreement after subtracting covariate predictions** |  |  |  |  |
| --- | --- | --- | --- | --- |
| Cohort | Complete decoder<br>unadjusted PCC | Complete decoder<br>adjusted PCC | No-image Decima ridge<br>adjusted PCC |  |
| Hippocampus | 0.8035 [0.8024, 0.8052] | 0.7964 [0.7954, 0.7981] |  | 0.8152 |
| DLPFC | 0.8048 [0.8012, 0.8097] | 0.7959 [0.7920, 0.8011] |  | 0.8225 |
| NAc | 0.7759 [0.7711, 0.7825] | 0.7672 [0.7623, 0.7734] |  | 0.7948 |
| HER2ST | 0.7385 [0.7283, 0.7459] | 0.7070 [0.6962, 0.7142] |  | 0.6806 |

Covariates were log gene length, GC and ambiguous-base fractions in the reference sequence window, chromosome, gene type and strand. Both ridge models were fitted on training genes and training-individual means and selected using the same validation subset. The pretrained-vector ridge and covariate ridge each give one deterministic result; complete-decoder summaries are mean [minimum, maximum] across three runs. Full-precision values are in `supp_stage2_simple_genomic_covariates.tsv` and `supp_stage2_covariate_residualized_abundance.tsv`.

**Table S20.** Fitted-gene fusion-architecture sensitivity. FiLM, direct concatenation and gene-to-spot cross-attention used the same graph-enabled background. Values are mean ± standard deviation across optimization seeds 42, 123 and 456.

| Cohort | Fusion | Mean within-gene<br>PCC | Top-50 PCC | Full-matrix<br>PCC | Overall MSE |
| --- | --- | --- | --- | --- | --- |
| Hippocampus | FiLM | 0.1112 ± 0.0034 | 0.2745 ± 0.0078 | 0.3997 ± 0.0166 | 2.5927 ± 0.5057 |
| Hippocampus | Concatenation | 0.1104 ± 0.0014 | 0.2701 ± 0.0031 | 0.3828 ± 0.0100 | 2.9314 ± 0.5376 |
| Hippocampus | Cross-attention | 0.1043 ± 0.0018 | 0.2486 ± 0.0193 | 0.3999 ± 0.0304 | 2.2403 ± 0.1561 |
| DLPFC | FiLM | 0.0828 ± 0.0068 | 0.2365 ± 0.0087 | 0.4065 ± 0.0243 | 3.3019 ± 0.5821 |
| DLPFC | Concatenation | 0.0631 ± 0.0025 | 0.1601 ± 0.0198 | 0.4042 ± 0.0089 | 3.0428 ± 0.2302 |
| DLPFC | Cross-attention | 0.0701 ± 0.0154 | 0.1884 ± 0.0614 | 0.4097 ± 0.0108 | 2.9729 ± 0.3256 |
| NAc | FiLM | 0.0768 ± 0.0062 | 0.1881 ± 0.0196 | 0.3959 ± 0.0181 | 3.3532 ± 0.7817 |
| NAc | Concatenation | 0.0765 ± 0.0051 | 0.1881 ± 0.0169 | 0.3804 ± 0.0398 | 2.6658 ± 0.2724 |
| NAc | Cross-attention | 0.0733 ± 0.0026 | 0.1695 ± 0.0097 | 0.3804 ± 0.0064 | 2.5448 ± 0.0339 |
| HER2ST | FiLM | 0.0982 ± 0.0041 | 0.1749 ± 0.0092 | 0.3504 ± 0.0094 | 2.3379 ± 0.0179 |
| HER2ST | Concatenation | 0.1016 ± 0.0078 | 0.1545 ± 0.0125 | 0.3491 ± 0.0085 | 2.4160 ± 0.1042 |
| HER2ST | Cross-attention | 0.0993 ± 0.0136 | 0.1519 ± 0.0155 | 0.3358 ± 0.0182 | 2.5359 ± 0.2256 |

All three fusion strategies used a learned Decima-derived mean anchor, the graph-enabled backbone and a learned scale correction. The FiLM row corresponds to fitted-gene variant 7; Figure 2a instead compares variant 8, which uses a fixed training-tissue mean anchor, with variant 4. These fitted-gene comparisons assess fusion strategies, not transfer to genes excluded from fitting. Top-50 targets follow the fitted-gene evaluation definition in Methods. Run-to-run standard deviations describe computational variation. Full-precision per-run values are supplied in Source Data as `supp_table_20_fusion_per_run.tsv`.

**Table S21.** External-model performance across gene-vector conditions. GeneQuery values are means (s.d.) over three independently fitted runs; DeepSpot-M values describe one fixed checkpoint. All settings use HER2ST patients G and H (six sections, 3,097 spots).

| a. Squared errors |  |  |  |  |
| --- | --- | --- | --- | --- |
| Setting | Vectors | Overall MSE | Gene-mean MSE | Centred MSE |
| GeneQuery, frozen | Correct | 6.6101 (0.2718) | 2.0568 (0.2719) | 4.5533 (0.0110) |
| GeneQuery, frozen | Shuffled | 7.6417 (0.1180) | 3.0011 (0.1991) | 4.6406 (0.0878) |
| GeneQuery, frozen | Random | 5.9594 (0.1228) | 1.5165 (0.1121) | 4.4429 (0.0108) |
| GeneQuery, frozen | Constant | 6.0327 (0.1084) | 1.5331 (0.0953) | 4.4997 (0.0150) |
| GeneQuery, end-to-end | Correct | 6.5021 (0.3245) | 1.9018 (0.3066) | 4.6003 (0.0828) |
| GeneQuery, end-to-end | Shuffled | 6.1173 (0.1027) | 1.6305 (0.1331) | 4.4867 (0.0624) |
| GeneQuery, end-to-end | Random | 6.3694 (0.1607) | 1.8965 (0.1501) | 4.4728 (0.0137) |
| GeneQuery, end-to-end | Constant | 6.1858 (0.2133) | 1.6674 (0.2584) | 4.5184 (0.0451) |
| DeepSpot-M | Correct | 0.5394 | 0.0307 | 0.5087 |
| DeepSpot-M | Shuffled | 0.9228 | 0.4015 | 0.5213 |
| DeepSpot-M | Random | 0.7757 | 0.2700 | 0.5058 |
| DeepSpot-M | Constant | 0.7216 | 0.2183 | 0.5033 |

| b. Correlation endpoints |  |  |  |  |  |  |  |
| --- | --- | --- | --- | --- | --- | --- | --- |
| Setting | Vectors | Full-matrix PCC | Gene-mean PCC | Section-centred full-matrix PCC | Pooled within-gene PCC | Section-centred within-gene PCC | Within-section PCC |
| GeneQuery, frozen | Correct | 0.1573 (0.0025) | 0.2627 (0.0065) | 0.1070 (0.0011) | 0.1132 (0.0022) | 0.1086 (0.0025) | 0.1078 (0.0022) |
| GeneQuery, frozen | Shuffled | -0.0089 (0.0181) | -0.1107 (0.0308) | 0.0911 (0.0055) | 0.1029 (0.0062) | 0.1006 (0.0068) | 0.1003 (0.0066) |
| GeneQuery, frozen | Random | 0.0974 (0.0335) | 0.0044 (0.1554) | 0.1230 (0.0029) | 0.1225 (0.0026) | 0.1204 (0.0025) | 0.1204 (0.0025) |
| GeneQuery, frozen | Constant | 0.0999 (0.0059) | NA | 0.1143 (0.0071) | 0.1143 (0.0061) | 0.1120 (0.0064) | 0.1113 (0.0061) |
| GeneQuery, end-to-end | Correct | 0.1553 (0.0213) | 0.2677 (0.0499) | 0.0985 (0.0023) | 0.1013 (0.0049) | 0.0968 (0.0055) | 0.0956 (0.0054) |
| GeneQuery, end-to-end | Shuffled | 0.0577 (0.0218) | -0.0839 (0.0481) | 0.1067 (0.0061) | 0.1068 (0.0055) | 0.1051 (0.0040) | 0.1043 (0.0039) |
| GeneQuery, end-to-end | Random | 0.0698 (0.0383) | 0.0063 (0.1459) | 0.0883 (0.0100) | 0.0862 (0.0115) | 0.0846 (0.0108) | 0.0837 (0.0107) |
| GeneQuery, end-to-end | Constant | 0.0875 (0.0073) | NA | 0.0993 (0.0085) | 0.0991 (0.0098) | 0.0969 (0.0103) | 0.0962 (0.0103) |
| DeepSpot-M | Correct | 0.4978 | 0.9280 | 0.0749 | 0.0764 | 0.0720 | 0.0719 |
| DeepSpot-M | Shuffled | 0.0170 | 0.0258 | 0.0218 | 0.0415 | 0.0433 | 0.0431 |
| DeepSpot-M | Random | -0.0059 | -0.0404 | 0.0388 | 0.0475 | 0.0492 | 0.0483 |
| DeepSpot-M | Constant | 0.0417 | NA | 0.0522 | 0.0585 | 0.0575 | 0.0571 |

GeneQuery uses $\log(1 + \text{CPM})$; DeepSpot-M uses $\log(1 + \text{CP10K})$, matching its documented training scale. MSE components use pooled test gene means. Correlations use unsmoothed expression. All 138 GeneQuery targets and all 135 DeepSpot-M targets satisfy the observed-expression eligibility rule for pooled and section-centred within-gene PCC in every condition. Within-section PCC averages correlations over eligible genes within section, then sections within patient and patients equally. GeneQuery controls are independently fitted; DeepSpot-M vectors are changed only at inference. Target coverage, expression scale and training history differ across implementations. NA denotes undefined constant-vector gene-mean PCC. Standard deviations describe computational variation, not biological uncertainty. Full-precision run, gene, section and patient records are provided in Source Data.
